# Deep Divergences Among Inconspicuously Colored Clades of *Epipedobates* Poison Frogs

**DOI:** 10.1101/2023.06.29.547117

**Authors:** Karem López-Hervas, Juan C. Santos, Santiago R. Ron, Mileidy Betancourth-Cundar, David C. Cannatella, Rebecca D. Tarvin

**Author notes:** Current Address: Department of Evolutionary Genetics, Max Planck Institute for Evolutionary Biology, Plön, Germany 24306 (Karem López-Hervas).

## Abstract

Poison frogs (Dendrobatidae) are famous for their aposematic species, having a combination of diverse color patterns and defensive skin toxins, yet most species in this family are inconspicuously colored and considered non-aposematic. *Epipedobates* is among the youngest genus-level clades of Dendrobatidae that includes both aposematic and inconspicuous species. Using Sanger-sequenced mitochondrial and nuclear markers, we demonstrate deep genetic divergences among inconspicuous species of *Epipedobates* but relatively shallow genetic divergences among conspicuous species. Our phylogenetic analysis includes broad geographic sampling of the inconspicuous lineages typically identified as *E. boulengeri* and *E. espinosai*, which reveals two putative new species, one in west-central Colombia (*E.* sp. 1) and the other in north-central Ecuador (*E.* aff. *espinosai*). We conclude that *E. darwinwallacei* is a junior subjective synonym of *E. espinosai*. We also clarify the geographic distributions of inconspicuous *Epipedobates* species including the widespread *E. boulengeri.* We provide a qualitative assessment of the phenotypic diversity in each nominal species, with a focus on the color and pattern of inconspicuous species. We conclude that *Epipedobates* contains eight known valid species, six of which are inconspicuous. A relaxed molecular clock analysis suggests that the most recent common ancestor of *Epipedobates* is ∼11.1 million years old, which nearly doubles previous estimates. Last, genetic information points to a center of species diversity in the Chocó at the southwestern border of Colombia with Ecuador. A Spanish translation this text is available in the supplementary materials on the publisher’s website (biorxiv does not allow posting in non-English languages).

## 1. Introduction

One of the challenges to taxonomists is the application of the bewildering array of species concepts (what a species is) to the practice of species delimitation (assigning individuals to species). de Queiroz (2005, 2007) attempted a unification of species concepts, proposing that a species is “a separately evolving lineage segment” or “a separately evolving metapopulation lineage,” suggesting that the only necessary property of species is its existence as a population-level lineage. Thus, under his unified species concept a lineage need not be phenotypically distinguishable, reproductively isolated, or ecologically divergent to be considered a species. In other words, the absence of one or more of these attributes does not mean that the lineage is not diverging, but rather is simply evidence that the lineage has not yet evolved those attributes. The attributes provide evidence for distinct lineages, and more lines of evidence provide a greater degree of support for species status.

Most descriptions of new species are based primarily on phenotypic differences (morphospecies; [Cain, 1954]) in the absence of information on genetic divergence. Ernst Mayr, who sought to elevate systematics and speciation to a central role in the evolutionary process through his biological species concept (Mayr, 1942), nonetheless accepted that taxonomists often resort to morphological distinctness as a proxy for genetic divergence or reproductive isolation. However, morphological and genetic divergence are not always tightly correlated, as evidenced by morphologically similar species and highly variable species, resulting in lumping or splitting of a morphospecies-based taxonomy when genetic data become available. Indeed, the rise of genomic-scale data has the potential to resolve many cryptic species complexes and to stimulate the identification of new species (Womack et al., 2022).

Poison frogs (Dendrobatidae), with 339 species (AmphibiaWeb, 2023), are among the most chromatically diverse amphibians and include some of the most exceptional cases of color polytypism and aposematism (i.e., the co-occurrence of warning signal and defense) among terrestrial vertebrates. For example, *Oophaga pumilio* includes 15 color morphs distributed from Nicaragua to northern Panama (Siddiqi, 2004). *Ranitomeya imitator* has four color morphs in a Müllerian mimicry complex including *Ranitomeya variabilis, R. summersi,* and *R. fantastica* (Brown et al., 2011; Symula et al., 2001; Twomey et al., 2016). Other examples of highly variable species include *Oophaga histrionica*, *Dendrobates tinctorius*, *Dendrobates auratus*, and the *Ameerega picta* complex (Hauswaldt et al., 2011; Brusa et al., 2013; Galindo-Uribe et al., 2014; Valero et al., 2017). On the other hand, some of the less colorful taxa (e.g., *Silverstoneia, Hyloxalus, Colostethus*) are notoriously difficult to distinguish based on external phenotype. These extreme examples indicate that dendrobatid species can be difficult to delimit without multiple lines of evidence.

*Epipedobates*, known as Chocoan poison frogs, with extensive variation in color pattern yet low genetic divergence among some species (<3% uncorrected p-distance in *16S*), is an example of challenges for species delimitation based on variation in color and pattern. *Epipedobates* contains both aposematic (brightly colored and chemically defended) species such as *E. tricolor* and inconspicuously colored species such as *E. boulengeri*. Accumulating evidence suggests that all *Epipedobates* species sequester lipophilic alkaloids (Santos et al., 2016; Tarvin et al., 2016, Cipriani and Rivera 2009; Gonzalez et al. 2021), although the efficacy with which the chemical defenses repel predators and parasites is unknown. *Epipedobates anthonyi* is the source of the toxic alkaloid epibatidine (Badio and Daly, 1994; the species from which epibatidine was first isolated was incorrectly identified as *E. tricolor*). Although all poisonous dendrobatids are known to obtain their alkaloids from dietary sources, the ultimate source of epibatidine is unknown. All *Epipedobates* species for which data are available have a unique set of amino acid substitutions in a nicotinic acetylcholine receptor that likely provides the frogs with resistance to epibatidine (Tarvin et al., 2017b). The seven recognized species of *Epipedobates* are found on the western side of the Andes from northern Peru to central Colombia: *Epipedobates anthonyi, E. boulengeri, E. darwinwallacei, E. espinosai, E. machalilla, E. narinensis,* and *E. tricolor*. Lötters et al. (2007) and Tarvin et al. (2017a) have provided the most thorough species-level treatments of *Epipedobates*.

Preliminary phylogenies of *Epipedobates* revealed complex patterns of diversification during the last 5 Myrs (Santos et al., 2009). Tarvin et al. (2017a) analyzed phylogenetic relationships of four of the seven species of *Epipedobates*, focusing on the origins of aposematism in *E. tricolor* and *E. anthonyi*. They found that low genetic divergence in the mitochondrial *16S* gene (<3% uncorrected p-distance) among *E. anthonyi, E machalilla*, and *E. tricolor* markedly contrasts with their high phenotypic disparity. The mating calls vary among some populations that are considered conspecific, suggesting prezygotic isolation, whereas in some allopatric species the calls are very similar, suggesting the absence of reproductive isolation (Santos et al., 2014; Tarvin et al., 2017a).

Tarvin et al. (2017a) provided the most recent review on the molecular systematics of the genus, but examined only a few samples of *E. boulengeri*, a nominal species widespread in northern Ecuador and southwestern Colombia, and they did not sample *E. darwinwallacei, E. espinosai*, or *E. narinensis*, which have narrow and relatively unstudied distributions. We expand that work with genetic analyses of all nominal species, including several additional populations of *E. boulengeri* and the first genetic data from *E. narinensis*. We also catalog the phenotypic diversity with emphasis on coloration and other anatomical features. We noted a surprisingly deep genetic divergence among samples identified as *E. boulengeri* and other species without bright coloration (hereafter, inconspicuous), which suggested an older divergence for the genus than previously recognized, to the upper Miocene (∼11 Mya). We revise and update the current taxonomy including new distributional information.

## 2. Material and methods

### 2.1. Sampling

This study was based on fieldwork during 2012, 2014, and 2016 by RDT, KLH, MBC, and SRR (see Supplementary Table S1 for a list of specimens) under the permits Res 1177-2014 by Autoridad Nacional de Licencias Ambientales-ANLA (Colombia), Res. 061-2016 by Parques Nacionales Naturales de Colombia (Colombia), Contrato Marco Acceso a los Recursos Genéticos Nro. 005-14 IC-FAU-DNB/MA (Ecuador). We examined genetic and phenotypic data for 116 individuals of *Epipedobates* that were initially identified in the field based on color pattern and proximity to type locality. Frogs were collected from 29 localities from southwest Colombia (Cauca, Valle de Cauca and Nariño) to southwest Ecuador (El Oro), including the entire range of *E. boulengeri*. We also sampled at or near the type localities of five of the seven known species: *E. boulengeri, E. darwinwallacei, E. machalilla, E. narinensis,* and *E. espinosai* (Fig. 1, type localities indicated by stars). The animal use protocol was approved by the University of Texas at Austin (IACUC AUC-2012-00032). Voucher specimens are deposited in the Museo de Zoología (QCAZ) de Pontificia Universidad Católica del Ecuador (PUCE) and Museo ANDES at Universidad de los Andes, Bogotá, Colombia. Throughout our paper we use globally unique identifiers (GUIDs) to refer to the voucher specimens at Museo de Zoología de la Pontificia Universidad Católica del Ecuador, Quito, Ecuador (QCAZ:A:XXXXX) and at Museo de Historia Natural C.J. Marinkelle at the Universidad de los Andes in Bogotá, Colombia (ANDES:A:XXXXX) in an effort to facilitate future machine-readability of this text and to adhere to Biodiversity Information Standards (Globally Unique Identifiers Task Group, 2011). In some cases, we also note field series numbers (RDT, Rebecca D. Tarvin, and TNHC-FS, Texas Natural History Collections Field Series). The specimens reviewed in this work are listed in Table S1.

**Figure 1.**
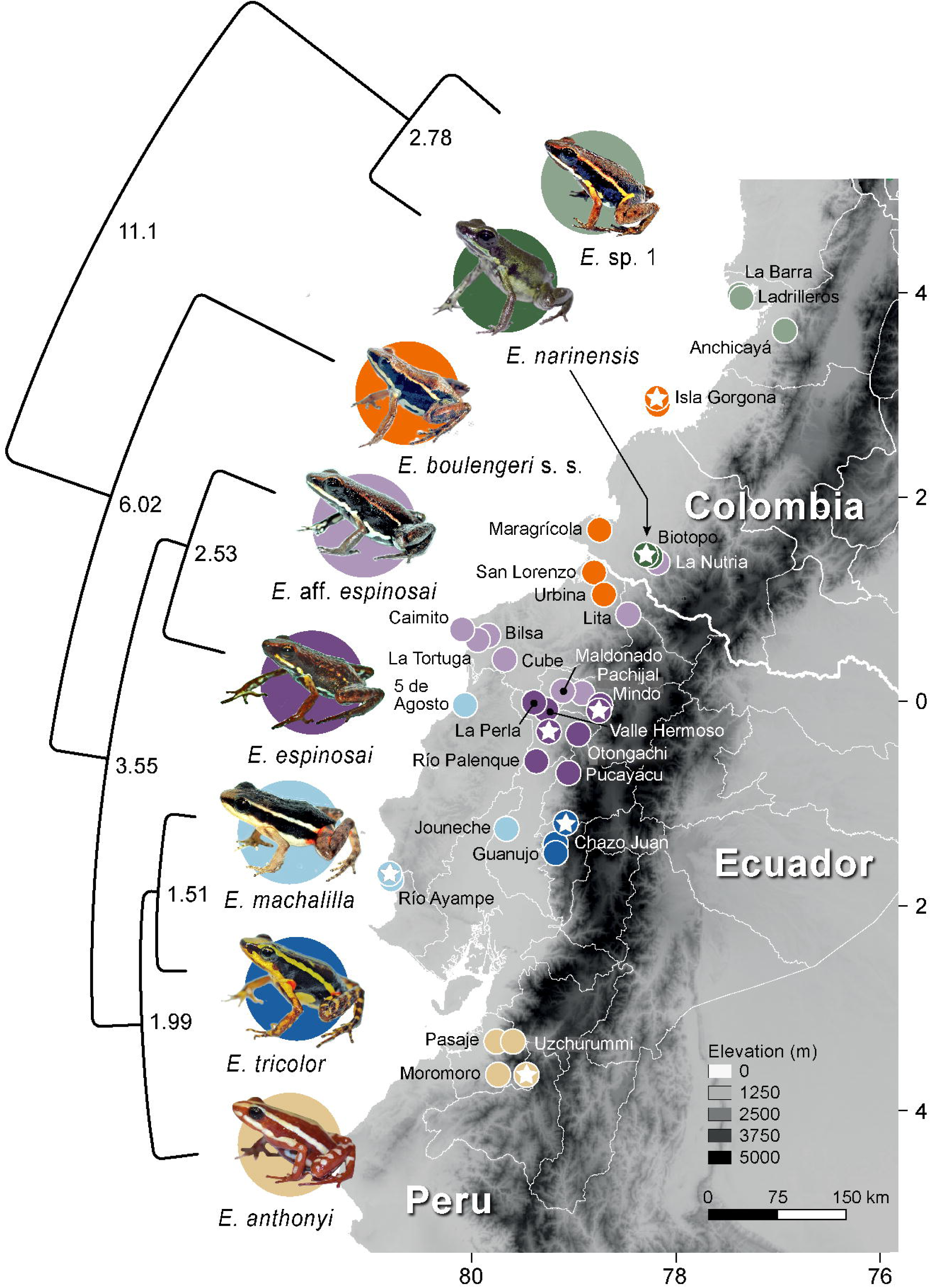
Distribution of sequenced individuals of *Epipedobates*, their type localities (stars), and estimated divergence times. The tree is an abbreviated chronogram based on Supplementary Fig. S2, and the node labels represent estimated ages in millions of years.

### 2.2. DNA sequencing

DNA was extracted from liver and muscle tissue using the Qiagen DNeasy Blood & Tissue kit (Valencia, CA) following the manufacturer’s protocol. We sequenced five gene fragments for 46 *Epipedobates* individuals: two nuclear markers (total length of 878 bp), histone H3 (*H3*), and bone morphogenetic protein 2 (*BMP2*); and three mitochondrial fragments (total length of 2412 bp), a fragment including parts of the 12S and 16S mitochondrial rRNA genes (*12S–16S*) and the intervening valine tRNA gene (*TRNV*), cytochrome b (*CYTB*), and the mtDNA control region (*CR*). The primer sequences and protocols are provided in Supplementary Table S2; the PCR protocols followed Goebel et al. (1999), Santos and Cannatella (2011), and Santos et al. (2003). PCR products were sequenced in both directions at the Institute for Cellular and Molecular Biology Core Facility at the University of Texas at Austin. The chromatograms were edited in Geneious 5.4.4 (Kearse et al., 2012), and forward and reverse reads were merged into consensus sequences for further analyses. New sequences were deposited in GenBank (accession numbers OR488977–OR48901, OR179756–OR179880, and OR734663–OR734707).

### 2.3. Phylogenetic analyses

We augmented our data with 74 GenBank sequences (accession numbers in Supplementary Table S1) of the voltage-gated potassium channel 1.3 (*K_v_1.3*), cytochrome b (*CYTB*), and *12S–16S* from 70 individuals included in Tarvin et al. (2017a). The combined dataset included sequences of *12S–16S* and *CYTB* for all individuals, but not for *H3*, *BMP2*, and *CR* (see Supplementary Table S1 for details). To estimate phylogenetic relationships, we constructed a concatenated matrix of six gene fragments with a total length of 4636 bp. In some cases the sequence data of an outgroup species includes a chimera of sequences of two individuals (Supplementary Table S1) to ensure maximum amount of sequence information. *Silverstoneia nubicola, Oophaga pumilio, Dendrobates sp., Dendrobates auratus*, *Oophaga histrionica, Adelphobates galactonotus, Dendrobates leucomelas, Leucostethus fugax,* and *Phyllobates aurotaenia* were used as outgroups. We note there is controversy over the elevation of subgenera such as *Oophaga* to genera (Grant et al., 2017; Santos et al., 2009), and we have followed the taxonomy of AmphibiaWeb (2023).

Sequences were aligned with MAFFT version 7 using “--auto” to choose the best alignment strategy (Katoh and Standley, 2013). Alignments were then slightly adjusted manually in Mesquite 3.2 (Maddison and Maddison, 2018) to fix obvious errors in alignment. The alignment is provided in Supplementary Data S1.

Maximum likelihood (ML) and Bayesian inference (BI) methods were used to estimate the phylogeny using the concatenated matrix. The protein-coding genes were partitioned by gene and codon position; the *12S–16S* fragment was not partitioned. Because the first and second codon-positions of the *BMP* and *H3* genes were invariant across all samples, including the outgroups, we excluded these from analysis. IQ-TREE v.2.1.2 (Kalyaanamoorthy et al., 2017; Minh et al., 2020) was used to select simultaneously the model of evolution and data partitions (– m TESTMERGE) under the Bayesian Information Criterion. The base-frequency parameters tested included equal and estimated base frequencies (–mfreq F, FO). Rate heterogeneity parameters included equal, gamma, and invariant (–mrate E, G, I). IQ-TREE2 (Minh et al., 2020) was run five times to check consistency of likelihood scores; all runs yielded the same topology and similar log-likelihood values (within a range of one likelihood unit). Ten thousand ultrafast bootstrap replicates (Minh et al., 2013) were used to assess branch support, and bootstrap proportions were plotted onto the optimal likelihood tree (Supplementary Fig. S1).

To check for the effect of missing data, we ran the ML analysis on *12S–16S* and *CYTB* partitions, which have almost complete data for every individual. This yielded the same topology of relationships among species (Supplementary Fig. S1), so we used alignment of all genes.

Bayesian inference (BI) analyses with nonclock and clock models were performed with MrBayes 3.2.7 (Ronquist et al., 2012) on the CIPRES portal (Miller et al., 2015) using the character partitions from the IQ-TREE analysis; model averaging was used to sample all possible reversible substitution models (*nst = mixed*). Among-site rate variation was modeled using the gamma distribution (*rates = gamma*). The priors for state frequency, rate multipliers, substitution rates, and shape parameter of the gamma distribution were unlinked among partitions, and the branch lengths were linked. The nonclock analysis was conducted with two runs of four chains each for 4 million generations, sampling every 500 generations. The MCMC posterior samples were visualized using Tracer 1.7 (Rambaut et al., 2018). The two runs were judged to be successful when (1) the success of swaps between chains was between 15-75%, (2) the Average Standard Deviation of Split Frequencies was <0.01, (3) the Potential Scale Reduction Factor (Gelman and Rubin, 1992) of parameters was between 1.00 and 1.02, (4) combined ESS values of almost all parameter estimates were above 200 (except in a few cases), as assessed by Tracer, and (5) the difference in the marginal likelihoods of the two runs was <6.

A Bayesian chronogram was constructed using MrBayes 3.2.7 on the CIPRES portal (Miller et al., 2015) with the same partitioned substitution model described above, under the Independent Gamma Rates relaxed clock model (IGR, Lepage et al., 2007). Preliminary analyses using all tips suggested that the large number of tips with low genetic divergence made it difficult to estimate the variance parameters of the branch lengths under the IGR model, so we pruned the matrix to 18 tips representing 2-3 individuals of each species (Supplementary Fig. S2). The tree age prior was drawn from a truncated normal distribution with mean 28.99 and standard deviation 3.0 to approximate the results of Feng et al. (2017) in which the estimated root node age was 28.99 Myrs with 95% credible intervals of 23.21–35.30 Myrs. The clock rate prior was drawn from a normal distribution with mean 0.005 and standard deviation of 0.001. The appropriateness of these priors was evaluated by comparing the induced prior distributions with the posterior distributions. In MrBayes, this is done by running the MCMC chain without data (*mcmc data = no*).

The analysis was conducted with two runs of four chains each for 200 million generations, sampling every 10000 generations, yielding 10000 samples from the two runs. The first 25% were discarded as burnin, so 7500 samples were retained. The credibility intervals of node ages are given as the region of 95% highest posterior density (HPD). The MrBayes script is provided in Supplementary Data S1.

### 2.4 Quantitative species delimitation

We used two distinct programs to estimate species limits in our dataset. The first, Assemble Species by Automatic Partitioning (ASAP; Puillandre et al., 2021), estimates pairwise distances of single loci to cluster sequences hierarchically and to identify putative species groups based on intra-versus inter-group distances. We ran ASAP using the Jukes-Cantor substitution model and an alignment of all mtDNA sequences (excluding nDNA) and a seed value of 123.

For the program to run, we needed to remove QCAZ:A:53636 and QCAZ:A:53641, which did not have any sequence overlap with some of the other sequences. Second, we used a Generalized Mixed Yule Coalescent analysis (GMYC; Pons et al., 2006; Fujisawa and Barraclough, 2013). The GMYC is a single-locus species delimitation method, which uses an ultrametric tree and its associated lineage-through-time plot (LTT) to infer the transition point between those branching events caused by speciation (the Yule birth-only process) and those caused by allele coalescence (Hudson, 1990). Because speciation should occur at a slower rate than allele coalescence, the model tries to identify the node at which the rate of branching dramatically increases (modeled by T, the threshold parameter).

For each internal node, the algorithm calculates the likelihood that the node is the transition point from a speciation (Yule) model to a coalescent model. The optimal threshold is the age (in substitutions/site) of the node with the maximum likelihood. The largest cluster of tips whose most recent common ancestor is less than or equal to this threshold is inferred to be a species. The total number of inferred species (“entities”) is calculated as the sum of the number of clusters (having more than one tip) and the number of singleton tips that diverged prior to the threshold. The 95% confidence limits of the number of species were determined by the number of species for all nodes within two units of the likelihood of the optimal node.

The GMYC analysis was performed using the gmyc() function in the splits R package, version 1.0-20 (Ezard et al. 2021), using the single-threshold model (Supplementary Data S2D-J). The null hypothesis is that the coalescent process explains the branching pattern of the entire tree. We used a likelihood-ratio test to assess whether our observed tree fit this model (Supplementary Data S2G).

To create an ultrametric tree for the GMYC analysis we used MrBayes as described above, with slight modifications to the alignment (Supplementary Data S2D). Duplicate sequences, outgroup sequences, and nuclear genes were removed. Details of the partitions, parameters, and priors are given in the MrBayes script (Supplementary Data S2F). The IGR relaxed clock model was parameterized with a fixed clock rate of 1.0 to remove the uncertainty of clock rate (because determining age was not necessary). Two analyses of four Markov chains were run for 50 million generations, sampled every 2000 generations, with a burn-in of 25%; this produced 37,500 posterior samples.

One GMYC analysis was performed on the MrBayes ultrametric consensus tree (Figure Z) to infer the number of species and the 95% confidence. A GMYC analysis of one tree does not account for the uncertainty in tree estimation. To take this into account, for the second analysis we used the last 1000 samples of the posterior distribution of trees of the MrBayes GMYC analysis (Supplementary Data S2H). A separate GMYC analysis was performed on each tree, and we used an R script (Supplementary Data S2G) to summarize (1) the number of estimated species and the proportion of significant (p < 0.05) tests over 1000 trees using histograms.

Trees were visualized using FigTree 1.4.4 (Rambaut, 2018). The maximum clade credibility tree (MCCT; Figure 2) was calculated using SumTrees from the DendroPy package (Sukumaran and Holder, 2010a, 2010b). R scripts (R Core Team, 2021) and the packages *ape* 5.6-2 (Paradis and Schliep, 2019) and *phytools* 1.2-0 (Revell, 2012) were used to plot the symbols for support values on the trees. Scripts are provided in (Supplementary Data S1). The outgroups were pruned from the tree figures to save space; complete tree files are available in Supplementary Data S1.

**Figure 2.**
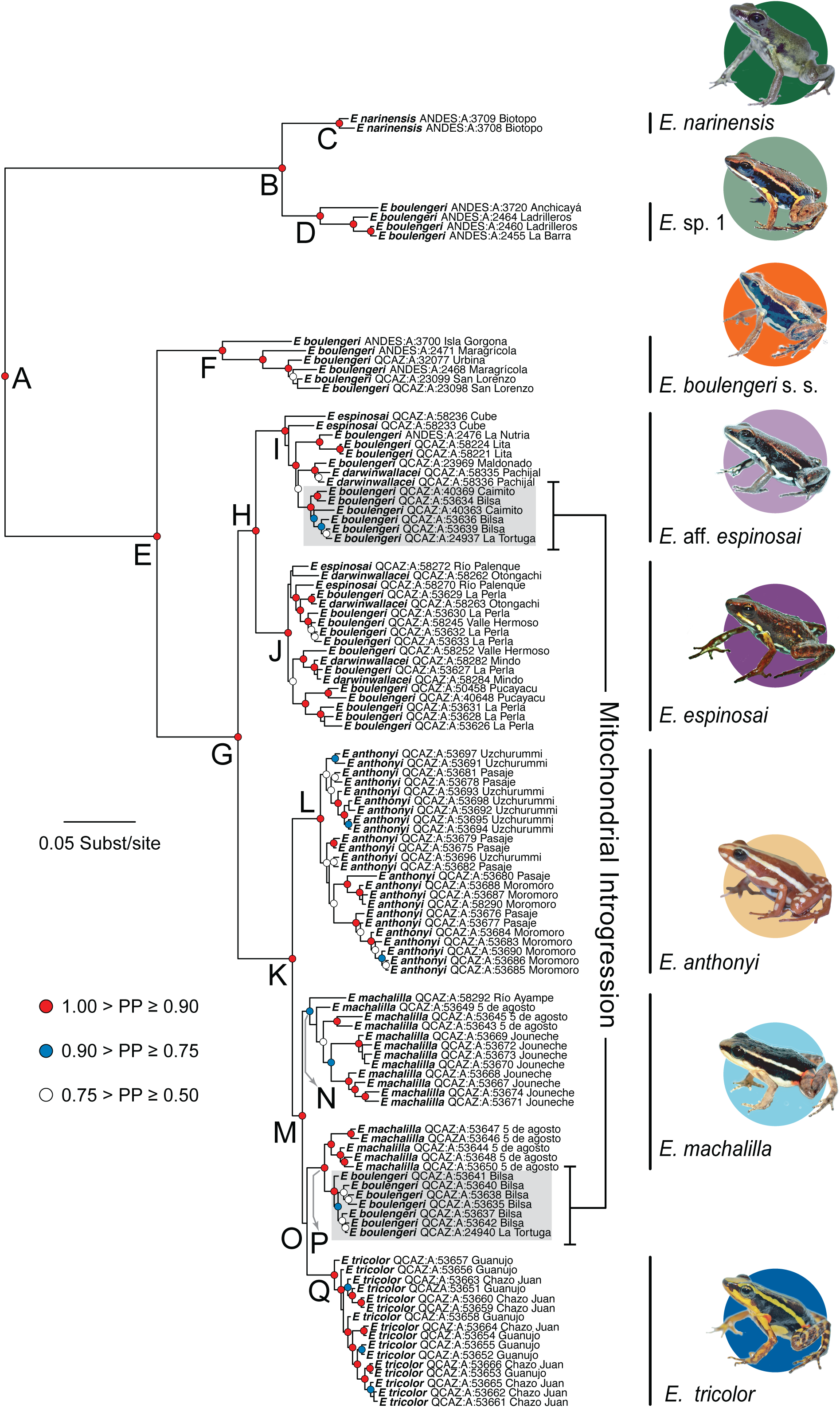
Bayesian phylogeny of *Epipedobates* based on *12S–16S, CYTB, CR, BMP2, H3*, and *K_V_1.3 gene sequences*. Tip labels correspond to initial identifications made in the field. Revised species identifications were made based on molecular clade membership (see photos and names on the right). See the text for further discussion of the revised species boundaries.

### 2.5. Genetic variation

We calculated mean pairwise p-distances (uncorrected) between species with an R script (Supplementary Data S1) with *ape*, using pairwise deletion for missing data. We excluded the individuals from the *E.* aff. *espinosai* Bilsa and La Tortuga localities, which show evidence of introgression with *E. machalilla*.

Haplotype networks were constructed for each gene using the *pegas* 1.1 package (Paradis, 2010) with modified script by Paradis (2021). The nuclear genes were phased using SeqPHASE (Flot, 2010) and PHASE 2.1.1 (Stephens et al., 2001); the R script and the alignments used for the network analysis are provided in Supplementary Data S1. For each gene, a network was built using the number of differing sites (Manhattan distance) between each pair of haplotypes to create a randomized minimum spanning haplotype network (Paradis, 2018), which displays equally parsimonious connections between haplotypes, from 10000 iterations. The R script is included in Supplementary Data S1.

### 2.6. Phenotypic data

Specimens were preliminarily identified to species in the field using color pattern and geographic location, which are the most obvious ways that the nominal species differ. We compared frogs in life and our color photographs (Supplementary Figs. S4-S27) with published descriptions and taxonomic keys (e.g., Cisneros-Heredia and Yánez-Muñoz, 2010; Funkhouser, 1956; Grant et al., 2006, 2017; Mueses-Cisneros et al., 2008; Myers, 1987; Silverstone, 1976; Tarvin et al., 2017a). Briefly, the color and pattern characters included the presence or completeness of the oblique lateral stripe (Edwards 1974; different than dorsolateral stripe), paracloacal mark (Grant et al. 2006), and ventrolateral stripe (Edwards 1974); the presence of flash spots in the inguinal region and concealed surfaces of the thigh and shank, and color pattern of dorsum, venter, flanks, head, and limbs. See the images of *E. tricolor* in Figure 5 for a visual guide and characterization of the phenotypic diversity of each species in Section 3.4 below.

In addition, we review the morphology of the tarsal keel (also known as tarsal fold) because it was used to diagnose *E. darwinwallacei* (Cisneros-Heredia and Yánez-Muñoz 2010), which our data suggest is not distinct from *E. espinosai*. The tarsal keel is a fold of skin (Silverstone, 1975; Grant et al., 2006) that in its most extreme condition extends from the lateral margin of the inner metatarsal tubercle, diagonally across the plantar surface of the tarsus, terminating in a tarsal tubercle located about one-fourth to one-third of the length of the tarsus. Because the shape of the tarsal keel was the only morphological trait used in the diagnosis of *E*. *darwinwallacei*, we compared this structure in species of *Epipedobates* using the character-state definitions of Grant et al. (2006: Fig. 31). The states were scored “blind” by DCC.

“0: Straight or very weakly curved, extending proximolaterad from preaxial edge of inner metatarsal tubercle;

1: Tubercle-like (i.e., enlarged), and strongly curved at proximal end, extending from metatarsal tubercle.

2: Short, tubercle-like, curved or directed transversely across tarsus, not extending from metatarsal tubercle.

3: Weak, short dermal thickening, not extending from metatarsal tubercle.”

The following specimens or photographs where indicated were examined for tarsal keel morphology. For *E. espinosai*: QCAZ:A:23783 (Santo Domingo. 9 Km vía Chone; photos), QCAZ:A:38058 (Centro Científico Río Palenque). For *E. machalilla*: QCAZ:A:10725 (male) and QCAZ:A:48557 (female). For *E. anthonyi*: QCAZ:A:65862 (male) and QCAZ:A:65864 (female). For *E. tricolor*: QCAZ:A:49821 (male) and QCAZ:A:49819 (female). For *E. darwinwallacei*: QCAZ:A:35612 (female, photos, Mindo) and QCAZ:A:35615 (female, Mariposario de Mindo), QCAZ:A:12807 (male, Mindo).

We measured snout-vent length (SVL) in living and museum specimens and, when possible, determined the sex by examination of gonads and color of the throat (in mature males the pattern of the throat tends to be obscured or disrupted by dark gray or diffuse pigmentation). Because most specimens were used to harvest skin and tissues for alkaloid and phylogenetic analyses, we were not able to make detailed measurements for morphometric analysis. We compared SVL among species, by sex, and by elevation using the *stats* package in R v4.1.2 and the *rstatix* 0.7.0 package (Kassambara, 2021). First, we tested for phylogenetic signal in SVL and elevation separately using Pagel’s λ test (*phylosig* function from the *phytools* R package). In both tests, λ was < 0.0001 (p = 1.0), indicating a lack of signal, so we used standard regression models. We performed an ANOVA using the *aov* function with sex, species, and their interaction as predictors, and SVL as the response variable. The residuals of this model were not normally distributed (Shapiro Wilk normality test, W = 0.96, p = 0.00066). Residuals from a similar model with sex as the only predictor and SVL as the response variable were also not normally distributed (Shapiro Wilk normality test, W = 0.97, p = 0.0071). Thus, we separated data into female-only and male-only datasets with species as the predictor and SVL as the response variable. For the female dataset, the null hypothesis was not rejected, but marginally so (W = 0.97, p = 0.055). The residuals for the male dataset were not normally distributed (W = 0.94, p = 0.0035).

We proceeded to perform an ANOVA followed by a Tukey test (*tukey_hsd* function from *statix*) to estimate significant pairwise differences in SVL among species for females. For males, we used non-parametric Kruskal Wallis rank-sum tests (*kruskal.test* function from the *stats* package) and the Dunn’s Test of Multiple Comparisons (*dunn_test* function from *rstatix* R package) to conduct post-hoc pairwise tests corrected for multiple testing. To compare SVL between sexes within each species, we checked for normality of residuals from a linear model (*lm* with sex as predictor and SVL as response variable). If residuals were normal, we used a Student’s t-test (*t.test* in the *stats* package) to compare male and female SVL. Otherwise, we used a Wilcoxon rank-sum test (*wilcox.test* in *stats* package). Finally, to compare SVL and elevation, we tested the normality of residuals from a linear model with elevation as predictor and SVL as the response variable for datasets subsetted to males only, females only, and all individuals excluding juveniles. We also repeated these analyses using means at each locality instead of individual data. If the residuals were not normal, we proceeded with a one-sided non-parametric test, Spearman’s *rho (cor.test* with “method = spearman” and “alternative = greater”); otherwise we used a linear model.

### 2.7. Species distributions

A list of collection sites was derived from bibliographic and museum data and our own collection efforts (Supplementary Data S3). Localities were georeferenced where necessary; only records with a museum voucher were included. Throughout the text we refer to our collection sites with simple names such as “Mindo” and “Río Ayampe”. However, these names represent the general area where the specimens were collected and not the exact locality; i.e., “Mindo” is not only the town of Mindo, but also its vicinity. Detailed georeferenced localities are provided in Supplementary Table S1. Distances (great circle distance) between localities were calculated using the *geodist* R package (Pagdham 2021).

## 3. Results

### 3.1. DNA sequencing

We obtained a total of 4636 bp: *12S–16S*: 611 bp (64 parsimony-informative sites), *CYTB*: 657 (146), *CR*: 1821 (349), *H3*: 346 (1), *BMP2*: 532 (7), *K_V_1.3*: 1349 (31). The first and second codon-positions of *BMP2* and *H3* had no variable sites. See Supplementary Table S1 for GenBank accession numbers).

### 3.2. Phylogenetic analyses

Maximum likelihood analysis of the partitioned dataset recovered log-likelihoods of – 22563.498 to –22563.543 (three replicate analyses) for a five-partition scheme chosen under the Bayesian Information Criterion, BIC (Supplementary Data S1 and Fig. S1). The nonclock Bayesian analysis (Fig. 2) and the ML analysis shared the same strongly supported clades (95– 100% BS or 0.95–1.00 PP). The harmonic means of the two replicate Bayesian runs were – 23381.14 and –23376.16, with a 95% credible set of 11882 trees. A burnin of 25% of the samples was judged to be sufficient; the average effective sample size (ESS) was > 800 for all parameters; the average standard deviation of split frequencies at the end of the run was 0.0076; the ratio of successful chain swaps was 0.53–0.57 for both runs, within the recommended range of 0.20 and 0.70.

All parameters in the Bayesian chronogram analysis yielded an ESS of > 5000, indicating that the number of generations was far above what was needed. A burn-in of 25% was sufficient. The chains were well-mixed; the percent of successful chain swaps was 0.60–0.89 for each run. Convergence between the two runs was excellent; the PSRF was 1.000 for all parameters and the final average standard deviation of split frequencies was 0.001475. The harmonic means of the two runs were –12695.20 and –12693.41. The 95% credible set contained 11 trees.

The chronogram (Fig. 1, Fig. S2) places the origin of the crown clade *Epipedobates* at 11.1 Myrs (95% HPD = 8.61–13.86 Myrs) and its divergence from *Silverstoneia* at 13.91 Myrs (11.01–17.22 Myrs). Because our sample of populations was not extensive (i.e., we may have missed samples with deeper divergences), these age estimates likely underestimate the true age of the most recent common ancestor of each species. Nonetheless, the estimates provide a useful minimum age.

Under the general lineage concept (de Queiroz et al. 2005, 2007), we hypothesize that the seven well-supported molecular clades diagnose species lineages, and we discuss their morphological distinctness below: *Epipedobates narinensis* (node C), *Epipedobates* sp. 1 (node D), *E. boulengeri* s. s. (node F), *E.* aff*. espinosai* (aff. = affinis, a closely related but different species) (node I), *E. espinosai* (node J), *E. anthonyi* (node L), and *E. tricolor* (node Q). We also hypothesize that individuals assigned to *Epipedobates machalilla* represent an eighth species even though the samples do not form a monophyletic group on the molecular phylogeny (Fig. 2; see also section 4.1.3). Some *E. machalilla* cluster with some La Tortuga and Bilsa individuals of *E.* aff. *espinosai* rather than other *E. machalilla*, but the support is very weak, similar to results of Tarvin et al. (2017a); this is not surprising because we included the Tarvin et al. sequences in our analysis. A revision of the species status of *E. machalilla* would be premature because the “paraphyly” of *E. machalilla* is poorly supported, we have few samples, and we are currently analyzing genome-scale data with additional samples to examine the possibility of mitochondrial introgression or more extensive gene flow.

The deepest split among *Epipedobates* (Fig. 2, node A, 11.10 Myrs, 95% HPD = 8.61– 13.86) is between the *E. narinensis + E.* sp. 1 clade (node B, 2.78 Myrs, 1.72–4.00) and the rest of *Epipedobates* (node E, 6.02 Myrs, 4.45–7.87). Node B includes not only *E. narinensis* individuals from the type-locality (Reserva Biotopo, Valle del Cauca, Colombia), but also individuals from western central Colombia that resemble *E. boulengeri*. We hypothesize that these individuals represent an undescribed species that we designate *Epipedobates* sp. 1.

A second deep and well-supported node (E) separates *E. boulengeri* s. s. (node F) from the rest of *Epipedobates* (node G). *Epipedobates boulengeri* s. s. contains individuals from Isla Gorgona (the type-locality), northwestern Ecuador, and southwestern Colombia at elevations of 0–136 masl; the divergence time between the mainland and island populations is 3.06 Myrs (1.87–4.46), which is slightly greater than that between *E. narinensis* and *E.* sp. 1.

Node H (2.53 Myrs, 1.71–3.53) includes two putative species we call *E. espinosai* (node J) and *E.* aff. *espinosai* (node I), which comprise specimens from north-central Ecuador. Each clade is strongly supported, but the sister-group relationship between the two is less so. The species delimited by Node J takes the name *E. espinosai* because it includes individuals from Río Palenque and Valle Hermoso (Fig. 1), which are near Hacienda Espinosa, the type-locality of *Phyllobates espinosai* Funkhouser 1956 (see section 4.5). This clade also contains individuals from Mindo, near the type locality of *Epipedobates darwinwallacei* Cisneros-Heredia and Yánez-Muñoz 2011. However, *E. espinosai* is the older name and has nomenclatural priority. Specimens from this clade are distributed at elevations from 152–1375 m (Provinces of Santo Domingo, Pichincha, Esmeraldas, and Los Ríos). Some individuals we initially identified as *E. boulengeri* also fall within this clade.

We designate the clade at node I as *E.* aff. *espinosai* because of its similarity to *E. espinosai*. Individuals identified in the field as *E. darwinwallacei, E. espinosai,* and *E. boulengeri* are found in this clade, but none of these names can be applied to this clade, the species status of which we are further investigating. This clade is allopatric to *E. espinosai* (Fig. 1), being distributed in northwestern Ecuador (Imbabura, Santo Domingo, Pichincha, and Esmeraldas) to southern Colombia (Nariño) between 87–907 m (Fig. 1).

Node K includes a shallow (1.99 Myrs, 1.27–2.95) radiation of individuals identified as *E. machalilla* (node P and part of node N), *E. anthonyi* (node L), *E. tricolor* (node Q), and some of the individuals identified as *E. boulengeri* from Bilsa and La Tortuga (subset of node N, indicated by the gray box in Fig. 2). The clustering of some individuals from Bilsa and La Tortuga with some *E. machalilla* is notable because other individuals from these two localities are grouped in *E.* aff. *espinosai* (Fig. 2, node I, gray box).

Node M (1.51 Myrs, 0.92–2.28) includes *E. tricolor*, all *E. machalilla*, and some of the Bilsa-La Tortuga individuals. However, the minimum age of divergence of *E. machalilla* from *E. tricolor* is 1.51 Myrs.

The *E. anthonyi* and *E. tricolor* clades (L and Q, respectively) are each strongly supported, although few localities were sampled.

### 3.2. Quantitative species delimitation

We ran two programs to estimate and evaluate genetic clusters (ASAP) or coalescent patterns (GMYC) in our data to help provide additional objective estimates of species hypotheses (see Supplementary Data S2 for input and output files). The top two ASAP results predicted that our dataset contained either 2 (asap-score = 7.50) or 66 species (asap-score = 9.50); the remaining top 8 results ranged from 56 (asap-score = 11.50) to 73 (asap-score = 12.00) groups (Supplementary Data S2A-C). The top ASAP result split the genus into two groups, with individuals that we designate as *E. narinensis* and *E.* sp. 1 as one group, and all other individuals as the other group (Supplementary Data S2C). The partition with 66 groups had more than half as many groups as individuals in the analysis (N = 106).

We ran GMYC with a timetree estimated from the three mtDNA genes and default parameters (Supplementary Data S2D-F). The maximum log-likelihood of the best GMYC model was 439.1158 and that of the null model was 436.2457. A likelihood-ratio test (likelihood ratio = 5.740, df = 2) indicated that the difference was marginally not significant (p = 0.05670). This phylogeny, inferred using only the mtDNA (Fig. 3), was generally consistent with the phylogeny inferred with nDNA and mtDNA (Fig. 2), indicating that the bulk of the phylogenetic signal in our data comes from the mtDNA.

**Figure 3.**
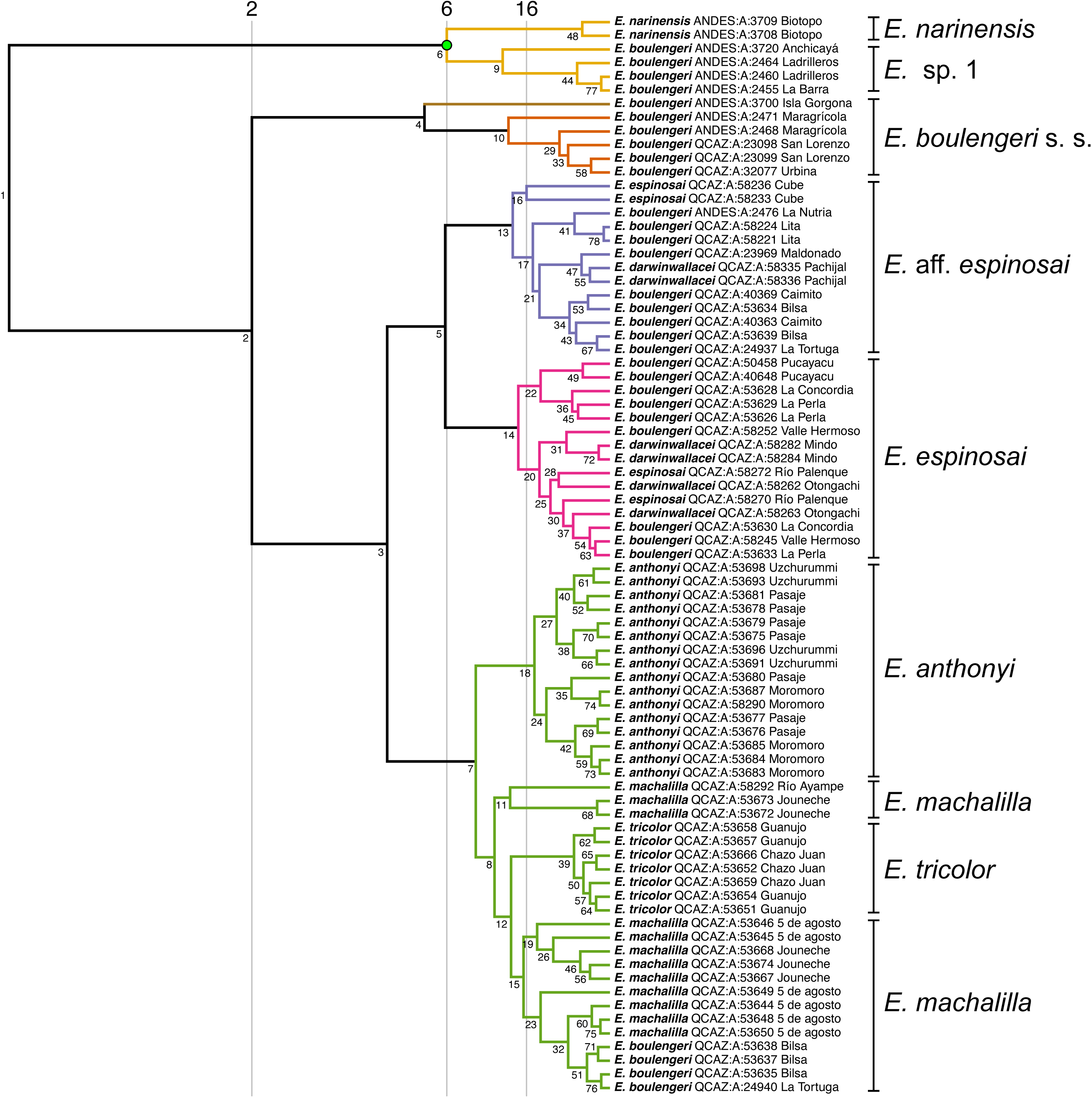
An ultrametric tree that was inferred with mtDNA and used for GMYC species delimitation estimation. The threshold with maximum likelihood is at Node 6 (node numbers assigned by the GMYC analysis), indicated by the green filled circle. Six species are identified, each in a different color, with 95% confidence intervals of 2–16, indicated by vertical lines. Our delimitation of species is given on the right side.

For the single GMYC analysis using the MrBayes consensus tree, the number of species estimated was six with a 95% confidence interval of 2-16 species (Fig. 3). The label of the optimal threshold was 6, with a threshold of –0.1152 (Supplementary Fig. S3; Supplementary Data S2). Accordingly, the five highest AIC weights (nodes 6, 8, 5, 7, and 9) were similar— 0.0974, 0.0909 0.0887, 0.0851, and 0.0726—indicating that no single threshold substantially influenced the results.

A summary of the 1000 GMYC analyses on the posterior sample of trees showed that the modal number of species is 5 (310/1000) and next largest is 6 (198/1000) (Supplementary Fig. S3; Supplementary Data S2G-J). 95% of the estimates of species numbers lie between 3 and 8, inclusive. Of the 1000 significance tests, 402 p-values were ≤0.05, indicating lack of support for rejecting the null hypothesis (Supplementary Fig. S3).

### 3.3. Genetic variation

#### 3.3.1. Genetic distances

We computed the mean uncorrected p-distances between clades/species (Fig. 4). Interspecific uncorrected p-distances range from 0.00925 (between *E. anthonyi* and *E. machalilla*) to 0.0737 (between *E. narinensis* and *E. boulengeri*) for *12S–16S* and from 0.0104 (between *E. tricolor* and *E. machalilla*) to 0.136 (between *E. narinensis* and *E. boulengeri*) for *CYTB*.

**Figure 4.**
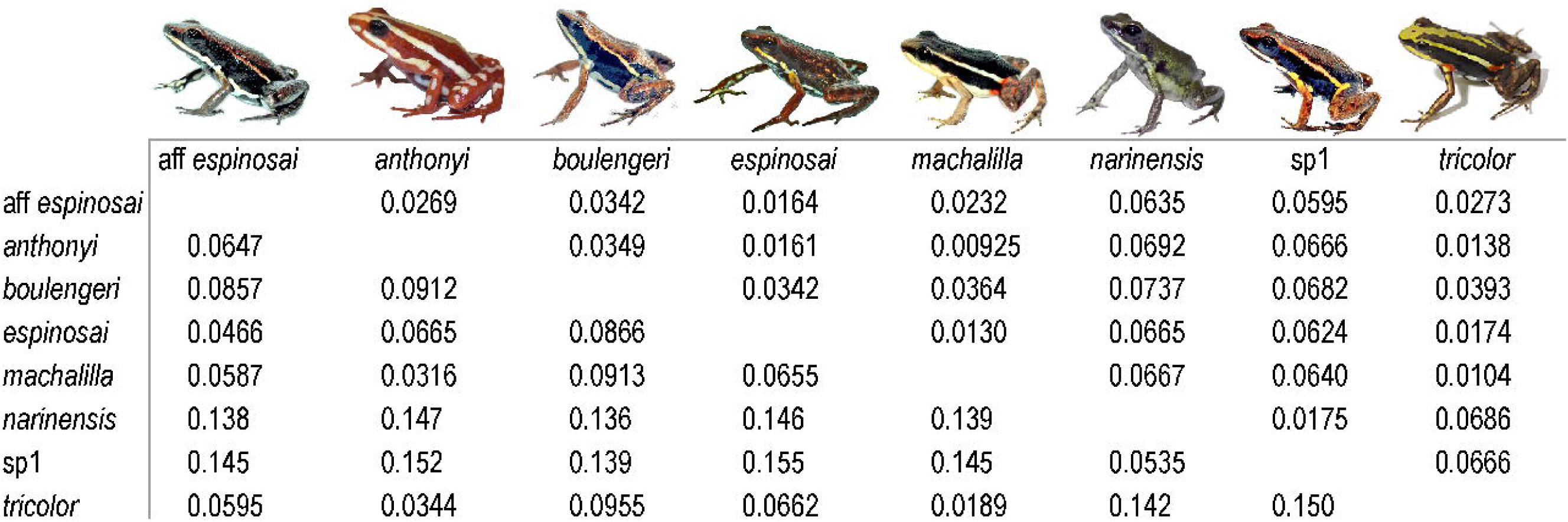
Matrix of mean pairwise uncorrected p-distances between clades. The lower triangle distances are from *CYTB* and the upper triangle distances are from *12S–16S*. Some species are highly divergent (e.g., *E. tricolor* and *E.* sp. 1 are 15% divergent in *CYTB*), while others are more closely related (e.g., *E. machalilla* and *E. anthonyi* are 3.1% divergent in *CYTB*).

#### 3.3.2. Haplotype analyses

We present randomized minimum spanning haplotype networks for the six evaluated genes in Fig. 5 (see also Supplementary Table S3). The number of haplotypes for each gene varies greatly. The three mitochondrial genes (Fig. 5ABC) have almost as many haplotypes as individuals and in no case does any species share haplotypes with another species. In the fast-diverging mtDNA control region (*CR*) only one haplotype is shared between two individuals; all other haplotypes are unshared among individuals (Fig. 5B). In contrast, the nuclear genes *H3* and *BMP2* (Fig. 5DE) have 7–8 times fewer haplotypes than phased sequences. *K_V_1.3* (Fig. 5F) had roughly half as many haplotypes as phased sequences, resulting in a lack of haplotype sharing among species except between *E. anthonyi* and *E. tricolor*. Note that some populations and species (*E.* sp. 1, *E. boulengeri* s. s., *E. narinensis*) are missing from the *K_V_.1.3* network because these sequences were pulled from a previously published dataset focusing on a different phylogenetic question (Tarvin et al. 2017a).

**Figure 5.**
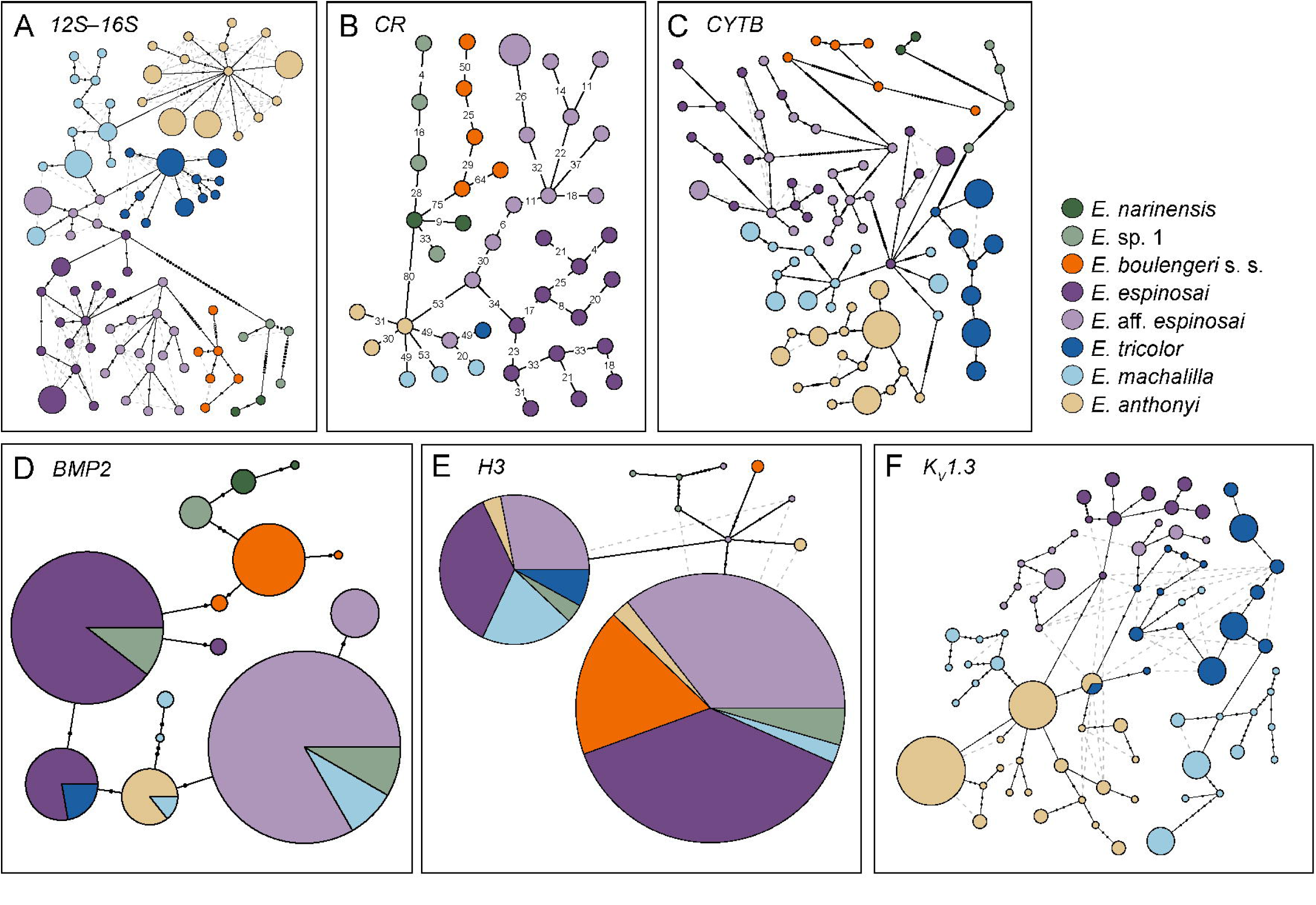
Randomized minimum spanning haplotype network for A) 12S and 16S mitochondrial rRNA genes (*12S–16S*), B) Control Region (*CR*), C) cytochrome B (*CYTB*), D) Bone morphogenetic protein (*BMP2)*, E) Histone 3 (*H3*), and F) voltage-gated potassium channel 1.3 (*K_V_1.3*). We note that sequences for *K_V_1.3* were pulled from previously published data, so sequences are unavailable for many of the focal lineages studied in this paper. Each circle represents a haplotype, and the diameter of the circle is proportional to the number of individuals having that haplotype. The color of the wedges identifies the species. Each black dot on the branches indicates one state change between haplotypes.

### 3.4. Phenotypic variation in color and pattern

We describe color and pattern variation for each of the putative *Epipedobates* species (Fig. 6).

**Figure 6.**
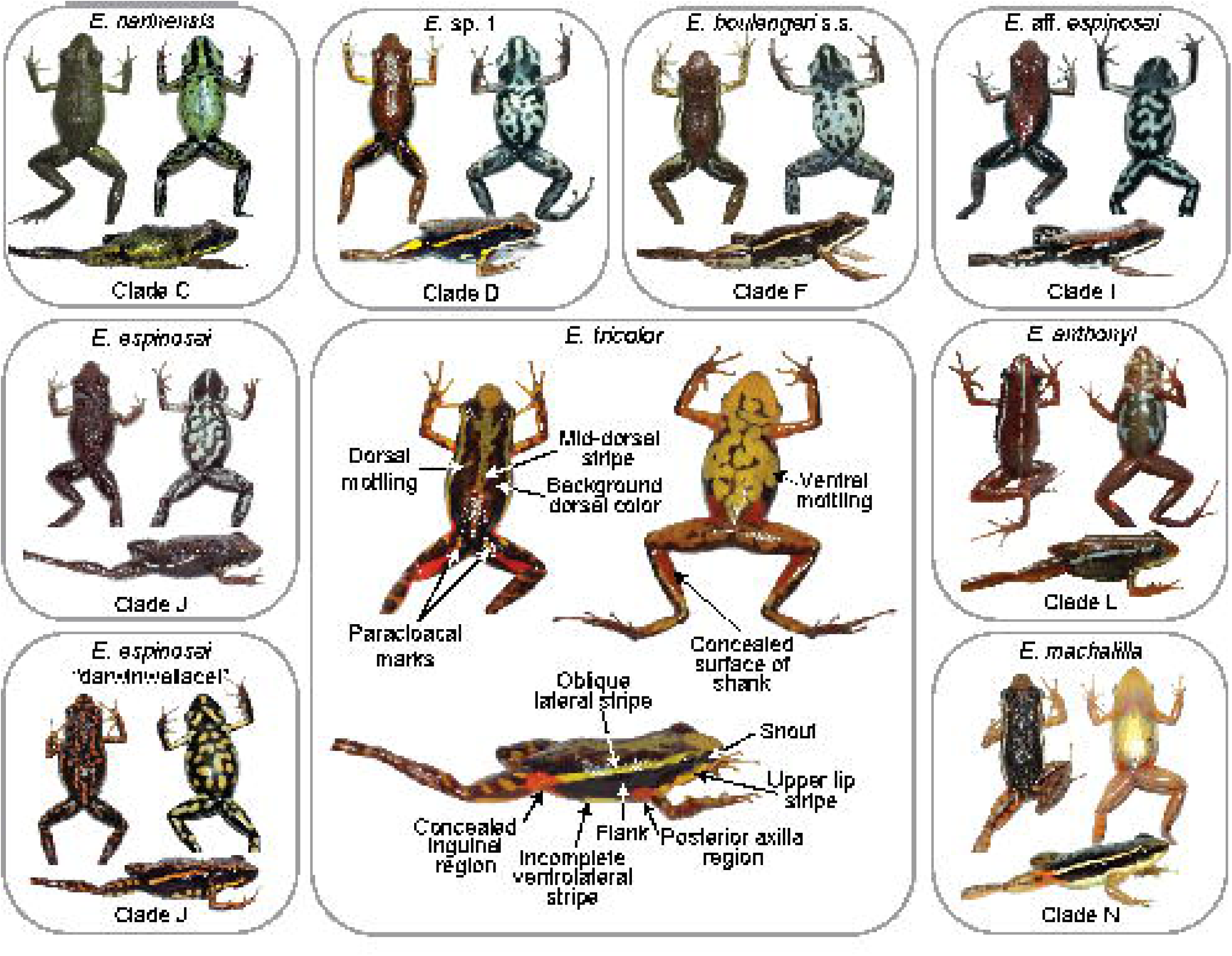
Phenotypic variation among species of *Epipedobates*. The letters correspond to clades in Fig. 2. The photographs are of the following individuals (all taken by RDT): node C, *E. narinensis* (ANDES:A:3709; RDT0220); node D, *E*. sp. 1 (ANDES:A:2455; RDT0147); node F, *E. boulengeri* s. s. (ANDES:A:3699, RDT0210); node I, *E*. aff. *espinosai* (QCAZ:A:58221, RDT0001); node J, *E. espinosai* (QCAZ:A:58271, RDT0052); node J, *E. espinosai* “darwinwallacei” (QCAZ:A:58282, RDT0062); node L, *E. anthonyi* (QCAZ:A:53698; TNHCFS-6875); node N, *E. machalilla* (QCAZ:A:58292, RDT0072); and node Q, *E. tricolor* (QCAZ:A:58307, RDT0087). See Table S1 for locality data.

#### 3.4.1. Epipedobates narinensis

Images of *Epipedobates narinensis* are represented by individuals from Biotopo, Colombia, the type locality (Figs. 1–2, node C; Supplementary Fig. S4) and the only known locality for this species.

Dorsal surfaces. The dorsal surfaces including the top of the snout are uniformly dark olive-green and lack markings.

Lateral and concealed surfaces. The flanks and sides of the snout of most examples are solid black, although some specimens have greenish yellow mottling on the flanks. The oblique lateral stripe is yellow-green and varies from being diffuse and indistinct to slightly discernible. In no specimen is it well defined, and in any case does not extend beyond the midlength of the body and fades anteriorly. Posteriorly it reaches the anterior inguinal region, which is black. In all individuals a thin, continuous greenish yellow white stripe on the upper lip extends from the snout tip to the axilla and in some it extends posteriorly as an indistinct ventrolateral stripe, reaching the inguinal region (Supplementary Fig. S4, ANDES:A:3710). However, in most frogs it is difficult to distinguish a ventrolateral stripe.

The concealed surfaces of the groin are black with diffuse olive mottling in some, and the black region extends onto the anterior concealed surface of the thigh. A diffuse, oval to elongate black blotch with a yellow-green suffusion lies on the anterior concealed surface of the thigh, but posterior to the inguinal fold. An elongate and diffuse yellow-green paracloacal mark (character 49, Grant et al. 2006) is present in some.

Ventral surfaces. The ventral surfaces of the throat, belly, thigh, and shank range from yellowish to greenish coloration, bright in some but mostly dull in luminosity. Some individuals have distinct black mottling, but overall most of the ventral surface lacks markings. Two irregular and elongate black blotches extend from the tip of the chin to the pectoral region. This pattern is somewhat obscured by diffuse gray-black pigment in the area of the vocal sac in males.

#### 3.4.2. Epipedobates *sp. 1*

Images of *Epipedobates* sp. 1 are from populations in southwest Colombia (Figs. 1, 2, node D; description of this new species is in progress) and include Anchicayá, Ladrilleros, La Barra, and Pianguita (Supplementary Figs. S5–S7). In general, the color pattern of the Anchicayá population differs slightly from that of the other three populations.

Dorsal surfaces. Specimens of the four populations have a uniformly brown dorsal coloration, lacking spots or blotches. The skin texture of the dorsum, head, thigh, and shanks is covered with tiny uniform tubercles, resembling a fine sandpaper.

Lateral and concealed surfaces. The flanks and sides of the snout are solid black and smooth. In the Ladrilleros population (Supplementary Fig. S6) they are separated from the dorsum by an orange-yellow oblique lateral stripe with weakly defined margins, extending from the anterior part of the groin forward to the eye, becoming more brownish orange than yellow, and onto the upper eyelid, and fading onto the canthus rostralis. In the Anchicayá population (Supplementary Fig. S5) this stripe is yellowish, and in some specimens it narrows, breaks into blotches, and fades out before reaching the eye. In some Pianguita individuals (Supplementary Fig. S7) the oblique lateral stripe is periodically broken along its entire length. In all populations a distinct gap separates the posterior end of the oblique lateral stripe from the blotch on the anterior surface of the thigh. This gap is much larger than that of *E. boulengeri* s. s. In all individuals a white to creamy upper labial stripe is present, distinctly paler than the oblique lateral stripe, and in some slightly iridescent, extends from below the nostril to the axilla. In some, it extends posteriorly as a poorly defined ventrolateral stripe (Supplementary Fig. S7, ANDES:A:3693).

The concealed surfaces of the groin are dark; posterior to the inguinal fold and the anterior concealed surface of the thigh lies a somewhat diffuse, oval to elongate yellow (Anchicayá) or orange-yellow to whitish yellow (Ladrilleros, Pianguita, and La Barra) blotch. A similar, but smaller and more elongate paracloacal mark is present on the posterior surface of the thigh in the latter three populations. Only a couple of the Anchicayá frogs have a hint of a yet smaller, yellowish mark; it is absent in other individuals. Most individuals have a yellow patch on the upper arm matching the color of the blotch on the anterior concealed surface of the thigh. These yellow patches are qualitatively different from the flash marks in the *E. anthonyi + E. machalilla + E. tricolor* clade (node K), which are situated more anteriorly, on the concealed surfaces of the groin area at the confluence of the thigh and flanks.

Ventral surfaces. In the Anchicayá population the ventral surfaces of the throat, belly, thigh, and shank are heavily mottled, with a pale blue turquoise background and irregular black blotches and vermiculations. Two irregular and elongate black blotches extend from the tip of the chin to the pectoral region. In males this pattern is obscured or disrupted by diffuse dark gray pigment just anterior to the pectoral region in the area of the vocal sac. A few individuals have yellowish diffuse coloration on the outer edges of the light blue surface of the belly.

The ventral surfaces in the other three populations are similar except that they range from paler blue to almost white. The variation in the ratio of dark to light areas is larger in these populations. In some individuals the pattern on the ventral thighs is almost absent. The yellowish tinge at the outer edge of the surface of the belly is present in some individuals.

#### 3.4.3. Epipedobates boulengeri *s. s*

Images of *E. boulengeri* s. s. (Figs. 1, 2, node F) are represented by populations from Isla Gorgona (the type locality) and Maragrícola on mainland Colombia (Supplementary Figs. S8-S9).

Dorsal surfaces. In both populations the dorsal surfaces are dark brick red to dark brown; a couple of individuals show a slight mottling at mid-dorsum, with darker, almost black regions adjacent to the oblique lateral stripe. Overall, the dorsal colors are slightly paler than those of *E.* sp. 1.

Lateral and concealed surfaces. The flanks and sides of the snout are solid black and smooth. In both populations they are separated from the dorsum by a pale orange-gold (Maragrícola) to cream (Isla Gorgona) oblique lateral stripe with weakly defined margins, extending from the anterior part of the groin forward to the eye and onto the upper eyelid, and fading onto the canthus rostralis. In the Isla Gorgona population (Supplementary Fig. S8) this stripe is yellow to cream and is thicker than that of the Maragrícola frogs. In several of the Isla Gorgona frogs the stripe stops short of the eye, whereas in all Maragrícola frogs it reaches the eye and extends along the margin of the upper eyelid.

In all individuals a white to creamy stripe on the upper lip, distinctly paler than the oblique lateral stripe, extends from below the nostril to the axilla. In some it extends posteriorly as a poorly defined ventrolateral stripe (Supplementary Fig. S8, ANDES:A:3695). In some individuals from Isla Gorgona, the stripe is broken, but in all Maragrícola individuals it is complete.

The concealed surfaces of the groin are dark. In the Gorgona population, a somewhat diffuse, oval to elongate blotch with a yellow-white suffusion lies on the anterior concealed surface of the thigh, but posterior to the inguinal fold. A smaller and more elongate paracloacal mark is present on the posterior thigh surface. The yellow hue is much less pronounced than that of the Anchicayá population of *E.* sp. 1.

In the Maragrícola population (Supplementary Fig. S9) the blotch is more diffuse and lacks the yellow suffusion altogether, appearing as a dull gray region.

Ventral surfaces. In the Gorgona population the ventral surfaces of the throat, belly, thigh, and shank range from pale blue turquoise to nearly white, with irregular black blotches and vermiculations. The mottling ranges from 3-4 patches on a white background to a predominance of black pigment (Supplementary Fig. S8). Two irregular and elongate black blotches extend posteriorly from the tip of the chin but in the Maragrícola population these do not reach the pectoral region. This pattern is obscured or disrupted by diffuse dark gray pigment in the area of the vocal sac in males.

Almost all Maragrícola individuals have gray ventral limb surfaces that lack mottling. The ventral surfaces of the belly are similar to those of individuals from Isla Gorgona. As noted above, some individuals have a ventrolateral stripe.

#### 3.4.4. Epipedobates *aff*. espinosai

Images of *E.* aff*. espinosai* are from the Bilsa, Cube, Lita, La Nutria, and Pachijal populations (Figs. 1–2, node I; Supplementary Figs. S10-S13).

Dorsal surfaces. In all populations, the dorsal surfaces are dark brick red to dark brown with a hint of orange. The dorsal surfaces of the Bilsa frogs are slightly more orange than the other populations, possibly because the photos were taken under natural light rather than with a flash.

Lateral and concealed surfaces. The flanks and sides of the snout are solid black and smooth and are separated from the dorsum by a pale orange-cream oblique lateral stripe with weakly defined margins, extending from the anterior part of the groin forward to the eye. The populations differ in the anterior extent of the oblique lateral stripe. In Bilsa, Cube, and Lita populations, the stripe usually reaches the eye, becoming more diffuse and extending onto the canthus rostralis in a few individuals. In the La Nutria and Pachijal populations, the stripe almost reaches to the level of the scapula. In all populations the stripe color becomes white posteriorly as it reaches the groin region. Regardless of the extent of the stripe, a distinct and continuous white upper labial stripe is present, extending from the tip of the snout to the axilla. In some individuals in all populations it extends posterior to the axilla to merge with the ventrolateral stripe (Supplementary Fig. S11, QCAZ:A:58239). In one individual (Supplementary Fig. S13, QCAZ:A:58335), the most posterior edge of the ventrolateral stripe is yellow, as is the edge of the blotch posterior to the inguinal area.

The concealed surfaces of the groin are mostly black. In some populations, and in some individuals within populations, an oval to elongate whitish blotch is present on the anterior concealed surface of the thigh, but posterior to the inguinal fold. However, its presence and distinctness is highly variable both within and among populations. It is evident in only one individual in the Bilsa population, and ranges from absent to distinct in the Cube population (and is continuous with the oblique lateral stripe in some individuals). It is present and distinct in all frogs of the Lita population, and variably present in the Pachijal and La Nutria populations. A smaller and more elongate paracloacal mark blotch on the posterior thigh shows the same type of variation in the corresponding populations

Ventral surfaces. In all but the Bilsa population, the ventral surfaces of the throat, belly, thigh, and shank range from pale blue turquoise to nearly white, with irregular black blotches and vermiculations. In the Lita population, the venter of individuals from the same collection series varies from pale blue venter to white venter. The venter is highly variable in the Bilsa population, in which some frogs have no mottling or very diffuse mottling (Supplementary Fig. S10) while others have a predominantly black venter. As noted above, some individuals have a ventrolateral stripe. Additionally, in almost all individuals from Bilsa the pale regions of the venter are cream-colored; two frogs have a hint of blue. Interestingly, most of the frogs with largely pale venters are those that cluster with *E. machalilla* (Fig. 2).

Two irregular and elongate black blotches extend from the tip of the chin to the pectoral region, which are obscured by diffuse dark gray pigment in the area of the vocal sac in males. The variation of mottling on the chin corresponds to that of the venter.

#### 3.4.5. Epipedobates espinosai

Images of *E. espinosai* are represented by populations from La Perla, Mindo (a few kilometers from the type locality of *E. darwinwallacei*), Otongachi, Río Palenque, and Valle Hermoso (closest to the type locality) (Figs. 1–2, node J, Supplementary Figs. S14–S18). Individuals are highly variable in color and pattern, and include *E. darwinwallacei*, known as the Mindo morph. The populations from La Perla, Valle Hermoso, and Río Palenque are similar; all have a brick-red dorsum, with a hint of orange, and pale blue mottled venters. In contrast, the Mindo and Otongachi populations, which are geographically close, have black-brown dorsal surfaces with dark orange spots and blotches, and white or pale-yellow mottled venters.

Dorsal surfaces. Across all populations the dorsal surfaces are dark brick-red to dark brown with a hint of orange; the mottling on the dorsal surfaces ranges from barely perceptible to very distinct.

Lateral and concealed surfaces. The flanks and sides of the snout are solid black and smooth in all populations. The oblique lateral stripe is barely evident in the Río Palenque, La Perla, and Valle Hermoso populations, ranging from absent to reaching only about half the distance between the groin and eye (see Silverstone [1976:11] Figure 1 for an illustration of the paratype of *E. espinosai*). The stripe is white near the groin and becomes orange anteriorly as it fades out. In these populations the stripe is generally thinner than in *E.* aff. *espinosai*. In most frogs, the stripe does not reach as far into the inguinal region in *E. espinosai* as it does in *E.* aff. *espinosai*. In areas where the stripe is absent the dorsal color grades into the black flanks.

In almost all individuals the upper lip stripe does not reach the snout tip (Supplementary Fig. S14, QCAZ:A:52636) and in some the stripe is broken into elongate blotches or reduced to a few spots (Otongachi frogs). The stripe is wider in the Mindo and Otongachi frogs than in the other populations. In a very few individuals does it extend posteriorly as a broken series of blotches (Supplementary Fig. S15, QCAZ:A:58283). An elongate whitish to gray paracloacal mark is present on the posterior surface of the thigh in a few individuals in the Río Palenque and Valle Hermoso populations but absent in the La Perla frogs.

In the Mindo and Otongachi populations the oblique lateral stripe is wider and more prominent. It extends further anteriorly, reaching the axilla in the Mindo population, but is slightly shorter and paler in the Otongachi frogs, particularly at the posterior end. In a few individuals the stripe is broken into two or three elongate blotches. In the Mindo population it matches the color of the dorsal blotches. The paracloacal mark is dark orange in the Mindo and Otongachi frogs, distinct and more elongate than in the other populations, and in several frogs is accompanied by small spots of the same color and pattern as the dorsal markings.

Ventral surfaces. The ventral surfaces are pale blue with black mottling in most of the Río Palenque, La Perla, and Valle Hermoso frogs. Some Río Palenque frogs have pale gray rather than pale blue background color; this population is also extremely variable in the melanism of the ventral pattern. The ventral surfaces of the Mindo frogs have a pale yellow to yellow-orange background, but the black regions predominate in all specimens. Individuals from Otongachi are similar to Mindo but vary in creamy yellow to blue coloring with predominant black patches. A few individuals have a poorly defined and broken ventrolateral stripe.

As in *E.* aff. *espinosai*, two irregular and elongate black blotches extend from the tip of the chin to the pectoral region, which are obscured in males by dark gray pigment on the vocal sac. The mottling on the chin varies similarly to that of the venter and in some frogs the chin is almost entirely black.

#### 3.4.6. Epipedobates machalilla

Images of *E. machalilla* are represented by populations from 5 de Agosto, Jouneche, and Río Ayampe (near the type-locality; Figs. 1–2, nodes N and P; Supplementary Figs. S19-S21).

Dorsal surfaces. *Epipedobates machalilla* have a light brown dorsum with irregular darker X-shaped marks in the scapular region (as opposed to a solid brick-red dorsum in *E. espinosai, E.* aff. *espinosai*, and *E. boulengeri*) and similarly colored dark brown mottling across dorsal surfaces. In some frogs the “X” is fragmented. The dorsal surfaces of the limbs are similarly colored.

Lateral and concealed surfaces. The flanks and sides of the snout are solid black and smooth. The oblique lateral stripe cleanly separates the dorsal color from the flanks. It is distinct, complete, and extends posteriorly into the inguinal region further than in the other species. The stripe is goldish cream through its length, fading slightly near the eye. It extends to the eye but not onto the snout. The flanks and sides of the snout are solid dark brown to black and smooth in all populations.

An upper labial stripe is present in all individuals; it is wide, continuous, and reaches the tip of the snout. It also broadly contacts the lower margin of the eye. In all frogs the labial stripe continues posteriorly as a wide ventrolateral stripe located at the region of contact between the flank color and belly color. Although the belly of *E. machalilla* is yellowish white, the ventrolateral stripe is distinctly pigmented with white and is easily distinguished from the belly color (Supplementary Fig. S21, QCAZ:A:58296).

In the Río Ayampe frogs, the inguinal region is suffused with a bright orange marking that extends onto the proximal region of the thigh. It is also present on the posterior concealed surfaces of the thigh as a blotch or series of 2-3 blotches extending most of the length of the thigh, contrasting with a black-brown background. The orange color of the posterior thighs is continuous proximally with a paler paracloacal mark. The orange color is also present on the concealed surface of the shank. In some specimens a small orange patch is present in the axilla.

In contrast, no bright orange regions are present in the 5 de Agosto and Jouneche populations, although the anterior and posterior concealed thigh surfaces have a faint orange cast in some individuals. An indistinct paracloacal mark is present in some.

Ventral surfaces. The belly surfaces of all populations are immaculate, creamy white to yellowish white. The chin is yellow in females and yellowish gray in males and lacks the two irregular and elongate black blotches. The ventral surfaces of the limbs are tan to flesh-colored. No spots or mottling are present in any specimens. As noted above, some individuals have a ventrolateral stripe.

#### 3.4.7. Epipedobates tricolor

Images of *Epipedobates tricolor* (node Q) are represented by populations from Chazo Juan, Guanujo, and San José de Tambo (Figs. 1–2, node Q, Supplementary Figs. S22-S24). The first two populations are geographically proximate and are phenotypically very similar. San José de Tambo is further south, and the coloration is somewhat different, as described below.

Dorsal surfaces. In the Guanujo/Chazo Juan populations (known as the “Río” morph in the pet trade, these sites are sometimes also called Echeandia after the canton name), the background dorsal color is copper-brown with a hint of iridescence. Yellow to yellow-green mottling is present, in some cases covering half of the dorsal surfaces; in most individuals these light markings form a mid-dorsal stripe with irregular edges. In almost all frogs the dorsal surface of the snout is completely yellow-green.

In contrast, the dorsal surfaces of the San José de Tambo population (known as the “Cielito” morph in the pet trade) are dark brown with a reddish cast in some. Yellowish markings are present in almost all specimens except one in which the markings are copper to dark orange. The markings are either small elongate blotches or irregular spots and do not form a middorsal pattern in any frogs. The top of the snout is the same in color and pattern as the dorsum.

Lateral and concealed surfaces. The flanks and sides of the snout are black in the three populations. In the Chazo Juan/Guanujo populations the flanks are separated by a broad yellow or yellow-green oblique lateral stripe, much broader than in species such as *E. espinosai* or *E. boulengeri*. The stripe extends forward onto the eyelid and merges with the coloration of the snout. In the San José de Tambo populations the stripe is also broad and extends on to the eyelid, but in some individuals it is broken into a series of blotches. It is less yellow and is almost pale blue in a couple of specimens.

An upper lip stripe is present in all individuals; it is wide, and does not reach the tip of the snout in most frogs. It also contacts the lower margin of the eye. In contrast to some other species of *Epipedobates*, the lip stripe does not appear to form a more or less discrete ventrolateral stripe because the pale mottling of the venter intrudes into this area. In the San José de Tambo population, only one individual appears to have a hint of a ventrolateral stripe (Supplementary Fig. S24, QCAZ:A:53806).

The concealed surfaces of the inguinal region are bright orange to orange-red in the Chazo Juan/Guanujo frogs, and red in the San José de Tambo specimens. The size and shape of the blotch vary greatly within populations, but it is consistently present. Smaller blotches are evident in the posterior region of the axilla in all three populations and are pigmented like those of the inguinal region.

Similarly colored but elongate paracloacal marks are present on the posterior concealed surface of the thigh, and similar blotches occur on the concealed surface of the shank in most individuals of the Chazo Juan/Guanujo frogs, and in all individuals of the San José de Tambo specimens. In several of the Chazo Juan/Guanujo frogs the blotches are more diffuse and not as bright. In a few San José de Tambo specimens, a very small red blotch is present in the concealed surface of the ankle.

Ventral surfaces. The ventral surfaces of the throat and belly are brownish black and covered by extremely variable pale yellow (Guanujo and Chazo Juan) or pale blue (San Jose de Tambo) mottling. As in other species, the throat of males is grayish in the region of the vocal sac. In a very few individuals two irregular and elongate black blotches extend from the tip of the chin to the pectoral region. The ventral surfaces of the limbs are highly variable, from flesh or pale yellow with indistinct mottling (Guanujo) to yellow or dull orange (Chazo Juan) to black with pale blue mottling (San José de Tambo). As noted above, some individuals have a ventrolateral stripe.

#### 3.4.8. Epipedobates anthonyi

Images of *E. anthonyi* (node L) represent populations from Moromoro, Pasaje, and Uzchurummi (Figs. 1–2, node L; Supplementary Figs. S25-27). *Epipedobates anthonyi* expresses great variation in color, as evidenced by the numerous informal pet trade names for color morphs. Including the Peruvian populations, this species has the widest geographic and elevational distribution of any *Epipedobates* species (Fig. 1).

Dorsal surfaces. Frogs from our sample of only three populations have a brown to dark scarlet dorsum. All specimens have a mid-dorsal stripe with the same color (or greener color in the Pasaje frogs) as the oblique lateral stripe. The stripe varies in length, width, and continuity. In a few frogs, it extends from the tip of the snout to the cloaca (Supplementary Fig. S27, QCAZ:A:53697), but in some it is broken into a continuous series of short blotches (Supplementary Fig. S26, QCAZ:A:53679), while in others only 4–5 blotches are present (Supplementary Fig. S27, QCAZ:A:53695). In the Moromoro population the stripe is continuous and the broadest.

In some individuals of the Moromoro population, discrete blotches of the same color as the mid-dorsal and oblique lateral stripes occur on the dorsal surfaces of the hind limbs.

Lateral and concealed surfaces. The flanks and sides of the snout vary from dark reddish brown, matching the dorsum, to solid black. The oblique lateral stripe ranges from very pale blue to gray to pale yellow. Like the mid-dorsal stripe, it may be continuous (from the tip of snout, extending over the upper eyelid to inguinal region), broken into several blotches, or reduced to a very few blotches. In the Moromoro population the stripe is continuous and broad; in the other two it is narrower and discontinuous in some individuals.

The upper labial stripe varies in the same way as the oblique lateral stripe and mid-dorsal stripe. At its greatest extent it is wide, continuous, and reaches the tip of the snout, but it may be reduced to a series of small blotches; it broadly contacts the eye in some individuals. In most frogs the labial stripe continues posteriorly as a wide ventrolateral stripe located at the region of contact between the flank color and belly color. In some frogs the stripe is discontinuous and broken into blotches. In individuals in which the belly is whitish, the stripe nonetheless is distinctly pigmented with white (Supplementary Fig. S25, QCAZ:A:53688).

The concealed surfaces of the groin have a dark background similar to the flank color. In many individuals, an irregular brilliant red or red-orange blotch lies in the inguinal region and the anterior concealed surface of the thigh. The presence/absence, size, and shape of this marking vary greatly both within and among populations; in some, the blotch is divided into two parts. Inspection of images from the three populations does not indicate a difference among the populations in the degree of variation. All three have individuals with a large blotch and others with no blotch. We did not photograph both sides of the frog and so do not know whether the blotch characteristics have left-right symmetry.

In some individuals, the red or red-orange blotch extends posteriorly onto the anterior thigh surface. A smaller paracloacal mark is variably present on the posterior thigh surface and the proximal part of the concealed surface of the shank. We did not observe any individual with reddish orange markings in the axilla (as is present in some individuals of *E. tricolor*). In general, the size and extent of these flash markings are greater in *E. tricolor* than in *E. anthonyi*, at least for the populations we examined.

Ventral surfaces. The belly and throat are highly variable. In the Pasaje population the surfaces are brick-red with dull white mottling; the degree of mottling varies from complete to covering about one-third of the venter. The Uzchurummi population has reddish brown to chocolate ventral surfaces with dull gray to pale blue mottling (matching the mid-dorsal stripe) that is less defined than in the Pasaje frogs. In some, the whitish mottling is mostly absent. The ventral surfaces of the limbs vary from dull orange-brown to chocolate brown.

The Moromoro frogs display the greatest variation in ventral coloration and pattern. In a continuous field series of eight frogs (Supplementary Fig. S25), the venter varies from almost solid black (QCAZ:A:53689) to black with whitish mottling to whitish (QCAZ:A:5368). In some specimens (e.g., QCAZ:A:53686), the whitish belly appears to indicate the lack of the opaque white pigment that is seen in QCAZ:A:53687, and more closely resembles the belly of *E. machalilla*. As noted above, some individuals have a ventrolateral stripe.

In all three populations, males (with the exception of some Moromoro frogs) have diffuse darker pigment in the region of the vocal sac, but this varies from pale to dark gray to dull orange. In no individuals do two irregular and elongate blotches extend from the tip of the chin to the pectoral region.

### 3.5. Phenotypic variation in size and tarsal keel

#### 3.5.1 Snout-vent length

Females are significantly larger than males in *E.* sp. 1, *E. espinosai*, *E. machalilla*, and *E. tricolor* (Table 1, Fig. 7). Across species, female and male *E. tricolor* were larger than females and males of other species, while *E. machalilla* males and females tended to be smaller than other species. We found a statistically significant and positive correlation between SVL and elevation for females of all species pooled (S = 57296, p = 0.030, rho = 0.22), but not for males of all species pooled (t = 1.50, p = 0.14). In a larger model including males, females, and individuals without a determined sex (but excluding juveniles) pooled from all species, the association between SVL and elevation was also positive (S = 812726, p = 0.0015, rho = 0.22), indicating that regardless of species, *Epipedobates* frogs tend to be larger at higher elevations. When comparing mean SVL (by population for all species, rather than by individual) with elevation, we found a marginally significant, positive correlation between females and elevation (t = 2.17, p = 0.042), but not for males (t = 1.08, p = 0.30) or all data excluding juveniles (t = 1.59, p = 0.12). Finally, we note that the Isla Gorgona population of *E. boulengeri* s. s. was statistically larger than its phylogenetically closest mainland population in Maragrícola (Welch two-sample t-test: t = 4.13, df = 13, p = 0.0012).

**Figure 7.**
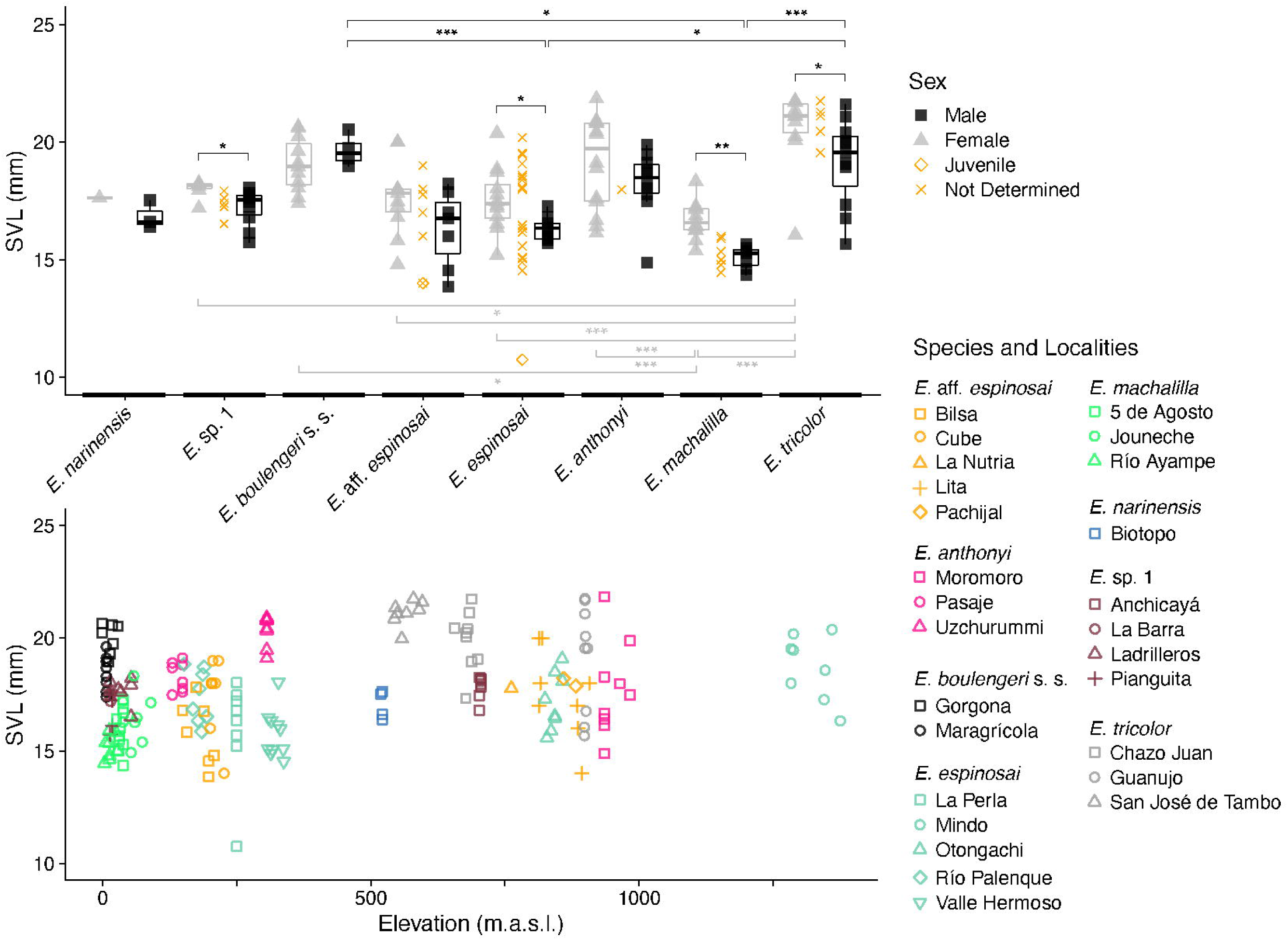
(Top) Snout-vent length (SVL) varies significantly within and among some of *Epipedobates* species. Significant comparisons are denoted with asterisks: *, p<0.05; **, p <0.001; ***, p <0.0001. See Table 1 for statistical test details. (Bottom) SVL increases with elevation across the genus.

**Table 1.** Statistical comparisons of snout-vent length (SVL) within and among species, also shown in Figure 7. Upper triangle values are comparisons of males; lower triangle values are comparisons of females; the diagonal shows within-species comparisons between the sexes and sample sizes for each sex. See Materials and Methods for a description of the parametric and non-parametric approaches. N_M_, number of males; N_F_, number of females; df, degrees of freedom; t, t-test statistic; W, Wilcoxon rank-sum test statistic; p_adj_, adjusted p-value.

|  | *E. narinensis* | *E. sp. 1* | *E. boulengeri*<br>s. s. | *E. aff. espinosai* | *E. espinosai* | *E. anthonyi* | *E. machalilla* | *E. tricolor* |
| --- | --- | --- | --- | --- | --- | --- | --- | --- |
| *E. narinensis* | $N_M = 3$<br>$N_F = 1$<br>N/A | $Z = 0.41$<br>$p_{adj} = 1$ | $Z = 1.99$<br>$p_{adj} = 0.79$ | $Z = -0.30$<br>$p_{adj} = 1$ | $Z = -0.60$<br>$p_{adj} = 1$ | $Z = 1.28$<br>$p_{adj} = 1$ | $Z = -1.59$<br>$p_{adj} = 1$ | $Z = 1.74$<br>$p_{adj} = 1$ |
| *E. sp. 1* | $t = 0.39$<br>$p_{adj} = 1$ | $N_M = 10$<br>$N_F = 7$<br>$W = 62$<br>$p = 0.0068$ | $Z = 2.12$<br>$p_{adj} = 0.65$ | $Z = -0.96$<br>$p_{adj} = 1$ | $Z = -1.39$<br>$p_{adj} = 1$ | $Z = 1.28$<br>$p_{adj} = 1$ | $Z = -2.76$<br>$p_{adj} = 0.13$ | $Z = 1.98$<br>$p_{adj} = 0.79$ |
| *E. boulengeri*<br>s. s. | $t = 1.39$<br>$p_{adj} = 0.98$ | $t = 1.0$<br>$p_{adj} = 0.82$ | $N_M = 4$<br>$N_F = 11$<br>$t = -1.28$<br>$df = 9.41$<br>$p = 0.23$ | $Z = -2.75$<br>$p_{adj} = 0.13$ | $Z = -3.15$<br>$p_{adj} = 0.039$ | $Z = -1.15$<br>$p_{adj} = 1$ | $Z = -4.17$<br>$p_{adj} = 0.00081$ | $Z = -0.66$<br>$p_{adj} = 1$ |
| *E. aff. espinosai* | $t = 0.0046$<br>$p_{adj} = 1$ | $t = -0.38$<br>$p_{adj} = 1$ | $t = -1.38$<br>$p_{adj} = 0.31$ | $N_M = 7$<br>$N_F = 11$<br>$t = 1.70$<br>$df = 12.34$<br>$p = 0.11$ | $Z = -0.33$<br>$p_{adj} = 1$ | $Z = 2.12$<br>$p_{adj} = 0.65$ | $Z = -1.66$<br>$p_{adj} = 1$ | $Z = 2.77$<br>$p_{adj} = 0.13$ |
| *E. espinosai* | $t = -0.79$<br>$p_{adj} = 1$ | $t = -0.47$<br>$p_{adj} = 1$ | $t = -1.5$<br>$p_{adj} = 0.22$ | $t = -0.84$<br>$p_{adj} = 1$ | $N_M = 9$<br>$N_F = 12$<br>$t = 2.94$<br>$df = 14.87$<br>$p = 0.010$ | $Z = 2.64$<br>$p_{adj} = 0.17$ | $Z = -1.43$<br>$p_{adj} = 1$ | $Z = 3.35$<br>$p_{adj} = 0.020$ |
| *E. anthonyi* | $t = 1.53$<br>$p_{adj} = 0.97$ | $t = 1.15$<br>$p_{adj} = 0.66$ | $t = 0.15$<br>$p_{adj} = 1$ | $t = 1.53$<br>$p_{adj} = 0.15$ | $t = 1.61$<br>$p_{adj} = 0.089$ | $N_M = 10$<br>$N_F = 14$<br>$t = 1.34$<br>$df = 22.00$<br>$p = 0.19$ | $Z = -3.92$<br>$p_{adj} = 0.0023$ | $Z = 0.68$<br>$p_{adj} = 1$ |
| *E. machalilla* | $t = -0.96$<br>$p_{adj} = 1$ | $t = -1.34$<br>$p_{adj} = 0.49$ | $t = -2.34$<br>$p_{adj} = 0.0041$ | $t = -0.96$<br>$p_{adj} = 0.73$ | $t = -0.88$<br>$p_{adj} = 0.80$ | $t = -2.59$<br>$p_{adj} = 0.00073$ | $N_M = 7$<br>$N_F = 12$<br>$t = 5.50$<br>$df = 16.81$<br>$p = 0.000040$ | $Z = 4.61$<br>$p_{adj} = 0.00013$ |
| *E. tricolor* | $t = 3.0$<br>$p_{adj} = 0.49$ | $t = 2.58$<br>$p_{adj} = 0.0094$ | $t = 1.58$<br>$p_{adj} = 0.19$ | $t = 2.97$<br>$p_{adj} = 0.00023$ | $t = 3.05$<br>$p_{adj} = 0.000094$ | $t = 1.44$<br>$p_{adj} = 0.23$ | $t = 3.93$<br>$p_{adj} = 0.0000003$<br>0 | $N_M = 11$<br>$N_F = 10$<br>$W = 88.5$<br>$p = 0.020$ |

#### 3.5.2. Tarsal keel

Given that *E. darwinwallacei* was described as a new species based on color and tarsal keel morphology, and we found genetic evidence that it is not a distinct genetic lineage, we reviewed tarsal keel morphology in several species of the *Epipedobates* genus. In the diagnosis of *E. darwinwallacei*, Cisneros-Heredia and Yánez-Muñoz (2010) characterized the tarsal keel as displaying two conditions: “tarsal keel straight or weakly curved extending proximolateral from preaxial edge of inner metatarsal tubercle or short, tubercle-like, transversely across tarsus, not extending from metatarsal tubercle.” This wording exactly matches states 0 and 2 of Grant et al. (2006). However, it is not entirely clear whether they intended to reflect true intraspecific variation (some individuals have state 0 and others have state 2) or uncertainty in the characterization of the tarsal keel shape as either state 0 or 2. Grant et al. (2017) scored *E. darwinwallacei* as 0&2, identical to Cisneros-Heredia and Yánez-Muñoz (2010).

Using the state definitions of Grant et al. (2006), we scored *E. anthonyi*, *E. tricolor*, and *E. machalilla* as state 2, *E. espinosai* from Río Palenque as state 0, *E. espinosai* from Vía Chone (near the type locality of *E. espinosai*) as 2, *E. “darwinwallacei”* specimens from Mindo as 0 and 2 (different specimens). In general, we found it difficult to apply the criteria for states 0 and 2 consistently in the specimens we examined.

Thus, our data, although meager, indicate that specimens from near the type locality of *E. espinosai* and specimens from near Mindo (considered to be *E. darwinwallacei*) show intraspecific variation, and that the two groups cannot be distinguished based on this character.

However, this variation is apparently due to the difficulty of assessing state in specimens with different conditions of preservation, and not necessarily to true polymorphism of states within a species.

Cisneros-Heredia and Yánez-Muñoz (2010) characterized the tarsal keel of other *Epipedobates* (*anthonyi, tricolor, machalilla, boulengeri, narinensis*, and *espinosai*) as “tarsal keel large and strongly curved,” which corresponds roughly to state 1 of Grant et al. (2006). However, this contrasts with Grant et al. (2006, 2017), who scored *E. anthonyi*, *E. boulengeri* s. s., *E. “boulengeri,” E. “espinosai,” E. machalilla*, and *E. tricolor* as state 2 rather than state 1. It also disagrees with our characterization of the states for *E. anthonyi, E. tricolor*, and *E. machalilla*. We discuss this problem further in section 4.5.5 on *Epipedobates espinosai*.

### 3.6. Species distributions

Based on the specimens included in this study, the ranges of *Epipedobates* species appear to be allopatric except for *E. narinensis* and *E.* aff. *espinosai* (Fig. 1; but see section 4.1.3 below for more information regarding recent field work and the prevalence of parapatry). In terms of elevation, each species tends to occupy a lowland distribution (e.g., *E. boulengeri* s. s.: 0–156 masl; *E.* sp. 1: 0–700 masl; *E. machalilla*: 0–400 masl), an Andean mountain foothill range (*E. espinosai*: 152–1375; *E.* aff. *espinosai*: 87–907 masl; *E. narinensis*: ∼530 masl), or an extensive lowland-to-highland distribution, as in *E. anthonyi* (50–2100 masl) and *E. tricolor* (83–1800 masl) (Pontificia Universidad Católica del Ecuador, 2021).

## 4. Discussion

Herein we describe the genetic and phenotypic diversity of 29 populations of frogs within the dendrobatid genus *Epipedobates*. With an integrative taxonomic approach (Padial and de la Riva 2010), we use multiple lines of evidence to propose species boundaries under the general lineage species concept (de Queiroz 2005, 2007). We compare these to existing taxonomy and proposed species limits obtained from two objective species delimitation approaches (ASAP and GMYC). We recognize that the delimitation of species may in some cases be based on incomplete information and that future research may provide additional lines of evidence that support or reject our proposed species limits.

### 4.1. Phylogenetic patterns

#### 4.1.1. Phylogenetic analyses

We found seven strongly supported molecular clades that we associate with the following species (Fig. 2): *Epipedobates narinensis* (node C), *E.* sp. 1 (node D), *E. boulengeri* s. s. (node F), *E. espinosai* (node J), *E.* aff. *espinosai* (node I), *E. anthonyi* (node L), and *E. tricolor* (node Q). *Epipedobates machalilla* (nodes N and P) is not monophyletic in our molecular tree or that of Tarvin et al. (2017), but in neither is monophyly strongly rejected. The same six *E. boulengeri* (now *E.* aff. *espinosai*) individuals from the “Bilsa B” that clustered with *E. tricolor* in Tarvin et al. (2017a) cluster with individuals of *E. machalilla* from 5 de Agosto. However, that node (node P) has a posterior probability of <0.50 in the Bayesian tree (Fig. 2), so the topological conflict is not meaningful.

Tarvin et al. (2017a: Fig. 2) analyzed fewer individuals and populations of *Epipedobates* “*boulengeri*” and found that individuals assigned to the species represented several distinct lineages. The addition of many new sequences of *E.* “*boulengeri*” individuals to this study did not alter this pattern. Thus, in section 4.5 we propose taxonomic changes based on the previously unrecognized genetic diversity we identified.

The clade comprising *E. anthonyi, E. machalilla,* and *E. tricolor* (Fig. 2; node K) showed high support but lacked fine-scale resolution within each species. The phylogenetic relationship among species was similar to previous studies (Clough and Summers, 2000; Graham et al., 2004; Grant et al., 2006; Santos et al., 2003; Santos and Cannatella, 2011; Vences et al., 2003; Tarvin et al., 2017).

The times of divergence indicate a clade containing a cluster of species that arose in the last 3 Myrs. *Epipedobates narinensis* and *E.* sp. 1 diverged 2.78 Ma, *E. espinosai* and *E*. aff. *espinosai* diverged 2.53 Ma. *Epipedobates anthonyi* and *E. tricolor* diverged 1.99 Ma, and *E. machalilla* and *E. tricolor* diverged 1.51 Ma. Interestingly, the mainland and island populations of *E. boulengeri* s. s. diverged 3.06 Ma (Supplementary Fig. S2).

Using a large phylogeny based on mtDNA, Santos et al. (2009) estimated the age of dendrobatid clades. They found the age of the MRCA of *Silverstoneia* + *Epipedobates* (node 295 in their Supplementary Table S8 and Supplementary Fig. 5B) to be 15.9–16.9 Myrs (depending on the set of calibration constraints used). This accords well with our estimate of 13.91 Myrs (Supplementary Fig. S2). They also estimated the MRCA of their sample of *Epipedobates* species (their node 265, which excludes *E. narinensis* and *E.* sp. 1), occurs at 5.5–5.8 Ma, whereas our estimate for the same node (E in our Fig. 2) is 6.02 Ma. Thus, their estimated divergence times are remarkably congruent with ours, which were estimated using a secondary calibration of the root of the tree based on nuclear genes. However, our contribution of new genetic data of *E. narinensis* and *E.* sp. 1 extend the age of the *Epipedobates* clade by nearly 4 Myrs.

#### 4.1.2. Quantitative species delimitation

Results from two programs that estimate species limits did not provide high confidence in a specific number of species, nor in a set of groupings that were concordant with our proposed species limits based on monophyly and morphological differences. In ASAP, the two most highly supported partitions contained 2 or 56 groups. Similarly, the top GMYC result indicates six species (Fig. 3), but the 95% confidence region of 2–16 species indicates that confidence in the GMYC estimate of six species is low. Further, the likelihood ratio test of the null and alternative model in the GMYC analysis was indecisive (p = 0.05670), meaning there is equivocal evidence for a speciation model in our tree. When taking into account the effect of phylogenetic uncertainty on GMYC analyses using a posterior distribution of trees, the results are similar: about one-third of the trees supported recognition of five species and one-fifth supported six; the 95% confidence interval for the 100 samples is 3–8 species. Roughly 40% of the analyses yield p-values ≤ 0.05, again indicating poor support for any specific species delimitation scheme.

Although it is possible that the *Epipedobates* clade is composed of two or six species as these analyses might suggest (even weakly), after our review of a broader set of characters that are typically used for species delimitation in frog systematics, we propose that there are seven or eight species in *Epipedobates*. The top ASAP result, based on genetic clustering, seems to reflect only the relatively deep genetic divergence between the clade composed of *E. narinensis* and *E.* sp. 1 and other clades within the genus. The GMYC analysis, which considers branching patterns in a single-locus tree, suggest sets of species groupings that are closer but still distinct from those that we propose (Fig. 3). For example, we considered *E. narinensis* and *E*. sp. 1 to be distinct species (p-distance = 0.0535), whereas the GMYC analysis recovered them as one species.

Although the GMYC analysis revealed *E.* aff. *espinosai* and *E. espinosai* to be distinct species, as we did, it lumped *E. machalilla, E. tricolor,* and *E. anthonyi* into one species, whereas we recognize three. These species are phenotypically differentiated and easy to distinguish in the field, whereas their genetic distances are low, e.g., 3% between *E. anthonyi* and the other two.

Because the ASAP and GMYC analyses were indecisive as to the number of species, and because we could not decisively reject the coalescent model in GMYC, we conclude that the application of these methods to our dataset did not yield strong support for any specific partition scheme, including our proposal of eight species. A possible reason that our proposal is not a highly supported outcome of ASAP or GMYC analyses is that our dataset included several species that are weakly diverged. Various studies (e.g., Esselstyn et al. [2012] on bats, Talavera et al. [2013] on butterflies, Michonneau [2015] on holothurians) have examined the performance of GMYC by comparing clock models, tree models (Yule vs. coalescent), and the effects of identical haplotypes. We have not compared our results to these other studies, but in general we note these papers analyzed many more putative species than we did.

Although these quantitative species delimitation programs can be useful to produce a starting point for taxonomic revisions, in our case we found the results unconvincing.

Nevertheless, we highlight that additional data, especially of contact zones, may be necessary to verify the proposed species boundaries in *Epipedobates*.

#### 4.1.3. Possible historical and ongoing gene flow among Epipedobates species

Like Tarvin et al. (2017a), we found evidence of historical introgression between *E.* aff. *espinosai* (individuals previously assigned to *E. boulengeri* at Bilsa and La Tortuga sites) and *E*. *machalilla* based on the placement of individuals of the Bilsa population in disparate places of the phylogeny (see Fig. 2 in their paper). Tarvin et al. (2017) also found that *E. machalilla* is paraphyletic with respect to “*E. boulengeri* Bilsa B” (*E.* aff. *espinosai* in the present analysis) and *E. tricolor*. However, the bootstrap support for this relationship is <50. Given that all Bilsa individuals were collected on the same day and place by RDT, we are confident of the accuracy of the locality and correct association of sequences. This anomalous clustering is most readily explained by historical gene flow between the two species. In both studies, the disparate placement of individuals of *E.* aff. *espinosai* in two parts of the tree may be caused by *E. machalilla* haplotypes influencing the phylogenetic placement of some *E*. aff. *espinosai* (Fig. 5AB).

Additional fieldwork (Tarvin, Ron, pers. obs.) and specimen data (Pontificia Universidad Católica del Ecuador, 2021) show that most *Epipedobates* species pairs with adjacent distributions have ranges that overlap in contact zones, i.e., most species have parapatric distributions. In particular, *E. machalilla* is flanked by other *Epipedobates* species on all land-based edges of its distribution, which may have led to the multiple signatures of gene flow in this species. In addition to historical gene flow between *E. machalilla* and *E.* aff. *espinosai,* our phylogenetic reconstructions showed that *E. machalilla* individuals form a group that is paraphyletic to *E. tricolor*. In the *K_V_1.3* haplotype network (Fig. 5F), *E. machalilla* is split into two groups, one of which is closely related to *E. tricolor*, and the other of which is closely related to *E. anthonyi.* In addition, *E. tricolor* and *E. anthonyi* share a *K_V_1.3* haplotype. These patterns suggest historical and possibly ongoing gene flow within the subclade. Nevertheless, in the faster evolving mtDNA genes, none of these species share haplotypes, and our phenotypic data indicate substantial differentiation in color, advertisement call, and body size. Thus, we do not think that the genetic data alone warrant lumping these three species into a single unit at this time.

In amphibians, the speciation process has been suggested to be slow because reproductive incompatibilities tend to accumulate through many small-effect mutations rather than few large-effect ones (Dufresnes et al., 2021). It is plausible that some *Epipedobates* species such as *E. tricolor, E. machalilla,* and *E. anthonyi* are incipient species or subspecies that have not yet evolved complete reproductive isolation and remain “incompletely separated lineages” (de Queiroz, 2020); the alternative hypothesis is that these populations represent a single species with broad geographic variation in phenotype. Further assessment of gene flow and reproductive incompatibilities will be necessary to clarify species boundaries in these and other species of the *Epipedobates* genus. Recent advances in species delimitation methods have proposed comprehensive hybrid zone sampling as a step required for proper species delimitation (Chambers and Hillis, 2020; Hillis et al., 2021). Accordingly, we are investigating the possibility of more widespread and ongoing gene flow among putative *Epipedobates* species with more comprehensive genomic sampling of contact zones using ddRADseq.

### 4.2 Genetic variation

#### 4.2.1. Genetic distances

Our study confirms the low genetic divergence in the *12S–16S* among Ecuadorian *Epipedobates* species found by others (Clough and Summers, 2000; Santos et al., 2009; Santos and Cannatella, 2011; Tarvin et al., 2017a; Vences et al., 2003). Inclusion of Colombian populations (*E. narinensis* and *E*. sp. 1) revealed a deeper level of genetic divergence between *Epipedobates* species (> 7.0%) in the *12S–16S* sequences. These distances are noteworthy because phenotypically similar species such as *E. boulengeri* s. s. and *E.* sp. 1 show deep divergence (6.82%, Fig. 3), whereas phenotypically divergent species such as *E. machalilla* and *E. tricolor* show very low levels of divergence (1.04%, Fig. 3).

Researchers frequently use the level of sequence divergence in the *16S* gene to help guide species delimitation. In an analysis of *16S* variation in Malagasy frogs, Vences et al. (2005a: Fig. 5) found that uncorrected p-distances in the range of 2–6% overlapped between intraspecific and interspecific pairs. Vences et al. (2005b) proposed a “tentative” barcoding threshold of 5% in the *16S* fragment. Fouquet et al. (2007) narrowed the criterion, proposing that an uncorrected p-distance of 3% (in the same *16S* gene fragment) be used to identify candidate species, but not necessarily for formal species delimitation, which they suggested might include an evaluation of reproductive isolation and incomplete lineage sorting to avoid erroneous species descriptions.

The *12S–16S* fragment that we sequenced overlaps slightly with the more commonly used *16S* fragment. Because the uncorrected distances for the same pairs of species are almost identical between these two fragments, we can extend results by other authors to our data (Fig. 3). The smallest distances are 0.925% between *E. anthonyi* and *E. machalilla* and 1.04% between *E. tricolor* and *E. machalilla*. This is reflected in the haplotype networks (Fig. 5), in which *E. machalilla* haplotypes do not form a contiguous network. Based on the genetic distance criteria proposed by Vences et al. (2005b) and Fouquet et al. (2007), *E. machalilla* would be considered conspecific with *E. anthonyi* and *E. tricolor*. However, the three are phenotypically distinct in vocalization (Santos and Cannatella, 2011; Tarvin et al. 2017a) and color pattern throughout their ranges. Similarly, the genetic distance between *E.* sp. 1 and *E. narinensis* is only 1.75%, and yet the species are remarkably distinct phenotypically (Fig. 1); *E. narinensis* is greenish with little or no lateral stripe, whereas *E*. sp. 1 is very similar to *E. boulengeri*, yet the genetic distance between them is 7.4%. Overall, while some *Epipedobates* species retain the ancestral morphology of the genus, several species have experienced recent changes in morphology that created high phenotypic diversity within closely related lineages (i.e., *E. machalilla*, *E. tricolor*, and *E. anthonyi*; and *E. narinensis* and *E*. sp. 1). While other authors may consider that these low genetic distances indicate the presence of fewer species in the *Epipedobates* genus than we propose here, we identify distinct color patterns, acoustic signals, and evolutionary histories that are associated with each putative species. We recognize that future assessments may favor further lumping or splitting than what we propose here.

#### 4.2.2. Haplotype networks

The haplotype networks reinforce the conclusions based on genetic distance. The control region (Fig. 5B), which is well known to evolve quickly (Meyer, 1993), shows that only two individuals share a haplotype, yet the two most slowly evolving nuclear genes (*BMP2* and *H3*) have few haplotypes that are each shared by several individuals (Supplementary Table S3). The numerous equally parsimonious networks in the *K_V_1.3* network, indicated by dashed lines, indicate a lack of resolution among haplotype relationships, most likely due to the large number of heterozygotes in the *K_V_1.3* sequences.

There is some concordance between nDNA and mtDNA markers in the patterns of shared haplotypes of each species, especially for *K_V_1.3*. Although *BMP2* and *H3* are not very informative, *E. boulengeri* s. s. and *E. narinensis* do not share *BMP2* haplotypes with other species, supporting their genetic distinctiveness. In both nDNA and mtNDA genes, haplotype groups for each species are often interspersed with haplotypes of other species, e.g. compare *E. machalilla* and *E.* aff. *espinosai* or *E. anthonyi* in Fig. 5BD. In some cases, such patterns may have arisen because of equally parsimonious solutions in the network estimation (e.g., review grey dotted lines in Fig. 5C). Alternatively, these patterns could reflect the recent divergence among species and populations in this clade.

### 4.3. Phenotypic variation

#### 4.3.1 Patterns of conservative phenotypes and divergent aposematic coloration

We observed two general patterns in phenotypic variation: 1) inconspicuous species with low phenotypic variation but high genetic divergence in northern clades (nodes B, F, H); 2) two brightly patterned species with great phenotypic variation but low genetic divergence (*E. tricolor* and *E. anthonyi*) in the southern clades. Based on the phylogeny, the first pattern is explained by the plesiomorphic retention of an ancestral phenotype (e.g., cream oblique lateral stripe, brown-red dorsum, lack of flash colors in the inguinal region) in several species (*E*. sp. 1, *E*. *boulengeri* s. s., *E*. *espinosai*, and *E*. aff. *espinosai*). *Epipedobates narinensis* is exceptional in its green coloration.

With regards to the second pattern, many other aposematic poison frog species are polytypic, with distinct color patterns that vary widely among populations (Grant et al., 2017; Medina et al., 2013; Przeczek et al., 2008; Rojas, 2016; Roland et al., 2017; Wang, 2013). The most famous is *Oophaga pumilio*, with extreme pattern and color variation among island populations. This variation seems related to mating success because females show preference for males with the brightest dorsum (Maan and Cummings, 2009), and tadpoles imprint on their mother’s color (Yang et al. 2019).

In Dendrobatidae, conspicuousness is associated with body size (Santos and Cannatella 2011). In *Epipedobates*, we find that in general body size increases with elevation. Simultaneously, populations within a species tend to have brighter phenotypes at higher elevations. For example, the two known populations of *E. espinosai* with bright dorsal marks, Pucayacu and Mindo, are highland populations (900 and 1375 masl, respectively). This elevation pattern is similar to that reported in *E. anthonyi* in which highland populations are more brightly colored than lowland populations (Páez-Vacas et al. 2021).

Three of the eight species analyzed here (*E.* sp. 1, *E. espinosai*, and *E.* aff. *espinosai*) have been identified as *E. boulengeri* in the literature, indicating that species identification was difficult. Individuals of these three species generally match the phenotype of *E. boulengeri* s. s., with a brown to brownish red dorsum, at times with some degree of red-orange mottling (the prominent mottling of the Mindo frogs is an outlier), an oblique lateral line separating the dorsum from black-brown flanks, and a marbled belly. Despite the morphological resemblance to three other species, *E. boulengeri* s. s. is genetically very divergent. It shows 3.42% divergence from *E. espinosai* and *E.* aff. *espinosai* and 7.37% from *E. narinensis* in the *12S–16S* gene fragment (Table 1).

*Epipedobates boulengeri* s. s. also differs from *E. espinosai* and *E.* aff. *espinosai* in the following phenotypic traits. *Epipedobates espinosai* has a dark red dorsum with small indistinct black blotches mixed with yellow, orange, or white blotches, and *E.* aff. *espinosai* has a generally uniform brown dorsum (Fig. 5), while *E. boulengeri* s. s. has a dark brown dorsum. The oblique lateral stripe is complete and present in all populations of *E. boulengeri* s. s. that we reviewed; in *E. espinosai* the oblique lateral stripe is incomplete in all populations. The presence and completeness of the oblique lateral stripe in *E.* aff. *espinosai* varies depending on the population: Bilsa, Cube and Lita have a complete oblique lateral stripe, but in other populations it is incomplete. Previously, the distribution of *E. boulengeri* was thought to encompass much of central-western Ecuador to central-western Colombia (Cisneros-Heredia and Kahn, 2016); however, we found that it is restricted to a narrow corridor along the coast from southwest Colombia (Isla Gorgona) to the northwest corner of Ecuador (Fig. 1), but more extensive sampling is needed, especially in southwestern Colombia.

#### 4.3.2 Color variation among the Colombian species

*Epipedobates narinensis* and *E.* sp. 1 share a consistent feature: the presence of an elongated yellow blotch originating on the anterior and posterior edges of the inguinal area, extending to the external side of the thighs and forward on the flanks. These yellow blotches are different from the flash colors in *E. machalilla, E. tricolor,* and *E. anthonyi*. The original description of *E. boulengeri* (Barbour, 1909) did not mention yellow blotches on the legs; however, Silverstone (1976:28) noted the yellow region in populations of *E.* “*boulengeri*” from La Guayacana, Colombia:

> “In life, in specimens from La Guayacana, Colombia (my notes), the iris was black. The dorsum was brown. The side of the body was black with a complete yellow lateral stripe, The upper lip and the proximodorsal surface of the upper arm were white. The dorsal surface of the thigh was black with yellow spots or two yellow stripes. The digits were banded brown and white. The venter and the ventral surface of the limbs were black with white marbling.”

La Guayacana is about 12 km west of the Biotopo Reserve, the type-locality of *E. narinensis*, on Highway 10. The color differences between the Biotopo and La Guayancana specimens suggest that greater assessment of *Epipedobates* in southwestern Colombia is needed.

Additionally, *E.* aff. *espinosai* is represented in Colombia by the La Nutria population, a few kilometers east of Reserva Biotopo. Given that *E.* aff. *espinosai* lacks the yellow spots (Supplementary Figs. S10-S13) that Silverstone (1976) described in the La Guayacana frogs, it is possible that there are three species in close proximity (Fig. 1): an unknown species at La Guayacana (possibly *E.* sp. 1), *E. narinensis* at Biotopo, and *E.* aff. *espinosai* at La Nutria. However, we found only one *E.* aff. *espinosai* individual at La Nutria, so there may be unsampled phenotypic diversity of *E.* aff. *espinosai* in this region, possibly with yellow markings as described by Silverstone (1976).

Silverstone (1976: 28) also noted: “In specimens from the lower Rio Calima, Colombia (I. Cabrera, notes at USNM), the lateral stripe and the dorsal thigh markings were red. The venter was greenish or white with black marbling.” Based on locality, these frogs might be referrable to *E.* sp. 1 or *E.* aff. *espinosai*; however, *E*. sp. 1 differs in having yellow to orange markings and a gray to pale blue venter and *E.* aff. *espinosai* differs in having white to orange stripes and white, yellow, or blueish venters.

Based on color pattern and locality, the photos in Lötters et al. (2007) are likely assignable to these species: Fig. 514, *E. boulengeri* s. s. (Gorgona Island); Figs. 515–516, *E.* aff. *espinosai* (Lita, Esmeraldas); Figs. 517–518, *E.* aff. *espinosai* (Esmeraldas); Figs. 519–520, *E. espinosai* (Santo Domingo, Pichincha).

#### 4.3.3 Northern Ecuadorian species

The northern Ecuadorian species include *E.* aff. *espinosai* and *E. espinosai*, although *E. boulengeri* s. s. is also found in the northwestern corner of Ecuador. Some populations of *E. espinosai* and *E.* aff. *espinosai* are extremely similar in color pattern, whereas others such as the *E. espinosai* Mindo and Pucayacu morphs are easily distinguished from *E*. aff. *espinosai*. However, the labial stripe is broken or reduced to a series of blotches in *E.* aff. *espinosai*, whereas it is continuous and reaches almost to the tip of the snout in *E. espinosai*.

#### 4.3.4 South-central Ecuadorian species

The populations with the greatest phenotypic variation in color are the aposematic species *E. tricolor* and *E. anthonyi*. Although the former has a relatively restricted distribution, it nonetheless shows broad phenotypic variation (Tarvin et al. 2017a; Figs. S22-S24). We did not attempt to document the full extent of phenotypic variation in *E. anthonyi* and *E. tricolor.* Tarvin et al. (2017a) compared genetic and phenotypic dissimilarity within and among the southern Ecuadorian species, showing that phenotypic diversity was greater than genetic diversity in these species. They also showed that the introgressed individuals of the *E. boulengeri* Bilsa population (now referred to *E.* aff. *espinosai*) were phenotypically similar to *E. machalilla*. Our expanded sampling of the *E. boulengeri* complex presented herein greatly expands the known genetic diversity of the genus and provides evidence for deeply diverged inconspicuous species (e.g., *E. narinensis* and *E. boulengeri* s. s.) in addition to shallowly diverged species with color polytypism (e.g., *E. tricolor* and *E. anthonyi*).

### 4.4. Species distributions and historical biogeography

No localities have sympatric species of *Epipedobates*, although different species occur at some geographically proximate localities (e.g., Biotopo with *E. narinensis* and La Nutria with E. aff. *espinosai*). These species pairs may have contact zones with ongoing or historical gene flow, a hypothesis supported by the apparent historical mitochondrial introgression between *E. machalilla* and *E.* aff. *espinosai* (Fig. 1, Tarvin et al. 2017a).

The oldest node in our phylogeny separates the clade containing *E. narinensis, E.* sp. 1, and *E. boulengeri* s. s., which are found in Colombia and far north Ecuador, from the clade containing the youngest nodes that lead to species distributed in southern Ecuador and northern Peru (*E. machalilla, E. tricolor,* and *E. anthonyi*). The species with the northernmost distribution is *E.* sp. 1 from Valle del Cauca (Colombia) near the Pacific coast up to 700 masl. In contrast, the southernmost species is *E. anthonyi* at Canchaque, Piura, Peru (Duellman and Wild 1993). The great circle distance between the distribution extremes of *Epipedobates* is 1,100 km, which includes a diversity of forest types and habitats that likely emerged from a complex geological history of northwestern South America. Our phylogenetic reconstruction suggests a pattern of intermittent speciation during southward dispersal from Colombia. *Silverstoneia*, the sister-group of *Epipedobates*, includes eight species found in Costa Rica, Panama, and the Chocó region of Colombia, but it is not known from Ecuador. This distributional pattern is consistent with a southward dispersal and/or cladogenesis from Central America into the lowland rainforests of the Chocó, trans-Andean Ecuador, and dry forests of southern Ecuador and northern Peru.

The estimated age of the crown clade *Epipedobates* is 11.1 Mya and corresponds to the late Miocene divergence between *Epipedobates* sp. 1 + *E. narinensis* and the remaining species. During the Miocene and Pliocene, the Chocó region and Northern Peru experienced cyclical sea-level fluctuations producing repeated marine incursions (Duque-Caro 1990; Borrero et al. 2012; Pérez-Escobar et al. 2019) and likely isolation of remnant islands that were later reconnected.

At its northernmost extent, *Epipedobates* includes *E.* sp. 1 and *E. narinensis* as sister species that shared a most recent common ancestor (MRCA) at 2.78 Mya. *Epipedobates* sp. 1 occurs near Buenaventura, Valle del Cauca (Colombia) and includes populations from La Barra, Ladrilleros and Anchicayá from near the Pacific coast up to 700 masl (at Anchicayá). *Epipedobates narinensis* has a narrow distribution around Biotopo, Nariño (Colombia) at ∼520 masl. The great circle distance between *Epipedobates* sp. 1 and *E. narinensis* is ∼280 km, which suggests that more populations and possibly more species might exist along this latitudinal and altitudinal gradient. To date, limited biodiversity surveys have been conducted in this region of the Colombian Chocó.

This clade, the sister taxon to *E.* sp. 1 and *E. narinensis,* includes *E. boulengeri, E.* aff. *espinosai, E. espinosai, E. machalilla, E. tricolor,* and *E. anthonyi* and has an age of 6.02 Mya. Notably, the most recent common ancestor of *E. boulengeri* s. s. and all other species corresponds to the oldest node in this clade; within *E. boulengeri* s. s., the oldest node splits lineages leading to mainland and island populations. *Epipedobates boulengeri* s. s. co-occurs with other Isla Gorgona endemics including another dendrobatid, *Leucostethus siapida* (Grant and Bolívar-Garcia 2021). Despite the proximity of Isla Gorgona to the mainland (25–36 km), the island seems to have remained unaffected by marine incursions, and it is likely a biotic refugium, as the island geology suggests it to be a relatively ancient igneous remnant formed in the Late Cretaceous (Serrano et al. 2011). During low sea levels, the island was reconnected to the continent by a relatively shallow strait that may have impeded movement; the strait is currently under water at 60-120 meters in depth. The connection of the island and mainland may have cycled during glacial periods. The latest continuous land bridge between the island and the continent may have occurred during the last glacial maximum (18–20,000 years ago; Fleming et al. 1998). The most recent event isolating the Isla Gorgona is not clear, but coral reefs around the island might have formed 2,000-3,000 years ago (Glynn et al. 1982; Grant and Bolívar-García 2021). Inland populations of *E. boulengeri* s. s. are located in Maragrícola (Nariño, Colombia) and San Lorenzo/Urbina (Esmeraldas, Ecuador). The great circle distance between the Isla Gorgona population and the closest continental population is ∼150 km, which suggests that other populations might exist in Nariño.

Interestingly, the mainland and island populations of *E. boulengeri* s. s. diverged 3.06 Mya, which suggests that the MRCA of this species’ island and inland populations predates the last time the island was reconnected to the continent. This divergence time is similar to the one calculated for the divergence between *Leucostethus bilsa* (Bilsa, Esmeraldas, Ecuador) and *L. siapida* from Isla Gorgona at 3 Mya (Vigle et al. 2020). These ages suggest that the dendrobatid species from Isla Gorgona and their continental relatives might have a similar pattern of divergence that traced periodic changes in environmental conditions and isolation-reconnection cycles caused by marine incursions during the Miocene/Pliocene geological history of northwestern South America. Our results highlight the relevance of the fauna and flora of Isla Gorgona as an important refugium of biodiversity and a source of new colonizers to the mainland after marine incursions.

The clade including *E.* aff*. espinosai, E. espinosai, E. machalilla, E. tricolor,* and *E. anthonyi* has an age of ∼3.55 My. Species of this clade are mostly endemic to Ecuador and inhabit the Chocó lowlands to the Andean foothills in Colombia (*E.* aff. *espinosai* from La Nutria) and Ecuador (*E. tricolor* found up to heights of 1,800 masl).

Lastly, *E. anthonyi* and *E. tricolor* diverged 1.99 Ma, and *E. machalilla* and *E. tricolor* diverged 1.51 Mya; yet the extent of genetic introgression is to be determined between such closely related species. *Epipedobates machalilla* seems to have a wide distribution, but some regions are not well-sampled. Gene flow between *E. machalilla* and geographically close species such as *E. espinosai* or *E. tricolor* might obscure some of the relationships between each species range and cladogenesis within this clade of *Epipedobates*.

### 4.5. Taxonomic assessments

Based on the phenotypic and phylogenetic results, we propose the recognition of eight species of *Epipedobates*, one of which is a new species under description (*E.* sp. 1), and another of which is a putatively new species under further investigation (*E.* aff. *espinosai*).

#### 4.5.1. Epipedobates narinensis Mueses-Cisneros, Cepeda-Quilindo, and Moreno-Quintero, 2008

We propose that *Epipedobates narinensis* is a valid species based on its phenotypic distinctiveness from other *Epipedobates* spp. and the 2.78-My divergence from its sister species, *E.* sp. 1. The species is recorded from only a single site (Biotopo, ∼530 masl; (Mueses-Cisneros, 2016). Other geographic surveys of its putative range (southwestern Colombia along the border with Ecuador and northward towards Popayán) have been hindered by ongoing civil conflict (pers. obs. Tarvin, Betancourth-Cundar).

#### 4.5.2. Epipedobates *sp. 1*

We propose that *Epipedobates* sp. 1 is a new species based on its monophyly, its phenotypic distinctiveness from other *Epipedobates* spp., and 2.78-My divergence from its sister species, *E. narinensis.* This species is currently being described using additional criteria by M. Betancourth-Cundar and others. Lötters et al. (2003) described a call from Anchicayá, Colombia; these frogs are referable to *E.* sp. 1. The elevational range of the three populations we collected is 0–705 masl.

#### 4.5.3. Epipedobates boulengeri *(Barbour 1909)*

*Epipedobates boulengeri* is a replacement name for *Prostherapis femoralis* Barbour 1905; it has no junior synonyms. We propose that *E. boulengeri* is a valid species based on its phenotypic distinctiveness from other *Epipedobates* spp. and its relatively deep divergence (6.02 My) from other *Epipedobates* spp. (Figs. 1, 2). *Epipedobates boulengeri* s. s. includes individuals from the type locality on Isla Gorgona, Colombia (Barbour, 1909) and from northwestern Ecuador and the Pacific coast of southern Colombia (Fig. 1; 0 to 156 masl). The estimated divergence time between mainland and island populations is 3.06 Ma (Fig. 1, Figs. S1, S2).

*Epipedobates boulengeri* s. s. differs from *E. narinensis* and *E.* sp. 1 in lacking the bright yellow groin markings. It differs from *E. narinensis* by having a dark brown rather than greenish dorsum, and by having a bluish-white rather than yellow venter. The lateral line also tends to be whiter and thicker in *E. boulengeri* s. s. than in *E.* sp. 1.

Lötters et al. (2003, 2007) suggested that the nominal species *E. boulengeri* is a complex of multiple species distributed from Colombia to Ecuador. Our analysis verifies this. For example, Lötters et al. (2003) found that the mating calls of the Anchicayá populations of *E. “boulengeri*” (Valle del Cauca, Colombia) differed from Lita populations (Imbabura, Ecuador). Our phylogenetic analyses show that individuals from Anchicayá likely represent *E*. sp. 1, and those from Lita, Ecuador are *E*. aff. *espinosai*.

Anganoy-Criollo and Cepeda-Quilindo (2017) found that tadpoles of *E. boulengeri* from southern Colombia (Maragrícola and Tumaco, Nariño) were phenotypically distinct from tadpoles of *E.* “*boulengeri*” from northwestern Ecuador. However, their Ecuadorian populations include several populations that we attribute to *E.* aff. *espinosai* (Lita, Carchi, Caimito, Esmeraldas, and additional populations from southwestern Esmeraldas province) and *E. espinosai* (Alluriquín, Santo Domingo de los Tsáchilas). Furthermore, at least one Ecuadorian population in their assessment, Urbina, Esmeraldas, we assign to *E. boulengeri* s. s. Although their data suggest some differentiation among tadpoles of these species, it is difficult to assess these differences given that the Ecuadorian tadpoles likely represent three species.

Based on locality, the individuals referred to *E. boulengeri* in Cisneros-Heredia and Kahn (2016:341, map) include *E*. *boulengeri* s. s., *E*. sp. 1, *E.* aff. *espinosai*, and *E*. *espinosai*.

#### 4.5.4. Epipedobates *aff*. espinosai

This putative undescribed species (node I) has no available name. It includes specimens from Lita (Lötters et al. 2003) and other locations in the northwest of Ecuador from 87–907 masl. Frogs from Pachijal (Fig. 2, Supplementary Fig. S13; northwest of Mindo), initially misidentified as *E. darwinwallacei* based on geographic range (they do not share the orange spotted phenotype), are also part of this species. Although the molecular clade has high support in our Bayesian Phylogeny (Fig. 2), the haplotype maps (Supplementary Fig. S3) indicate multiple genetic groups and support historical introgression with *E. machalilla*. The status of this putative species will require further assessment and more comprehensive genetic and phenotypic analyses.

#### 4.5.5. Epipedobates espinosai *(Funkhouser 1956)*

We propose that *Epipedobates espinosai* is a valid species based on its monophyly, its phenotypic distinctiveness from other *Epipedobates* spp., its estimated 2.53-My divergence from sister species *E.* aff. *espinosai* (Fig. 1), and its distinctive call as compared to *E.* aff. *espinosai* (Tarvin et al. 2017; their Fig. 6 “*E. boulengeri*” is *E.* aff. *espinosai*). We apply the name to the populations of individuals (Fig. 2, node J) in Northwestern Ecuador in the provinces of Pichincha, Santo Domingo, Cotopaxi, and Los Ríos from the foothills of the Andes (Mindo and Pucayacu localities) to the western edge of the Santo Domingo province (La Perla), from 152 masl to 1375 masl (Fig. 1). The type locality is “Hacienda Espinosa, elevation about 1,000 ft., 9 km. west of Santo Domingo de los Colorados, Province of Pichincha, northwestern Ecuador” and georeferenced data from the California Academy of Sciences, which holds the holotype, place this at –0.241, –79.259.

We consider *Epipedobates darwinwallacei* Cisneros-Heredia and Yánez-Muñoz 2011 to be a junior subjective synonym of *Epipedobates espinosai* (Funkhouser, 1956). The type-locality of *Epipedobates darwinwallacei* is “Saragoza-Río Cinto (78° 45′ 15.7″ W, 00° 07′ 44.1″ S, 1390 m), on the Lloa-Mindo old road, provincial de Pichincha, República del Ecuador.” The type locality of *E. espinosai* (Hacienda Espinosa) is 57.5 km from the type-locality of *E. darwinwallacei*.

We evaluate the evidence for the synonymization below, based on variation in coloration, tarsal keel morphology, amount of genetic divergence, call characteristics, and tadpole morphology. Our assessment is that none of these characters provides sufficient evidence for two species.

Cisneros-Heredia and Yánez-Muñoz (2011) diagnosed *Epipedobates darwinwallacei* from other *Epipedobates* by two traits, color pattern and tarsal keel. According to their diagnosis, the most obvious difference is the presence of bright orange to yellow irregular spots on the dorsum and a dark venter with large orange spots. They stated that no other *Epipedobates* species have this pattern.

Few populations have been previously attributed to *E. espinosai*, and most references to this species are based on specimens from close to the type locality. Similarly, the dorsal color pattern of *E. espinosai* in Cisneros-Heredia and Yánez-Muñoz (2010:Fig. 1), which is brick-red without pale spots and lacking a distinct lateral line, matches that of the holotype of *E. espinosai* (CAS:SUA:10577) as described by Funkhouser (1956) and by Funkhouser (1951, as *Phyllobates* sp. A). It also matches our images of *E. espinosai* from the Valle Hermoso and Río Palenque populations (Supplementary Figs. S17, S18). The type locality of *E. espinosai* lies between these two localities, 17.4 km from Valle Hermoso and 40.5 km from Río Palenque.

In the phylogeny (Fig. 2), specimens from Mindo, which are close to the type locality and phenotypically similar to the holotype of *E. darwinwallacei*, are nested within a strongly supported molecular clade of populations that phenotypically match *E. espinosai* (Río Palenque, La Perla, Otongachi, Valle Hermoso; Fig. 2, node J). Additional populations in this clade possess phenotypes similar to the *E. darwinwallacei* holotype, although they are not included in the original description of the species. The Pucayacu population has coloration similar to *E. darwinwallacei* but with bright yellow rather than orange dorsal marks (e.g., see Pontificia Universidad Católica del Ecuador [2021]: QCAZ:A:49789, QCAZ:A:40652; and two observations by bruno1958 on iNaturalist, #151284358 and #151284357). However, in our phylogeny the Pucayacu population is not closely related to our Mindo population. The Otongachi population, also not closely related to our Mindo population, has a dorsal color pattern intermediate between *E. espinosai* and *E. darwinwallacei*, with dark-brown/orange mottling on the dorsum (Supplementary Fig. S16). Therefore, the diagnostic coloration used in the description of *E. darwinwallacei* appears to be part of a continuum of variation that includes intermediate forms (Otongachi) and homoplastic variants (Pucayacu).

In addition to coloration, the shape of the tarsal keel was used by Cisneros-Heredia and Yánez-Muñoz (2010) to diagnose *E. darwinwallacei*. They described the keel as having two conditions: “tarsal keel straight or weakly curved extending proximolateral from preaxial edge of inner metatarsal tubercle or short, tubercle-like, transversely across tarsus, not extending from metatarsal tubercle.” which matches states 0 and 2 of Grant et al. (2006). However, it is not entirely clear whether they intended to reflect true intraspecific variation (polymorphism of states 0 and 2) or uncertainty in the characterization of the keel shape as either state 0 or 2.

In contrast, Cisneros-Heredia and Yánez-Muñoz (2010) characterized the tarsal keel of *E. anthonyi, E. tricolor, E. machalilla, E. boulengeri, E. espinosai*, and *E narinensis* as “large and strongly curved.” This corresponds more or less to state 1 of Grant et al. (2006), which describes the keel as “strong, tuberclelike (=enlarged, curved) proximally, ext[ending] from mt[metatarsal] tub[tubercle]”.

We observed that specimens of *E. espinosai* from the vicinity of the type-locality of this species and specimens of *E. darwinwallacei* from the Mindo area were difficult to characterize as either state 0 or 2, and we were unable to detect any consistent differences in keel shape between the two nominal species.

Other evidence indicates that keel shape is too variable for distinguishing the two species. Grant et al. (2006) scored *E. tricolor*, *E. anthonyi*, *E. machalilla*, *E. boulengeri*, and *E. espinosai* (near Santo Domingo and Río Palenque) as state 2. Grant et al. (2017; Appendix 5) scored three specimens they identified as *E. darwinwallacei* as 0 & 2, identical to the scoring of Cisneros-Heredia and Yánez-Muñoz (2010). These specimens—from Mindo, Union del Toachi, and Río Palenque—all correspond to populations included in clade J and assigned to *E. espinosai* in our study. Thus, specimens of *E. espinosai* have been inconsistently scored as state 2 or state 0 & 2. Given this, we conclude that the condition of the keel is not useful for distinguishing these populations and thus is not useful for diagnosing *E. darwinwallacei* from other species.

The genetic divergence between populations of *E*. *espinosai* and *E*. *darwinwallacei* is extremely low. We compared specimens from our Mindo locality (Fig. 2, Supplementary Fig. S15), which is 6.1 km from the type locality of *E. darwinwallacei*, to our specimens from Valle Hermoso (Supplementary Fig. S18), which is 17.4 km from the type locality of *E. espinosai*. The uncorrected genetic divergence between Mindo and Valle Hermoso is only 0.61–0.91% for *CYTB* and 0.15% for *12S–16S* (a difference of one base for *12S–16S*), which would be extremely low for two distinct species.

In contrast, the mean genetic distance between the *E. espinosai* clade and its sister-species *E.* aff. *espinosai* (node I) is 4.7% (3.6–5.7%, min-max) for *CYTB* and 1.6% (0.89–2.4%) for *12S–16S*. Surprisingly, our Mindo (*E. espinosai*) and Pachijal (*E.* aff. *espinosai*) localities are only 25.4 km apart, indicating clear differentiation over a short distance. Cisneros-Heredia (2016b) showed several map localities for *E. espinosai* as “unverified” and further sampling is needed.

Advertisement calls and tadpole morphology also verify close phenotypic similarity between *E. darwinwallacei* and *E. espinosai* and do not provide evidence of species-level differentiation. Tarvin et al. (2017a: Fig. 6) found very similar call structure between *E. espinosai* and *E. darwinwallacei* in a non-metric multidimensional scaling of acoustic variation. Additionally, tadpole features, such as the posteroventrad orientation of the vent tube and the conical spiracle, are common to *E. espinosai* (from Río Palenque) and *E*. *darwinwallacei* (from near Mindo), but different from other species of *Epipedobates* (dos Santos Dias et al., 2018).

The combined genetic and phenotypic data (dorsal coloration, tarsal keel shape, tadpole morphology, and advertisement call) indicate no substantial differences between the nominal species. Therefore, we apply the name *E. espinosai*, which has nomenclatural priority, to these populations.

Figures 519 and 520 in Lötters et al. (2007) are referable to this species.

#### 4.5.6. Epipedobates machalilla *(Coloma 1995)*

We propose that *Epipedobates machalilla* is a valid species with no synonyms based on its phenotypic distinctiveness from other *Epipedobates* spp. including its distinctive advertisement call (Tarvin et al. 2017a: Fig. 6).

Cisneros-Heredia (2016a) showed map localities for *E. machalilla* as “unverified,” but several are based on the original description by Coloma (1995) and so can be accepted as accurate. The localities on their map occurring east of the Andes are not correct; none of the ranges of currently known *Epipedobates* extend east of the Andes.

*Epipedobates machalilla* from near the type locality possess an inguinal flash mark that varies from light orange to bright red. Other populations have a similar flash mark: Camarones, Manabí (Pontificia Universidad Católica del Ecuador [2021]: QCAZ:A:67469), Reserva Ecológica Loma Alta, Santa Elena (Pontificia Universidad Católica del Ecuador [2021]:QCAZ:A:70322), Manglares-Churute, Guayas (Pontificia Universidad Católica del Ecuador [2021]: QCAZ:A:73078). However, in the other populations we reviewed here this mark is absent. It is plausible that the flash mark arose in the common ancestor of *E. machalilla, E. tricolor,* and *E. anthonyi* (Tarvin et al. 2017a) and was lost in some populations of *E. machalilla*. However, ongoing gene flow between *E. machalilla* and *E. anthonyi* or *E. tricolor* might also introduce genetic variation for flash markings into *E. machalilla*.

#### 4.5.7. Epipedobates tricolor *(Boulenger 1899)*

We propose that *Epipedobates tricolor* is a valid species based on its phenotypic distinctiveness from other *Epipedobates* spp. and its unique advertisement calls (Tarvin et al. 2017a: Fig. 6); there are no junior synonyms. See section 4.5.8 below on *E. anthonyi* for discussion of Duellman and Wild’s (1993) treatment of *Colostethus paradoxus* Rivero 1991. We note that *E. tricolor* is only 1.04% divergent from its sister species *E. machalilla* and thought to have diverged recently in the last 1.51 My.

#### 4.5.8. Epipedobates anthonyi *(Noble 1921)*

We propose that *Epipedobates anthonyi* is a valid species based on its unique advertisement call (Tarvin et al. 2017a: Fig. 6) and its phenotypic distinctiveness from other *Epipedobates* spp. Its type locality is “small stream at Salvias, Prov. del Oro, Ecuador.”

The nominal species *Colostethus paradoxus* was named by Rivero (1991) (type locality: “Lamtac, Cuenca, 2,535 m, Provincia Azuay, Ecuador”) but was synonymized by Rivero and Almendáriz C. (1992 “1991”:106) under *E. tricolor*, citing Luis Coloma as the authority.

Duellman and Wild (1993) also considered *C. paradoxus* Rivero 1991 to be a junior subjective synonym of *E. tricolor,* but they also considered *E. anthonyi* to be a synonym of *E. tricolor* as well. Given that *E. anthonyi* and *E. tricolor* are now considered to be distinct species, which one is the senior subjective synonym of *C. paradoxus*? We could not locate “Lamtac,” the type locality of *C. paradoxus*, in any gazetteers or in the Google Earth application. However, the city of Cuenca is clearly within the distribution of *E. anthonyi*, so it seems more likely that *C. paradoxus* is a junior synonym of *E. anthonyi* and not *E. tricolor*, which is distributed further north.

### 4.6. Other species previously attributed to the Epipedobates clade

#### 4.6.1 Epipedobates maculatus *(Peters 1873)*

The nominal species *Dendrobates trivittatus* var. *maculata* Peters, 1873 was long considered a junior synonym of *D*. *auratus* by Dunn (1931), Savage (1968), and Silverstone (1975). Myers (1982) called attention to its taxonomic uncertainty but retained it in *Dendrobates*. Myers (1987) moved the species into *Epipedobates*, which at that time included species now in *Ameerega*. Grant et al. (2006) provisionally placed the species in *Ameerega*, but Grant et al. (2017) transferred the species to *Epipedobates*; see discussion therein. Grant et al. (2017) noted that *E*. *maculatus* was the only species of *Epipedobates* lacking an oblique lateral stripe. We found the same character state in the Pucayacu population of *E*. *espinosai* (see section 4.5.5). At the moment we have no reason to allocate this species to a different genus based on morphology. However, the purported type locality (“Chiriqui”) is well outside the range of all species of *Epipedobates*, suggesting that it may not be correctly allocated to genus. At the time of the description “Chiriqui” included both Atlantic and Pacific slopes of extreme Panama (Myers, 1982), and no other species of *Epipedobates* are known from northern Colombia or Panama.

#### 4.6.2. Epipedobates robnatali *van der Horst and Woldhuis 2006*

This is a *nomen nudum* that resulted from an invalid nomenclatural act (Art. 16.4 of the International Code of Zoological Nomenclature, 1999) because the description lacks explicit designation of a holotype. The type locality was stated as “near Rio Mindo on trail from Mindo, ca. 1800 m elevation, Pichincha, Ecuador.”, very near the type locality of *E. darwinwallacei*.

## 5. Conclusion

*Epipedobates* forms a radiation of species on the west of the Andes in northern Peru, Ecuador, and Colombia. *Epipedobates* is a model clade for studying the relationship between phenotypic and genetic divergence. The boundaries between species pairs are difficult to assess. We found that the nominal species *Epipedobates “boulengeri”* included cryptic and understudied diversity; i.e., individuals included in that species now correspond to *E. espinosai*, *E.* aff. *espinosai,* and *E.* sp. 1. In contrast, a relatively young clade contains three phenotypically distinctive species, *E. anthonyi, E. machalilla* and *E. tricolor*, with very low genetic divergence despite high levels of morphological divergence. Two of these species might have evolved conspicuous coloration independently, yet introgression and relatively recent divergence in less than 2 Mya might also explain such events. We found evidence of gene flow between *E. machalilla* and *E.* aff. *espinosai*, and we suggest that further sampling using highly variable markers from genome-wide data is needed to assess introgression between these and possibly other pairs of *Epipedobates* species.

## Authorship Contributions

**Karem López-Hervas:** Fieldwork, Formal Analysis, Investigation, Data Curation, Writing – Original Draft Preparation, Writing – Review & Editing **Juan C. Santos:** Conceptualization, Investigation, Resources, Data Curation, Writing – Original Draft Preparation, Writing – Review & Editing, Supervision **Santiago R. Ron:** Conceptualization, Permits, Investigation, Resources, Data Curation, Writing – Review & Editing, Funding Acquisition, Supervision **Mileidy Betancourth-Cundar:** Fieldwork, Permits, Investigation, Resources, Writing – Review & Editing **David C. Cannatella:** Conceptualization, Formal Analysis, Investigation, Resources, Data Curation, Writing – Original Draft Preparation, Writing – Review & Editing, Supervision, Project Administration, Funding Acquisition **Rebecca D. Tarvin:** Conceptualization, Fieldwork, Formal Analysis, Investigation, Resources, Data Curation, Writing – Original Draft Preparation, Writing – Review & Editing, Supervision, Project Administration, Funding Acquisition

## Declaration of Generative AI and AI-assisted technologies in the writing process

A translation of the main text of the manuscript (Supplementary Material text) was first conducted using DeepL and then corrected by Karem López-Hervas, Rebecca Tarvin, and Santiago Ron. No generative or AI-assisted technologies were used for any other purpose during the production of this work.

## Acknowledgments

This work was supported by grants to RDT from the Society of Systematic Biologists, North Carolina Herpetological Society, Society for the Study of Reptiles and Amphibians, Chicago Herpetological Society, Texas Herpetological Society, the EEB Program at University of Texas at Austin, National Science Foundation Graduate Research Fellowship Program Graduate Research Opportunities Worldwide, National Geographic Young Explorer Grant (#9468-14).

RDT and MBC thank Santiago Vega, Daniela Pareja Mejía, Cristina Toapanta, Cristian Flórez Pai, Sandra Victoria Flechas, Daniel Nastacuaz and Nataly Portillo for assistance in fieldwork. Adolfo Amézquita facilitated permitting processes in Colombia. JCS was supported by a NSF-Macrosystems grant and acknowledges Jack Sites, Jr. (BYU) for his support and encouragement. DCC and RDT were supported by NSF 1556967. We also thank ANDES and QCAZ museum personnel including Andrew Crawford for their assistance in accessioning specimens. Field and laboratory work in Ecuador was partly funded by Secretaría Nacional de Educación Superior, Ciencia, Tecnología e Innovación del Ecuador SENESCYT (Arca de Noé initiative; SRR and Omar Torres principal investigators) and a grant from Pontificia Universidad Católica del Ecuador, Dirección General Académica. We thank three reviewers for their comments, which strengthened and clarified the manuscript. The authors declare no competing interests.

This publication is based in part on work by DCC while serving at the National Science Foundation. Any opinion, findings, and conclusions or recommendations expressed in this material are those of the author(s) and do not necessarily reflect the views of the National Science Foundation or United States government.

## Appendix A. Supplementary Material

All supplementary material is also available at https://doi.org/10.5061/dryad.4xgxd25h3 **Supplementary text.** A Spanish translation of the main text of the manuscript.

**Table S1.** Information about specimens and GenBank accession numbers used in this study.

**Table S2.** PCR primers and conditions used to amplify *CYTB, 12S-16S, BMP2, CR,* and *H3.*

**Table S3.** Summary of haplotype data

**Figure S1.** Complete maximum likelihood phylogeny.

**Figure S2.** Bayesian chronogram using a subset of tips that represent the major clades in Figures 1 and 2.

**Figure S3.** Top: Distribution of p-values from GMYC analyses of 1000 trees from the posterior distribution of MrBayes chronogram. The vertical blue line indicates p = 0.05. Bottom: Frequency of the number of inferred species based on GMYC analyses of 1000 trees from the posterior distribution.

**Figure S4.** *Epipedobates narinensis*, Biotopo: ANDES:A:3704–ANDES:A:3711

**Figure S5.** *Epipedobates* sp. 1, Anchicayá: ANDES:A:371–ANDES:A:3720

**Figure S6.** *Epipedobates* sp. 1, Ladrilleros: ANDES:A:2458–ANDES:A:2465

**Figure S7.** *Epipedobates* sp. 1, La Barra and Pianguita: ANDES:A:2455, ANDES:A:3690– ANDES:A:3694

**Figure S8.** *Epipedobates boulengeri* s. s., Isla Gorgona: ANDES:A:3695–ANDES:A:3701

**Figure S9.** *Epipedobates boulengeri* s. s., Maragrícola: ANDES:A:2468–ANDES:A:2475

**Figure S10.** *Epipedobates* aff. *espinosai*, Bilsa: QCAZ:A:53634–QCAZ:A:53642

**Figure S11.** *Epipedobates* aff. *espinosai*, Cube: QCAZ:A:58233–QCAZ:A:58240

**Figure S12.** *Epipedobates* aff. *espinosai*, Lita: QCAZ:A:58221–QCAZ:A:58228

**Figure S13.** *Epipedobates* aff. *espinosai*, La Nutria and Pachijal: ANDES:A:2476, QCAZ:A:58335–QCAZ:A:58336

**Figure S14.** *Epipedobates espinosai*, La Perla: QCAZ:A:53626–QCAZ:A:53633

**Figure S15.** *Epipedobates espinosai*, Mindo: QCAZ:A:58278–QCAZ:A:58285

**Figure S16.** *Epipedobates espinosai*, Otongachi: QCAZ:A:58259–QCAZ:A:58266

**Figure S17.** *Epipedobates espinosai*, Río Palenque: QCAZ:A:58270–QCAZ:A:58277

**Figure S18.** *Epipedobates espinosai*, Valle Hermoso: QCAZ:A:58245–QCAZ:A:58254

**Figure S19.** *Epipedobates machalilla*, 5 de Agosto: QCAZ:A:53643–QCAZ:A:53650, QCAZ:A:58231–QCAZ:A:58232

**Figure S20.** *Epipedobates machalilla*, Jouneche: QCAZ:A:53667–QCAZ:A:53674

**Figure S21.** *Epipedobates machalilla*, Río Ayampe: QCAZ:A:58291–QCAZ:A:58298

**Figure S22.** *Epipedobates tricolor*, Chazo Juan: QCAZ:A:53659–QCAZ:A:53666, QCAZ:A:58307

**Figure S23.** *Epipedobates tricolor*, Guanujo: QCAZ:A:53651–QCAZ:A:53658, QCAZ:A:58308

**Figure S24.** *Epipedobates tricolor*, San José de Tambo: QCAZ:A:58299–QCAZ:A:58306

**Figure S25.** *Epipedobates anthonyi*, Moromoro: QCAZ:A:53683–QCAZ:A:53690, QCAZ:A:58290

**Figure S26.** *Epipedobates anthonyi*, Pasaje: QCAZ:A:53675–QCAZ:A:53682

**Figure S27.** *Epipedobates anthonyi*, Uzchurummi: QCAZ:A:53691–QCAZ:A:53698

**Data S1.** Raw data (alignments) and scripts for constructing trees and haplotype networks, including the resulting tree files in newick format.

Data S1A: Epipedobates-6Genes-120-FINAL.nex – Alignment for IQTREE

Data S1B: run_Epipedobates_6Genes.sh – script for IQTREE

Data S1C: Epipedobates-6Genes-120-FINAL-Partitions-Invar-Omitted.nex – partition file for IQTREE

Data S1D: IQTREE2-MaximumLikelihood-Analysis8-For-FigS1.tre – Output tree file from IQTREE

Data S1E: Epipedobates-6Genes-120-FINAL-Analysis4.nex– Alignment for MrBayes

Data S1F: Bayesblock_Epipedobates.txt – Code to run MrBayes

Data S1G: MrBayes-120-FINAL-Analysis4-MCCT-For-Fig2.tre – Output tree file from MrBayes

Data S1H: Epipedobates-Pruned_IGR-TT10.nex – Code and alignment to run MrBayes pruned timetree (18 tips)

Data S1I: MrBayes-Chronogram-18Tips-TT10-For-FigS2.tre – Output tree file from MrBayes pruned timetree analysis

Data S1J: Epipedobates_12S_haplotypes.fasta – Haplotypes for *12S-16S*

Data S1K: Epipedobates_ControlRegion_haplotypes.fasta – Haplotypes for *CR*

Data S1L: Epipedobates_cytb_haplotypes.fasta – Haplotypes for *CYTB*

Data S1M: Epipedobates_BMP2_haplotypes.fasta – Phased haplotypes for *BMP2*

Data S1N: Epipedobates_H3_haplotypes.fasta – Phased haplotypes for *H3*

Data S1O: Epipedobates_KV13_haplotypes.fasta – Phased haplotypes for *K_V_1.3*

Data S1P: Epipedobates_Haplotype_Networks.Rmd – R script to analyze haplotypes

**Data S2**. Quantitative species delimitation analyses

Data S2A: Epipedobates_mtDNA-fewremoved_only_ASAP.fasta – input alignment for ASAP

Data S2B: Epipedobates_mtDNA-fewremoved_only_ASAP.fasta.spart– ASAP logfile

Data S2C: Epipedobates_mtDNA-fewremoved_only_ASAP.fasta.Partition_1.txt – ASAP partition results for N = 2 species groups

Data S2D: infile.nex – Infile for MrBayes mtDNA-only chronogram tree

Data S2E: GMYC4_Epipedobates-12S16S_CYTB_CR_IGR.log.txt – logfile for MrBayes

Data S2F: run_Epipedobates-CIPRES-IGR-for-GMYC4.nex – MrBayes script

Data S2G: GMYC_Epipedobates.Rmd – R script to analyze GMYC results

Data S2H: my_1000_GMYC4_Trees.nexus – 1000 trees from MrBayes

Data S2I: GMYC4-Epipedobates-12S16S-CYTB-CR-IGR.con.tre – consensus mtDNA tree

Data S2J: Supplementary-Data-Y-Summary-GMYC-Results.txt – summary of R script results

**Data S3**. Locality data for each species used in constructing Figure 1 and specimen data and R script used to compare SVL among species and with elevation.

Data S3A: Epiped_KLH_NewModel_20181026.voucher.txt – Locality data for each species used in constructing Figure 1

Data S3B: specimen-data-for-SVL_2023-02-05.csv – Specimen SVL, sex, and elevation data

Data S3C: specimen_data_2023-06-19.R– R script to analyze SVL, sex, and elevation

