## Supplementary Table S2 for "Deep Divergences Among Inconspicuously Colored Clades of *Epipedobates* Poison Frogs"

**Table S1.** PCR primers and conditions used to amplify *CYTB, 12S-16S, BMP2, CR,* and *H3.*

| **Gene** | **Primers** | **Sequence** | **PCR protocol** |   **References**   |
| --- | --- | --- | --- | --- |
| cytochrome b (*CYTB*) | CytbDen3-L  CytbDen1-H | Forward 5'-AAYATYTCCRYATGATGRAAYTTYGG-3'  Reverse 5'-GCRAANAGRAAGTATCATTCNGGYT -3' | 2 min 94°C  35 cycles: 30 s 94°C, 30 s 46.5°C, 1 min 72°C  7 min 72°C  infinite 4°C | Goebel et al 1999; Santos and Cannatella 2011 |
| control region (*CR*) | CytbA-L  ControlP-H_64 | Forward 5'GAATYGGRGGWCAACCAGTAGAAGACCC-3'  Reverse 5'- GTCCATAGATTCASTTCCGTCAG-3' | 2 min 94°C  40 cycles: 30 s 94°C, 30 s 46.5°C, 1 min 72°C  7 min 72°C, infinite 4°C |   Palumbi et al 1991; Pauly et al 2004; Heinicke et al 2007 |
| 12S-16S | 12Sa  16SH-H | Forward 5'- AACTGGGATTAGATACCCCACTAT -3'  Reverse 5'- TACCTTTTGCATCATGGTCTAGC -3' | 2 min 94°C  35 cycles: 30 s 94°C, 30 s 46.5°C, 1 min 72°C  7 min 72°C, infinite 4°C | Santos and Cannatella 2011 |
| histone H3 (*H3*) | Forward  Reverse | Forward 5'- ATGGCTCGTACCAAGCAGACVGC-3'  Reverse 5'- ATATCCTTRGGCATRATRGTGAC-3' | 3 min 94°C  35 cycles: 30 s 94°C, 30 s 60°C, 1 min 72°C  7 min 72°C, infinite 4°C | Colgan et al 1999 |
| bone morphogenetic protein 2 (*BMP2*) | BMP2_F7  BMP2_R3 | Forward 5'- TATGATGTACCAGTCTGGTGTCCAATAGTCT -3'  Reverse 5'- CRCAYCCCTCCACRACCATGTCTTGATA-3' | 4 min 94°C  40 cycles: 30 s 94°C, 30 s 50°C, 1.5 min 72°C  7 min 72°C, infinite 4°C | Santos and Cannatella, 2011; Folmer et al 1994 |
