## Supplementary figures and images for "Deep Divergences Among Inconspicuously Colored Clades of *Epipedobates* Poison Frogs"

### Supplementary Figure S1

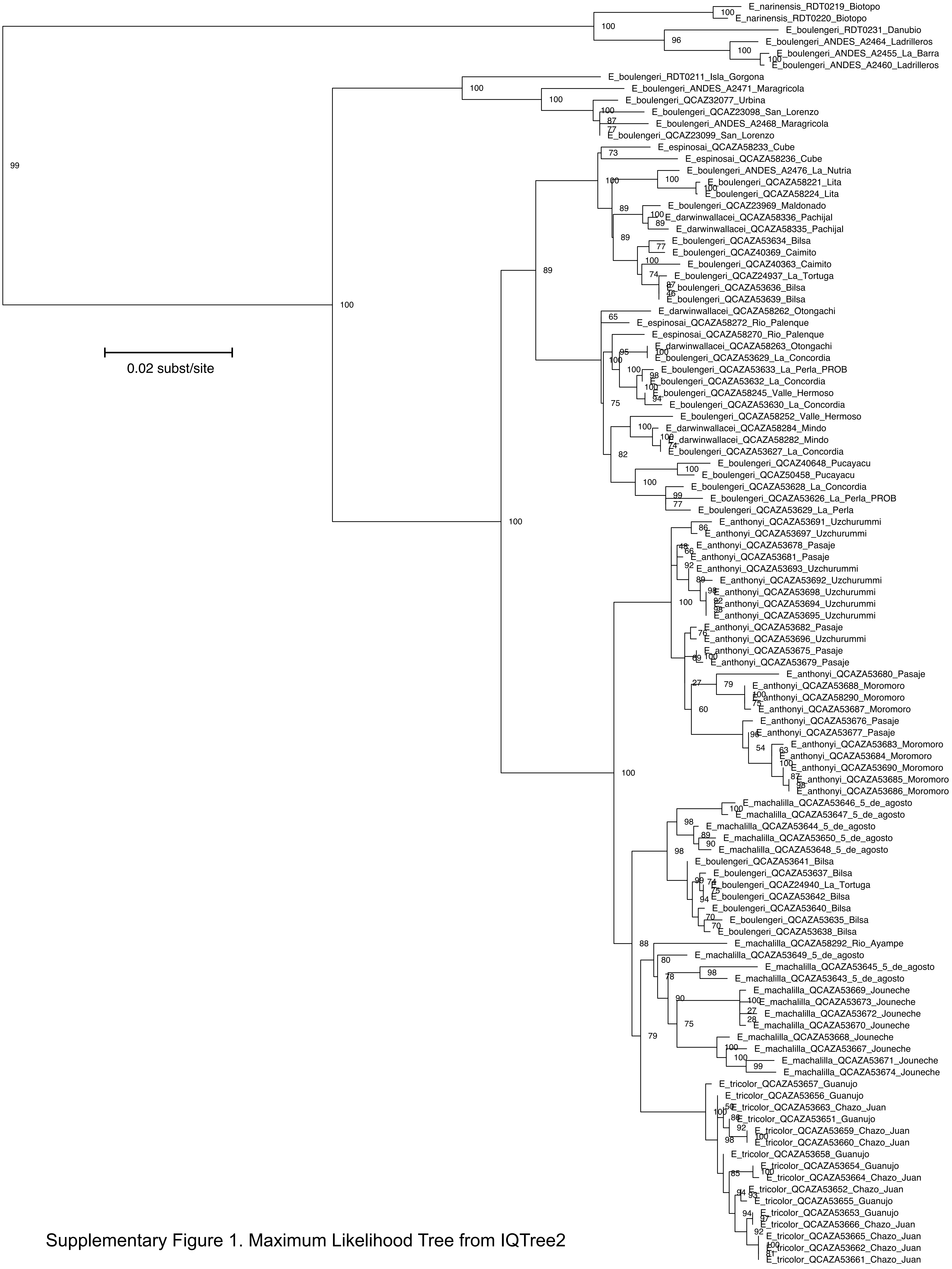


Supplementary Figure 1. Maximum Likelihood Tree from IQTree2

### Supplementary Figure S3

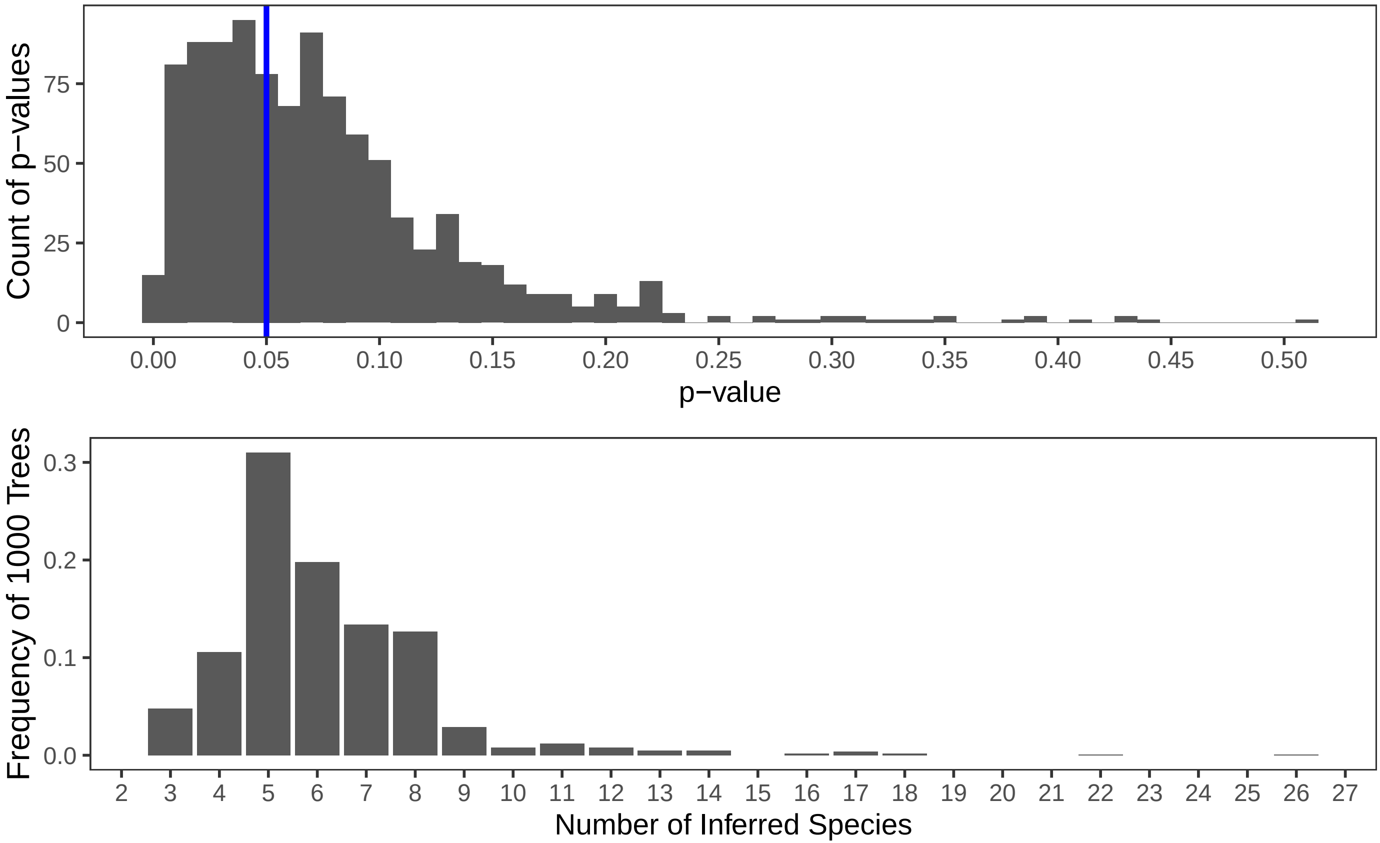
