## Supplementary Figure S2 for "Deep Divergences Among Inconspicuously Colored Clades of *Epipedobates* Poison Frogs"

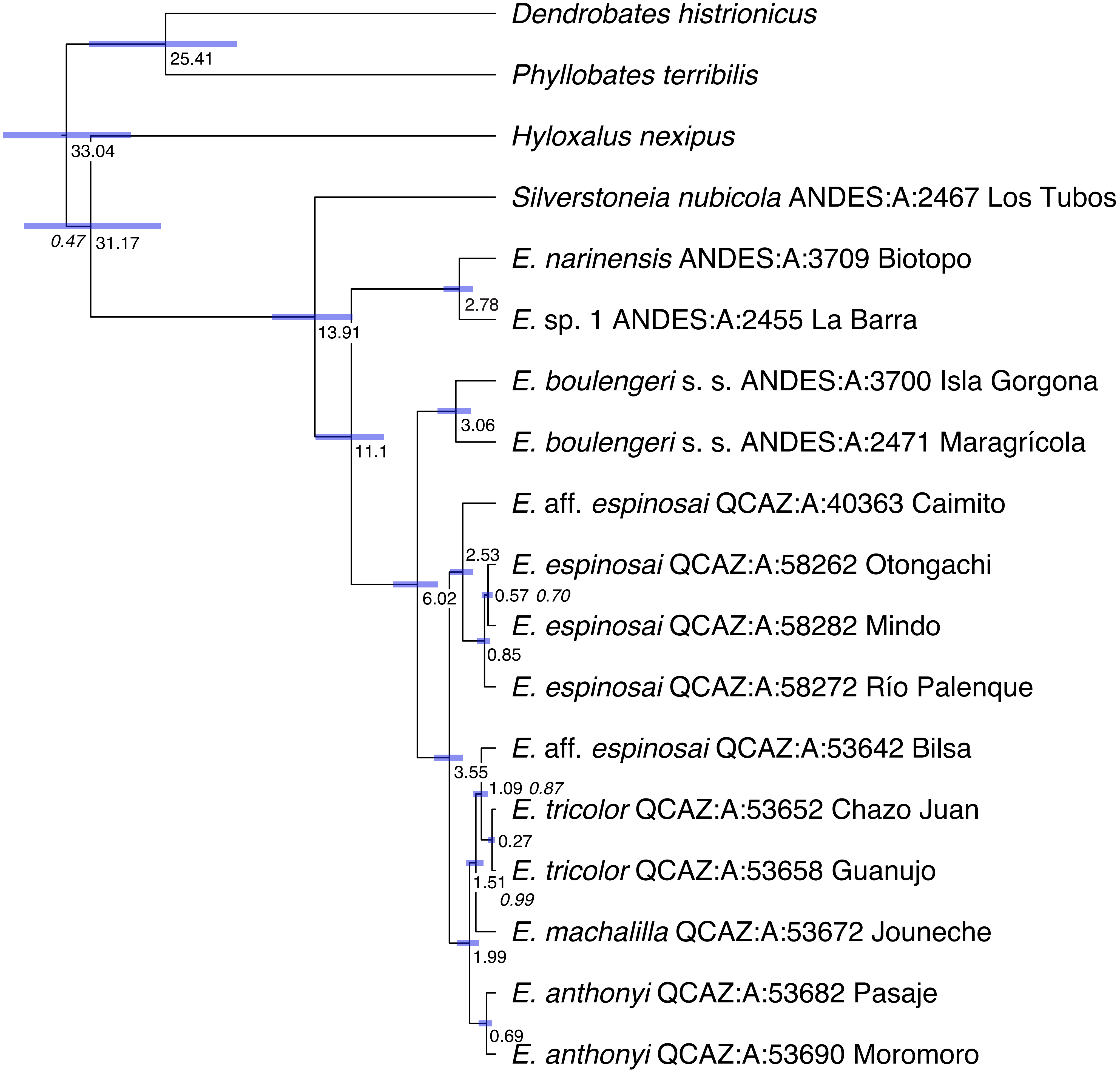


Supplementary Figure S2. Chronogram of *Epipedobates* species. Node ages in my shown in roman typeface. Posterior probability values for all nodes are 1.0, except where indicated with an italic value.
