## Supplementary Data S1 for "Deep Divergences Among Inconspicuously Colored Clades of *Epipedobates* Poison Frogs": run_Epipedobates_6Genes.sh

```
#!/bin/sh

./iqtree2  -s Epipedobates-6Genes-120-FINAL.nex -nt AUTO -spp Epipedobates-6Genes-120-FINAL-Partitions-Invar-Omitted.nex -mfreq F,FO -mrate E,G,I -m TESTMERGE -redo --runs 3 -B 10000 -pre Analysis8
```
