## Supplementary Data S1 for "Deep Divergences Among Inconspicuously Colored Clades of *Epipedobates* Poison Frogs": Epipedobates_Haplotype_Networks.Rmd

````
---
title: "Epipedobates Haplotype Networks"
subtitle: "Epipedobates ms"
author: "Cannatella"

date: "`r format(Sys.time(), '%B %e, %Y')`"
output: 
  bookdown::html_document2:
    toc: yes
    toc_float: yes
    self_contained: TRUE
    number_sections: TRUE
    toc_depth: 4
    pandoc_args: --number-offset=0
    html-math-method: katex
    keep_md: yes
    highlight: rstudio
    theme: cerulean

---

```{r setup, include=FALSE, message = F}
knitr::opts_chunk$set(echo = FALSE)
library(pegas)
library(tidyverse)
library(ape)
library(lubridate)
library(phangorn)
library(purrr)
library(knitr)
library(bookdown)
```

# How to Plot Haplotype Networks

## Introduction

This .Rmd (RMarkdown) script is designed for phylogeographic studies in which the goal
was to analyze each of several genes from a
concatenated gene alignment, such as is commonly used in phylogeographic analyses. 
Typically there will be sequences of several individuals of a several species.

The script was written for our manuscript and is not intended to be
comprehensive in its calcuation of pop gen stats. It can be modified to accommodate
a different number of genes or to calculate additional statistics or
make other types of haplotype networks using the `ape` and `pegas` packages.

## Credit

This script is mostly based on

*Paradis, E. 2018. Analysis of haplotype networks: the randomized minimum spanning tree method. Meth. Ecol. Evol. 9(5), 1308–1317.* https://doi.org/10.1111/2041-210X.12969

*Paradis, E. 2021. Plotting Haplotype Networks with pegas. at https://cran.r-project.org/web/packages/pegas/vignettes/PlotHaploNet.pdf*

## Assumptions and Comments

These scripts assume:

1. The .fasta files of aligned haplotypes are in the same folder as the .Rmd script. The folder is named "data"

2. Nuclear (diploid) sequences have been phased. For this analysis I used *Flot, J-F., 2010. SeqPHASE: a web tool for interconverting PHASE input/output files and FASTA sequence alignments Mol. Ecol. Res. 10, 162–166.* https://doi/10.1111/j.1755-0998.2009.02732.x.

2. The "knitr" button in RStudio can be used to execute this entire script and produce a .html file. I prefer to run each R chunk separately by clicking the green triangle at the upper right of each chunk.
    
     If you are analyzing several genes and like to write R scripts, you may want to modify the code to run all genes at once by incorporating a for loop or an sapply() function.
    
3. To color haplotype symbols by species (or any user-defined group),  
the group labels and their assigned colors are defined in the same order as 
the sequences in the .fasta file.

    For example, the script below specifies that the first 42 sequences 
    are to be grouped as "anthonyi", the next 16 as "espinosai", etc.
    *mySpecies_Names* is the grouping variable. The second line of the script
    assigns the colors of haplotype symbols to *mySpecies_Colors* in the same order.

```
mySpecies_Names <-  c(rep(c("anthonyi"), each = 42),
                      rep(c("espinosai"), each = 16),
                      rep(c("tricolor"), each = 30)
                      )
                      
mySpecies_Colors <-  c("#ad96ba",   # anthonyi
                       "#704985",   # espinosai   
                       "#1b5ea1")   # tricolor      
```

4. This Rmarkdown script embeds the network figure into the html. If you want 
 to adjust the position of haplotype symbols interactively (recommended), and also produce
an editable pdf file, then uncomment the lines indicated in the
`Plot_rmst()` function.

5. For each gene, the script writes a .tsv file with summary stats for the haplotypes

5. The script produces an html file that includes 

* A haplotype network for each gene, following the method used by Paradis (2021. Plotting Haplotype Networks with pegas.) at https://cran.r-project.org/web/packages/pegas/vignettes/PlotHaploNet.pdf

    Specifically, it creates an *rmst*, a randomized minimum spanning tree, as described in *Paradis, E., 2018. Analysis of haplotype networks: the randomized minimum spanning tree method. Meth. Ecol. Evol. 9(5), 1308–1317.* https://doi.org/10.1111/2041-210X.12969

    I prefer the rmst network to others (I don't care for PopArt) because it displays additional edges showing equally parsimonious connections between haplotypes.

* A  table summarizing haplotype statistics of each gene, for all species combined and 
for each species alone. This table is also written to a .tsv file.

-----

## User-Defined Functions

```{r Plot_rmst, echo = T, warning = F, message = F}

# Define function

Plot_rmst <- function(Gene_Name, Fasta_fn, mySpecies_Names, mySpecies_Colors, myScaleRatio) {
    
  # NOTE: The last parameter in this function, myScaleRatio, must be determined by trial and error
  # to get an esthetically pleasing plot.
  
  # Read sequences
  mySequences <- read.FASTA(Fasta_fn)
  
  # Define vector of species names matching order in data matrix.
  # Analysis for KV13./ Only five species have data

  # Convert to sequences to matrix, extract and summarize haplotypes
  mySequences <- as.matrix(mySequences)
  Epi_Haplo <- haplotype(mySequences)
  Summ_Haps <- summary(Epi_Haplo)
  
  ###############
  # Calculate uncorrected p-distance and the
  # randomized minimum spanning tree network.
  # Be sure to use pairwise deletion or else the entire site will be deleted
  # in the calculation of distances
  Epi_Dist <- dist.dna(Epi_Haplo, "N", pairwise.deletion = T)
  # Change the number of randomizations using the B parameter
  Epi_rmst <- rmst(Epi_Dist, B = 10000, quiet = TRUE)
  ###############

  
  # Re-order the labels to match
  (Epi_rmst.labs <- attr(Epi_rmst, "labels"))
  Summ_Haps <- Summ_Haps[Epi_rmst.labs]
  # Get haplotype frequencies to use in piechart, and re-order labels
  Haplo_by_Species <- haploFreq(mySequences, fac = mySpecies_Names, haplo = Epi_Haplo)
  Haplo_by_Species <- Haplo_by_Species[Epi_rmst.labs, ]

  # Make basic haplotype plot
  plot(Epi_rmst, 
       size = Summ_Haps,   # try sqrt(Summ_Haps) if circles are too large
       pie = Haplo_by_Species,
       scale.ratio = myScaleRatio,  # set piechart size relative to segments
       labels = F,
       show.mutation = 2,
       legend = F,
       bg = mySpecies_Colors,
       lwd = 1.2,
       directed = F)
  title(paste0(Gene_Name, " randomized minimum spanning tree, sensu Paradis"))

  # If desired (you will probably say yes), you can interactively drag piecharts 
  # on the graph (cool!) to place them esthetically, and 
  # store the coordinates in a .RDS file for re-plotting as pdf.
  # To do this, you must uncomment the code between the "##############" below
  # Hit escape to exit the interactive mode
  # For more info see the pdf  by Paradis (2021) titled "Plotting Haplotype Networks with pegas"
  ##########################
  # XY_Coords <- replot() 
  # RDS_fn <- paste0(Gene_Name, "_plot_coords.RDS")
  # saveRDS(XY_Coords, file = RDS_fn)
  # XY_Coords <- readRDS(RDS_fn)
  ## Comment: After storing coords, open pdf device and replot using user-defined coords
  # PDF_Name <- paste0("rmst_", Gene_Name, "-", today(), ".pdf")
  # pdf(PDF_Name)
  # replot(XY_Coords)
  # title(paste0(Gene_Name, " randomized minimum spanning tree, sensu Paradis"))
  # dev.off()
  # ##########################
  # Comment: un-comment next line to open pdf in window
  # system(paste0("open ", PDF_Name))
}

```

```{r Haplotype_stats, echo = T, warning = F, message = F}

# Define function
# Do stats on haplotypes only. No plots
Summarize_Haplotypes <- function(Gene_Name, Fasta_fn, mySpecies_Names) {
    
    # Determine Ploidy
    if (Gene_Name %in% c("KV13", "BMP2", "H3")) {
        Ploidy <- 2
    }  else   {
        Ploidy <- 1
    }

    # Read sequences
    mySequences <- read.FASTA(Fasta_fn)
    # Convert to matrix, extract and summarize haplotypes
    mySequences <- as.matrix(mySequences)
    Epi_Haplo <- haplotype(mySequences)
    Summ_Haps <- summary(Epi_Haplo)
    # Get haplotype frequencies
    Haplo_by_Species <- haploFreq(mySequences, fac = mySpecies_Names, haplo = Epi_Haplo)
    # Output some stats for de-bugging
    # print(paste0(Gene_Name, ". Ploidy: ", Ploidy), quote = F)
    # print(paste("Number of sequences is", nrow(mySequences)), quote = F)
    # print(paste("Number of individuals is", nrow(mySequences)/Ploidy), quote = F)
    # print(paste("Number of unique haplotypes is", length(Summ_Haps)), quote = F)
    # print(paste("Number of ALL haplotypes is", sum(Summ_Haps)), quote = F)
    # print(paste("Average number of sequences per unique haplotype is",
    #             round(sum(Summ_Haps)/length(Summ_Haps),3)), quote = F)
    # print(paste("Number of species is", ncol(Haplo_by_Species)), quote = F)
    # 
    
    # Augment matrix to include all 8 species, even those with no haplotypes
    # For some genes, sequences are missing for one or more of the species
    # below. We need to make all matrices the same size, so we add empty
    # column (all 0s)
    if (!"narinensis" %in% colnames(Haplo_by_Species))  {
        Haplo_by_Species <- cbind(Haplo_by_Species, narinensis = 0)
    }
    if (!"sp1" %in% colnames(Haplo_by_Species))  {
        Haplo_by_Species <- cbind(Haplo_by_Species, sp1 = 0)
    }
    if (!"boulengeri" %in% colnames(Haplo_by_Species))  {
        Haplo_by_Species <- cbind(Haplo_by_Species, boulengeri = 0)
    }
    # Sort 8-species matrix by column names
    Haplo_by_Species_Sorted <- Haplo_by_Species[, order(colnames(Haplo_by_Species), decreasing = F)]
    # print(Haplo_by_Species)
    
    # Count number of haplotypes in each species
    Species_Haplotypes <- vector(mode = "character")
    for (i in 1:ncol(Haplo_by_Species)) {
        # print(paste("i =", i))
        Haplotype_Present <- which(Haplo_by_Species_Sorted[,i] != 0)
        Total_Haplotypes <- length(Haplotype_Present)
        # print(paste(as.character(Total_Haplotypes), "haplotypes in", 
        #            colnames(Haplo_by_Species_Sorted)[i]), quote = F)
        
        # Loop over haplotype rows and tabulate unique and non-unique haplotypes
        N_unique <- 0
        N_nonunique <- 0
        # If this species has no haplotypes (meaning no individuals were sequenced
        # for the gene), then do not enter the for loop.
        if (Total_Haplotypes > 0)   {
            for(j in 1:length(Haplotype_Present)) {
                # print(paste("j =", j))
                if(Haplo_by_Species_Sorted[Haplotype_Present[j], i]  ==
                   sum(Haplo_by_Species_Sorted[Haplotype_Present[j], ])) {
                    #print(paste(i, j, "unique"))
                    N_unique <- N_unique + 1
                }  else   {
                    #print(paste(i, j, "not unique"))
                    N_nonunique <- N_nonunique + 1
                }
            }   # end for
        }
        # Next lines for debugging
        # cat(paste("Unique haplotypes = ", N_unique, "\n")) 
        # cat(paste("Non-unique haplotypes = ", N_nonunique, "\n\n")) 
        Species_Haplotypes[i] <- paste0(N_unique, "/", N_nonunique)
    }   # end for
    
    # Complicated way of creating a tibble from all the output
    # Make a first tibble with combined stats for all species combined
    myTibble1 <<- tibble(`Gene` = as.character(Gene_Name), 
                         Ploidy = Ploidy, 
                         `N sequences` = nrow(mySequences),
                         `N individuals` = nrow(mySequences)/Ploidy, 
                         `N haplotypes` = length(Summ_Haps),
                          `Average N sequences/haplotype` =
                             round(sum(Summ_Haps)/length(Summ_Haps),3))
    # Make a second tibble with stats for each species 
    names(Species_Haplotypes) <- colnames(Haplo_by_Species_Sorted)
    Species_Haplotypes <- as.list(Species_Haplotypes)
    myTibble2 <- as_tibble(Species_Haplotypes)
    # Combine the two tibbles and write to file
    myTibble <- bind_cols(myTibble1, myTibble2)
    write_tsv(myTibble, paste0(Gene_Name, "-haplostats.tsv"), quote = "all")
}

```


## Draw the Plots for Each Gene

The six chunks below are used to plot the rmst network and/or calculate the 
haplotype stats for each gene separately. In each script I have commented out the line (2nd to last) that actually runs the analysis so that they don't all run at once.

Example: `Plot_rmst(Gene_Name, Fasta_fn, mySpecies_Names, mySpecies_Colors, 1)`


### Plot KV1.3

(not plotted)
```{r plot_KV13_rmst, fig.height = 10, fig.width = 8, echo = T, warning = F}
# Set the four variables for the function
Gene_Name <- "KV13"
Fasta_fn <- "data/Epipedobates_KV13_haplotypes.fasta"
# The order of items in the vector of species names (group names) must match the 
# order in the fasta alignment
# There are 42 sequences of anthonyi
#           16 sequences of espinosai
# etc
mySpecies_Names <-  c(rep(c("anthonyi"), each = 42),
                  rep(c("espinosai"), each = 16),
                  rep(c("aff_espinosai"), each = 16),
                  rep(c("machalilla"), each = 32),
                  rep(c("tricolor"), each = 30)
                  )
 
# Some species colors are excluded because these species are
# not represented in the matrix
# Clearly, this code could be made more efficient!
mySpecies_Colors <-   c("#ad96ba", # aff espinosai   light purple
                          "#e1c691", # anthonyi        tan
  #                       "#ef6a00", # boulengeri      orange
                          "#704985", # espinosai       dark purple
                          "#9ccbe0", # machalilla      light blue
  #                       "#3f6742", # narinensis      dark green
  #                       "#8ca48e", # sp 1            light green
                          "#1b5ea1") # tricolor        dark blue

# Un-comment next line to plot the network
# Plot_rmst(Gene_Name, Fasta_fn, mySpecies_Names, mySpecies_Colors, 1)

Summarize_Haplotypes(Gene_Name, Fasta_fn, mySpecies_Names)
```

### Plot H3

(not plotted)
```{r plot_H3_rmst, fig.height = 10, fig.width = 8, echo = T, warning = F}
# Set the four variables for the function
Gene_Name <- "H3"
Fasta_fn <- "data/Epipedobates_H3_haplotypes.fasta"

# Analysis for H3./ Only five species have data
# Define vector of species names matching order in data matrix.

mySpecies_Names <-    c(rep(c("sp1"), each = 6),
                rep(c("boulengeri"), each = 10),
                rep(c("aff_espinosai"), each = 26),
                rep(c("espinosai"), each = 26),
                rep(c("machalilla"), each = 6),
                rep(c("tricolor"), each = 2),
                rep(c("anthonyi"), each = 4)
                )

# Some species colors are excluded because these species are
# not represented in the matrix
mySpecies_Colors <-    c("#ad96ba", # aff espinosai   light purple
                        "#e1c691", # anthonyi        tan
                        "#ef6a00", # boulengeri      orange
                        "#704985", # espinosai       dark purple
                        "#9ccbe0", # machalilla      light blue
#                        "#3f6742", # narinensis      dark green
                        "#8ca48e", # sp 1            light green
                        "#1b5ea1") # tricolor        dark blue

# Un-comment next line to plot the network
# Plot_rmst(Gene_Name, Fasta_fn, mySpecies_Names, mySpecies_Colors, 2)

Summarize_Haplotypes(Gene_Name, Fasta_fn, mySpecies_Names)

```

### Plot BMP2

(not plotted)
```{r plot_BMP2_rmst, fig.height = 10, fig.width = 8, echo = T, warning = F}
# Set the four variables for the function
Gene_Name <- "BMP2"
Fasta_fn <- "data/Epipedobates_BMP2_haplotypes.fasta"

# Define vector of species names matching order in data matrix.
mySpecies_Names <-  c(rep(c("narinensis"), each = 4),
                rep(c("sp1"), each = 8),
                rep(c("boulengeri"), each = 12),
                rep(c("aff_espinosai"), each = 26),
                rep(c("espinosai"), each = 26),
                rep(c("machalilla"), each = 6),
                rep(c("tricolor"), each = 2),
                rep(c("anthonyi"), each = 6)
                )

mySpecies_Colors <-    c("#ad96ba", # aff espinosai   light purple
                        "#e1c691", # anthonyi        tan
                        "#ef6a00", # boulengeri      orange
                        "#704985", # espinosai       dark purple
                        "#9ccbe0", # machalilla      light blue
                        "#3f6742", # narinensis      dark green
                        "#8ca48e", # sp 1            light green
                        "#1b5ea1") # tricolor        dark blue

# Un-comment next line to plot the network
Plot_rmst(Gene_Name, Fasta_fn, mySpecies_Names, mySpecies_Colors, 0.2)

Summarize_Haplotypes(Gene_Name, Fasta_fn, mySpecies_Names)
```

### Plot 12S

(not plotted)
```{r plot_12S_rmst, fig.height = 10, fig.width = 8, echo = T, warning = F}
# Set the four variables for the function
Gene_Name <- "12S"
Fasta_fn <- "data/Epipedobates_12S_haplotypes.fasta"

# Analysis for 12S
# Define vector of species names matching order in data matrix.
mySpecies_Names <-    c(rep(c("narinensis"), each = 2),
                rep(c("sp1"), each = 4),
                rep(c("boulengeri"), each = 6),
                rep(c("aff_espinosai"), each = 13),
                rep(c("espinosai"), each = 18),
                rep(c("aff_espinosai"), each = 8),
                rep(c("machalilla"), each = 16),
                rep(c("anthonyi"), each = 22),
                rep(c("tricolor"), each = 15)
                )

mySpecies_Colors <-    c("#ad96ba", # aff espinosai   light purple
                        "#e1c691", # anthonyi        tan
                        "#ef6a00", # boulengeri      orange
                        "#704985", # espinosai       dark purple
                        "#9ccbe0", # machalilla      light blue
                        "#3f6742", # narinensis      dark green
                        "#8ca48e", # sp 1            light green
                        "#1b5ea1") # tricolor        dark blue

# Un-comment next line to plot the network
# Plot_rmst(Gene_Name, Fasta_fn, mySpecies_Names, mySpecies_Colors, 0.05)

Summarize_Haplotypes(Gene_Name, Fasta_fn, mySpecies_Names)
```

### Plot Cytb

(not plotted)
```{r plot_cytb_rmst, fig.height = 10, fig.width = 8, echo = T, warning = F}
# Set the four variables for the function
Gene_Name <- "cytb"
Fasta_fn <- "data/Epipedobates_cytb_haplotypes.fasta"

# Define vector of species names matching order in data matrix.
mySpecies_Names <-    c(rep(c("narinensis"), each = 2),
                rep(c("sp1"),                each = 4),
                rep(c("boulengeri"),         each = 6),
                rep(c("aff_espinosai"),      each = 19),
                rep(c("espinosai"),          each = 18),
                rep(c("machalilla"),         each = 17),
                rep(c("anthonyi"),           each = 24),
                rep(c("tricolor"),           each = 16)
                )

# The colors match the filled circles on the other figures.
mySpecies_Colors <-    c("#ad96ba", # aff espinosai   light purple
                         "#e1c691", # anthonyi        tan
                         "#ef6a00", # boulengeri      orange
                         "#704985", # espinosai       dark purple
                         "#9ccbe0", # machalilla      light blue
                         "#3f6742", # narinensis      dark green
                         "#8ca48e", # sp 1            light green
                         "#1b5ea1") # tricolor        dark blue

# Un-comment next line to plot the network
#Plot_rmst(Gene_Name, Fasta_fn, mySpecies_Names, mySpecies_Colors, 0.05)

Summarize_Haplotypes(Gene_Name, Fasta_fn, mySpecies_Names)

```

### Plot Control Region

(not plotted)
```{r plot_CR_rmst, fig.height = 10, fig.width = 8, echo = T, warning = F}
# Set the four variables for the function
Gene_Name <- "CR"
Fasta_fn <- "data/Epipedobates_ControlRegion_haplotypes.fasta"

# Define vector of species names matching order in data matrix.
mySpecies_Names <-    c(rep(c("narinensis"), each = 2),
                rep(c("sp1"), each = 4),
                rep(c("boulengeri"), each = 5),
                rep(c("aff_espinosai"), each = 13),
                rep(c("espinosai"), each = 13),
                rep(c("machalilla"), each = 3),
                rep(c("anthonyi"), each = 3),
                rep(c("tricolor"), each = 1)
                )

mySpecies_Colors <-    c("#ad96ba", # aff espinosai   light purple
                        "#e1c691", # anthonyi        tan
                        "#ef6a00", # boulengeri      orange
                        "#704985", # espinosai       dark purple
                        "#9ccbe0", # machalilla      light blue
                        "#3f6742", # narinensis      dark green
                        "#8ca48e", # sp 1            light green
                        "#1b5ea1") # tricolor        dark blue

# Un-comment next line to plot the network
#Plot_rmst(Gene_Name, Fasta_fn, mySpecies_Names, mySpecies_Colors, 0.05)

Summarize_Haplotypes(Gene_Name, Fasta_fn, mySpecies_Names)
```

## Print Table of Haplotype Stats


```{r AssembleHaplotypeTable, echo=T, message=FALSE, warning=FALSE}

# Simple script to assemble table of Haplotype stats in tab-delimited format
# based on the stats for each gene

# Get the names of the .tsv files, one for each gene
my_haplotype_files <- dir(pattern = "*haplostats.tsv")

my_Table <- my_haplotype_files %>%
    map(read_tsv) %>%    # read in all the files individually
    reduce(rbind)        # reduce with rbind into one dataframe

# Sort rows and write to final table as .tsv
my_Table <- arrange(my_Table, Ploidy, `N haplotypes`)
write_tsv(my_Table, "Final_Haplotype_Table.tsv")

# Now, output the table into Rmarkdown html document
my_Caption <- paste("Table of Haplotype Statistics for Genes and Species. For each species column (last 7 cols), a slash separates the number of unique haplotypes from the number of non-unique haplotypes.")

knitr::kable(my_Table,
      align = "cccccccccccccc",
      caption = my_Caption)
```

end

---
````
