## Supplementary Data S2 for "Deep Divergences Among Inconspicuously Colored Clades of *Epipedobates* Poison Frogs": GMYC_Epipedobates.Rmd

````
---
title: "GMYC for Epipedobates"
output: html_document
author: David Cannatella
---


```{r setup, include=FALSE}
knitr::opts_chunk$set(echo = TRUE)

library(tidyverse)
library(ggplot2)
library (ape)
library(phytools)
library(splits)
library(cowplot)
library(RColorBrewer)
sessionInfo()
```

This is an Rmarkdown script, and requires that package.

```{r GMYC_Epipedobates, fig.height = 11, fig.width = 8.5}
# Assign a short name and read the chronogram
Short_name <- "GMYC4"
# This tree has only tips with unique sequences
Tree_fn <- "GMYC4-Epipedobates-12S16S-CYTB-CR-IGR.con.tre"
my_Tree <- ladderize(read.nexus(Tree_fn))
plot(my_Tree)

# Get the branching times for all internal nodes and sort them
my_Sorted_Times <- sort(branching.times(my_Tree), decreasing = T)
my_Sorted_Times_Node_Nums <- as.numeric(names(my_Sorted_Times))

# Associate new GMYC node numbers with original tree node numbers and sort
# Make two data frames with nodes sorted either by GMYC or Original Numbers
# Node labels sorted by GMYC labels
my_Nodes_Sorted_by_GMYC <- bind_cols(GMYC_Node_Nums = 1:my_Tree$Nnode, Orig_Node_Nums = my_Sorted_Times_Node_Nums)
# Node labels sorted by original node labels
my_Nodes_Sorted_by_Orig <- arrange(my_Nodes_Sorted_by_GMYC, Orig_Node_Nums, GMYC_Node_Nums)

# RUN GMYC analysis!
GMYC_Results <- gmyc(my_Tree, method = "single", interval = c(0, 5))

# Make tibble of results for further use
my_Results_tbl <- bind_cols(Likelihood     = GMYC_Results[["likelihood"]],
                            my_Threshold   = GMYC_Results[["threshold.time"]],                     
                            GMYC_Node_Nums = my_Nodes_Sorted_by_GMYC$GMYC_Node_Nums,
                            Orig_Node_Nums = my_Nodes_Sorted_by_GMYC$Orig_Node_Nums,
                            Num_Clusters   = GMYC_Results[["cluster"]],
                            Num_Entities   = GMYC_Results[["entity"]])
print(my_Results_tbl)

# Extract number of clusters and Confidence Intervals
# Get the number of inferred clusters from the max index of the cluster vector.
# Then get the confidence limits of the inferred number.
# The likelihood ± 2 units provides roughly 95% confidence limits for the best.
# NOTE: These confidence limits are not the same as the *confidence set* below. The latter is the 
# cumulative set of Akaike weights summing to 0.95.
Number_Inferred_Clusters   <- my_Results_tbl$Num_Clusters[which.max(my_Results_tbl$Likelihood)]
CI_Clus    <- vector()
CI_Clus[1] <- min(my_Results_tbl$Num_Clusters[my_Results_tbl$Likelihood > (max(my_Results_tbl$Likelihood) - 2)])
CI_Clus[2] <- max(my_Results_tbl$Num_Clusters[my_Results_tbl$Likelihood > (max(my_Results_tbl$Likelihood) - 2)])
print(paste("Estimated Number of Clusters:", Number_Inferred_Clusters))
print(paste("Cluster CI. Min:", CI_Clus[1], "    Max:", CI_Clus[2]))

# Extract number of species (called "entities") (=clusters + singletons)
# The above rationale also applies to the Entities in object$entity
Number_Inferred_Entities   <- my_Results_tbl$Num_Entities[which.max(my_Results_tbl$Likelihood)]
CI_Ent    <- vector()
CI_Ent[1] <- min(my_Results_tbl$Num_Entities[my_Results_tbl$Likelihood > (max(my_Results_tbl$Likelihood) - 2)])
CI_Ent[2] <- max(my_Results_tbl$Num_Entities[my_Results_tbl$Likelihood > (max(my_Results_tbl$Likelihood) - 2)])
print(paste("Estimated Number of Entities (Species):", Number_Inferred_Entities))
print(paste("Entity CI. Min:", CI_Ent[1], "    Max:", CI_Ent[2]))

# Calculate LR test for fun
LR <- 2 * (max(GMYC_Results$likelihood) - GMYC_Results$likelihood[1])
pvalue <- 1 - pchisq(LR, df = 2) # 0.05669745
# Check critical values of chisq
curve(dchisq(x, 1),  from = 0 , to = 10, n = 1000)
qchisq(0.95, 1)  # 3.841459
qchisq(0.95, 2)  # 5.991465


# Confidence Set of nodes: those within ± 2 likelihood units of best node
# Get the numbers of labels of GMYC nodes that are within CI
ConfLimits_GMYC_Numbers <- which(my_Results_tbl$Likelihood > (max(my_Results_tbl$Likelihood) - 2))
# Get Original Node numbers, threshold times, and likelihoods.
ConfLimits_Orig_Numbers <- my_Results_tbl$Orig_Node_Nums[which(my_Results_tbl$Likelihood > (max(my_Results_tbl$Likelihood) - 2))]
ConfLimits_GMYC_Thresholds <- my_Results_tbl$my_Threshold[ConfLimits_GMYC_Numbers]
ConfLimits_GMYC_Likelihoods <- my_Results_tbl$Likelihood[ConfLimits_GMYC_Numbers]

# Extract 95% confidence set of nodes of Models (all the elements of my_Confset_Obj[["aw"]])
my_Confset_Obj <- confset.gmyc(GMYC_Results, p = 0.95)
my_Confset_GMYC_Numbers <- names(my_Confset_Obj[["aw"]])
my_Best_Model_GMYC_Num <- my_Confset_GMYC_Numbers[which.max(my_Confset_Obj[["aw"]])]
my_Best_Model_Orig_Num <- my_Results_tbl$Orig_Node_Nums[as.numeric(my_Best_Model_GMYC_Num)]
my_Best_AIC_Weight <- my_Confset_Obj[["aw"]][my_Best_Model_GMYC_Num]
my_Model_Confset_Orig_Nums <- as.numeric(my_Sorted_Times_Node_Nums[as.numeric(my_Confset_GMYC_Numbers)])

# Get AIC-based support values for plotting on tree later
# The support value of a node is defined as the
# sum of Akaike weights of candidate delimitation
# models in which the node is included.
my_Support_Values <- gmyc.support(GMYC_Results, p = 0.95) 
my_Support_Values[my_Support_Values == 0] <- NA

# Save lots of basic results to text file.
# Write various results to text file
my_Divider <- paste(replicate(60, "="), collapse = "") 
sink(paste0(Short_name, "-Summary_GMYC_results.txt"))
cat(paste("Results of GMYC analysis of ", dQuote(Tree_fn), "\n", today()))
cat("\n", my_Divider, "\n\n")
cat("----------Summary --------\n")
summary(GMYC_Results)   # Show summary with confidence intervals!
GMYC_Results
cat("\n", my_Divider, "\n\n")
cat("----------Species IDs --------\n\n")
Species_IDs <- spec.list(GMYC_Results) # Show species-sample associations
colnames(Species_IDs)[1] <- "Species_assignment"
print.data.frame(Species_IDs, row.names = F)
cat(my_Divider, "\n")
cat("\n", "---Confidence Set of Models; Nodes with AIC weights--------\n")
my_Confset_Obj[["aw"]]
cat(paste("\nBest AIC Weight:", 
          round(my_Best_AIC_Weight, 5), 
          "at GMYC node", 
          my_Results_tbl$GMYC_Node_Nums[as.numeric(my_Best_Model_GMYC_Num)]))
cat(paste("\n\nCumulative AIC weight of", length(my_Confset_Obj[["aw"]]), "models:", round(sum(my_Confset_Obj[["aw"]]), 4)))
sink()


############
# Plot tree with best value and confidence intervals for Entities
# Also plot both new (GMYC) and original node numbers for reference
my_Best_lik <- round(max(my_Results_tbl$Likelihood), 4)
my_Best_Node <- my_Results_tbl$GMYC_Node_Nums[which.max(my_Results_tbl$Likelihood)]
# sort df by original node numbers
GMYC_Nodes_Sorted <- arrange(my_Results_tbl, Orig_Node_Nums)

# Next lines paint each "species" in a different color
# The number of species was determined a posteriori, for plotting
my_PaintedTree <- paintSubTree(my_Tree, node = 83, state = 1, anc.state = "0")
my_PaintedTree <- paintSubTree(my_PaintedTree, 89, state = 2, anc.state = "0")
my_PaintedTree <- paintSubTree(my_PaintedTree, 101, state = 3, anc.state = "0")
my_PaintedTree <- paintSubTree(my_PaintedTree, 115, state = 4, anc.state = "0")
my_PaintedTree <- paintSubTree(my_PaintedTree, 153, state = 5, anc.state = "0")
my_PaintedTree <- paintBranches(my_PaintedTree, 8, state = 6, anc.state = "0")

# Set colors for simmap subtree painting
simmap_cols <- brewer.pal(n = 7, name = "Dark2")
names(simmap_cols) <- 0:6
simmap_cols[1] <- "black"

pdf(paste0(Short_name,"-Species-CI.pdf"), height = 11, width = 8.5)
# plot(my_Tree, cex = 0.6, plot = F)   # plot = FALSE
# lastPP <- get("last_plot.phylo", envir = .PlotPhyloEnv)
#par(new = T)
#plot(my_Tree, cex = 0.6, plot = T)
plotSimmap(my_PaintedTree, colors = simmap_cols, 
           node.numbers = F, fsize = 0.6,
           mar = c(5.1, 4.1, 4.1, 2.1), plot = F)
lastPP <- get("last_plot.phylo", envir = .PlotPhyloEnv)
par(new = T)
plotSimmap(my_PaintedTree, colors = simmap_cols, 
           node.numbers = F, fsize = 0.6,
           mar = c(5.1, 4.1, 4.1, 2.1), plot = T)
abline(v = lastPP$xx[c(my_Results_tbl$Orig_Node_Nums[c(min(ConfLimits_GMYC_Numbers), 
                                                       Number_Inferred_Entities, 
                                                       max(ConfLimits_GMYC_Numbers))])], 
       col = "gray")
# Plot Original node labels
#nodelabels(cex = 0.4, frame = "n", adj = c(1.1, -1.2), col = "red")
# Plot GMYC node labels
nodelabels(text = as.character(GMYC_Nodes_Sorted$GMYC_Node_Nums),
           cex = 0.4, frame = "n", adj = c(1.5, 1.20))
nodelabels(node = my_Best_Model_Orig_Num, 
           cex = 1.0, 
           pch = 21,
           bg = "green",
           col = "black")
text(x = lastPP$xx[my_Results_tbl$Orig_Node_Nums][c(min(ConfLimits_GMYC_Numbers), 
                                                         Number_Inferred_Entities,
                                                         max(ConfLimits_GMYC_Numbers))], 
     y = length(my_Tree$tip.label) + 1, 
     #y = 81, 
     labels = as.character(my_Results_tbl$Num_Entities[c(min(ConfLimits_GMYC_Numbers), 
                                                         Number_Inferred_Entities,
                                                         max(ConfLimits_GMYC_Numbers))]))
title(main = paste("Number of Inferred Species (entities) with 95% min and max.\nBest likelihood = ", 
                   my_Best_lik, "at node", my_Best_Node),
      sub = Tree_fn)
dev.off()


# Make three plots in one pdf
# Plot 1 is LTT
# Plot 2 is likelihood through time
# Plot 3 is tree with species clusters in red
pdf(paste0(Short_name,"-Plots.pdf"), height = 11, width = 8.5)
plot(GMYC_Results)
add.scale.bar()
nodelabels(my_Nodes_Sorted_by_Orig$GMYC_Node_Nums, cex = 0.4, 
            frame = "n", adj = c(1.7, 1.5))
# Plot dots on the 95% confidence set of models (nodes)
# This nodelabels() added by DCC
nodelabels(node = my_Model_Confset_Orig_Nums, 
           cex = 0.8, 
           pch = 21,
           bg = "yellow",
           col = "black")
# Plot green dot on the node for the best model
# This nodelabels() added by DCC
nodelabels(node = my_Best_Model_Orig_Num, 
           cex = 1.0, 
           pch = 21,
           bg = "green",
           col = "black")
# Add support values if desired
# nodelabels(round(my_Support_Values, 2), cex = 0.5, frame = "n", adj = c(1.7, 1.5))
title(main = paste("95% Conf set (yellow) of", 
                   length(my_Confset_Obj[["aw"]]), 
                   "Models (not species), and Species Clusters"),
      sub = Tree_fn)
dev.off()


# test for paintSubtree
# Paint clusters in different colors


# Next lines paint each "species" in a different color
my_PaintedTree <- paintSubTree(my_Tree, node = 83, state = 1, anc.state = "0")
my_PaintedTree <- paintSubTree(my_PaintedTree, 89, state = 2, anc.state = "0")
my_PaintedTree <- paintSubTree(my_PaintedTree, 101, state = 3, anc.state = "0")
my_PaintedTree <- paintSubTree(my_PaintedTree, 115, state = 4, anc.state = "0")
my_PaintedTree <- paintSubTree(my_PaintedTree, 153, state = 5, anc.state = "0")

#cols <- paletteer_d("ggsci::default_aaas", 6)
cols <- brewer.pal(n = 6, name = "Dark2")
names(cols) <- 0:5
cols[1] <- "black"
plot(my_Tree, plot = F)
lastPP <- get("last_plot.phylo", envir = .PlotPhyloEnv)
plotSimmap(my_PaintedTree, cols, node.numbers = F, fsize = 0.6)
abline(v = lastPP$xx[c(my_Results_tbl$Orig_Node_Nums[Number_Inferred_Entities])], 
       col = "blue")
```

```{r GMYC_Posterior_Sample_Trees}

# This nexus file contains the last 1000 ultrametric trees from the 
# posterior distribution of the MrBayes analysis
Tree_fn <-  "trees/my_1000_GMYC4_Trees.nexus"
my_Trees <- read.nexus(Tree_fn)
my_p_values <- vector()
my_Num_Entities <- vector()

# Run a GMYC analysis on each of 1000 trees and summarize the
# number of inferred species and distribution of p-values
for (i in 1:length(my_Trees)) {
   my_Sorted_Times <- sort(branching.times(my_Trees[[i]]), decreasing = T)
   my_Sorted_Times_Node_Nums <- as.numeric(names(my_Sorted_Times))
   
   # Associate new GMYC node numbers with original tree node numbers and sort
   # Make two data frames with nodes sorted either by GMYC or Orig Numbers
   # Node labels sorted by GMYC labels
   my_Nodes_Sorted_by_GMYC <- bind_cols(GMYC_Node_Nums = 1:my_Trees[[i]]$Nnode, Orig_Node_Nums = my_Sorted_Times_Node_Nums)
   # Node labels sorted by original node labels
   my_Nodes_Sorted_by_Orig <- arrange(my_Nodes_Sorted_by_GMYC, Orig_Node_Nums, GMYC_Node_Nums)
   
   # RUN GMYC analysis!
   GMYC_Results <- gmyc(my_Trees[[i]], method = "single", interval = c(0, 5), quiet = T)
   # Make tibble of results for further use
   my_Results_tbl <- bind_cols(Likelihood     = GMYC_Results[["likelihood"]],
                               my_Threshold   = GMYC_Results[["threshold.time"]],                     
                               GMYC_Node_Nums = my_Nodes_Sorted_by_GMYC$GMYC_Node_Nums,
                               Orig_Node_Nums = my_Nodes_Sorted_by_GMYC$Orig_Node_Nums,
                               Num_Clusters   = GMYC_Results[["cluster"]],
                               Num_Entities   = GMYC_Results[["entity"]])
   # summary(GMYC_Results)
   LR <- 2 * (max(GMYC_Results$likelihood) - GMYC_Results$likelihood[1])
   pvalue <- 1 - pchisq(LR, df = 2)
   cat("\nlikelihood ratio: \t", LR, "\n", "p = \t\t\t", pvalue, "\n\n", sep = "")
   
   # Extract number of species (called "entities") (=clusters + singletons)
   # The above rationale also applies to the Entities in object$entity
   Number_Inferred_Entities   <- my_Results_tbl$Num_Entities[which.max(my_Results_tbl$Likelihood)]
   my_Best_lik <- max(my_Results_tbl$Likelihood)
   cat(my_Best_lik)
   
   CI_Ent    <- vector()
   CI_Ent[1] <- min(my_Results_tbl$Num_Entities[my_Results_tbl$Likelihood > (max(my_Results_tbl$Likelihood) - 2)])
   CI_Ent[2] <- max(my_Results_tbl$Num_Entities[my_Results_tbl$Likelihood > (max(my_Results_tbl$Likelihood) - 2)])
   my_Best_lik <- round(max(my_Results_tbl$Likelihood), 4)
   print(paste("Estimated Number of Entities (Species):", Number_Inferred_Entities))
   print(paste("Entity CI. Min:", CI_Ent[1], "    Max:", CI_Ent[2]))
   
   my_p_values[i] <- pvalue
   my_Num_Entities[i] <- Number_Inferred_Entities
}

mean(my_Num_Entities)  # = 6.148
table(my_Num_Entities)
sum(my_p_values <= 0.05)  # 402 ≤ 0.05

# Plot the histogram of p-values for 1000 trees
my_Histogram <- ggplot(data.frame(P_Values = my_p_values), aes(x = P_Values)) + 
   geom_histogram(binwidth = 0.01) + 
   xlab("p-value") +
   ylab("Count of p-values") +
   geom_vline(xintercept = 0.05, 
                color = "blue", size = 1.0) +
   scale_x_continuous(n.breaks = 10) +
   theme_bw() +
   theme(axis.title.y = element_text(size = 12)) +
   theme(panel.grid.major = element_blank(), panel.grid.minor = element_blank()) 
   ggsave("GMYC4_Histogram_P_Values.pdf", width = 8.5, height = 5)
   

# Plot the barplot of number of  species inferred for each of 1000
# trees
my_Barplot <- ggplot(data.frame(NumberSpecies = my_Num_Entities), aes(x = NumberSpecies)) + 
   geom_bar(aes(y = (..count..)/sum(..count..))) + 
   scale_x_continuous(breaks = 1:30) +
   xlab("Number of Inferred Species") +
   ylab("Frequency of 1000 Trees") +
   theme_bw() +
   theme(axis.title.y = element_text(size = 12)) +
   theme(panel.grid.major = element_blank(), panel.grid.minor = element_blank()) 
   ggsave("GMYC4_Barplot_Num_Species.pdf", width = 8.5, height = 5)
   
# combine the two plots into one figure
plot_grid(my_Histogram, my_Barplot, ncol = 1)
ggsave("Combined_Post_Analysis_Suppl_Fig_GMYC4.pdf")   

```

```

## This code was used temporarily when dropping redudant tips
## from an existing tree. Not needed now.
#my_Tree <- read.nexus("trees/GMYC4_Epipedobates-12S16S_CYTB_CR_IGR.con.tre")
# Drop outgroup tips and duplicate tips
# DropThese <- grep("E_", my_Tree$tip.label, invert = T)
# DropThese_Dups <- c("E_boulengeri_QCAZA53641_Bilsa",
# "E_boulengeri_QCAZA53642_Bilsa",
# "E_boulengeri_QCAZA53629_La_Concordia",
# "E_boulengeri_QCAZA53627_La_Concordia",
# "E_boulengeri_QCAZA53632_La_Concordia",
# "E_boulengeri_QCAZA53640_Bilsa",
# "E_boulengeri_QCAZA53636_Bilsa",
# "E_machalilla_QCAZA53643_5_de_agosto",
# "E_machalilla_QCAZA53669_Jouneche",
# "E_machalilla_QCAZA53670_Jouneche",
# "E_machalilla_QCAZA53647_5_de_agosto",
# "E_machalilla_QCAZA53671_Jouneche",
# "E_anthonyi_QCAZA53688_Moromoro",
# "E_anthonyi_QCAZA53682_Pasaje",
# "E_anthonyi_QCAZA53697_Uzchurummi",
# "E_anthonyi_QCAZA53686_Moromoro",
# "E_anthonyi_QCAZA53690_Moromoro",
# "E_anthonyi_QCAZA53692_Uzchurummi",
# "E_anthonyi_QCAZA53694_Uzchurummi",
# "E_anthonyi_QCAZA53695_Uzchurummi",
# "E_tricolor_QCAZA53655_Guanujo",
# "E_tricolor_QCAZA53664_Chazo_Juan",
# "E_tricolor_QCAZA53656_Guanujo",
# "E_tricolor_QCAZA53663_Chazo_Juan",
# "E_tricolor_QCAZA53660_Chazo_Juan",
# "E_tricolor_QCAZA53653_Guanujo",
# "E_tricolor_QCAZA53661_Chazo_Juan",
# "E_tricolor_QCAZA53662_Chazo_Juan",
# "E_tricolor_QCAZA53665_Chazo_Juan")
# my_Tree <- drop.tip(my_Tree, DropThese)
# my_Tree <- drop.tip(my_Tree, DropThese_Dups)

```
````
