## Supplementary Data S3 for "Deep Divergences Among Inconspicuously Colored Clades of *Epipedobates* Poison Frogs": specimen_data_2023-06-19.R

```
## set working directory
setwd('/Volumes/GoogleDrive/My\ Drive/Documents//Projects/Epipedobates/Epipedobates_project')

## required libraries
library(ggplot2)
library(cowplot)
library(paletteer) 
library(dplyr)
library(ggpubr)
library(rstatix)

## load data and add new species column
dat<-read.csv("specimen-data-for-SVL_2023-02-05.csv")

dat$New_ID<-factor(dat$New_ID,levels=c("narinensis","spnov","boulengeriSS","aff_espinosai",
                                       "espinosai","anthonyi","machalilla","tricolor"))
dat$Full_ID <- recode_factor(dat$New_ID, narinensis="E. narinensis",
                             spnov="E. sp 1",boulengeriSS="E. boulengeri s.s.",
                             aff_espinosai="E. aff. espinosai",espinosai="E. espinosai",
                             anthonyi="E. anthonyi",machalilla="E. machalilla",
                             tricolor="E. tricolor")
dat$Sex<-as.factor(dat$Sex)

local <-dat %>% group_by(Locality,Full_ID) %>% summarize(ele=mean(na.omit(Elevation))) %>% as.data.frame() 

colors=c("#1DF964FF","orange","#3173BDFF","grey60","#803342FF","#000000FF","aquamarine3","deeppink")
names(colors)<-unique(local$Full_ID)
shapes=c(15,17,5,4)
names(shapes)<-c("M","F","J","ND")
sex_col<-c("black","grey","orange","orange")
names(sex_col)<-c("M","F","J","ND")

## assign shapes for each population of each species
pop_shapes<-c(0,0,1,NA,NA,0,1,2,3,0,NA,1,2,NA,3,NA,5,0,1,2,NA,3,5,0,1,2,0,1,2,0,1,2)

names(pop_shapes)<-local[order(local$Full_ID),]$Locality

# reduce width of boxes in B
byele<-ggplot(dat,aes(Elevation,SVL,shape=Locality,color=Full_ID)) + 
  geom_point(size=2,alpha=0.8,stroke=1) + scale_y_continuous(limits=c(10,25))+
  theme_cowplot() + theme(legend.text=element_text(face="italic"))+
  geom_text(data=local,aes(x=ele,y=25, label=Locality,color=Full_ID),show.legend=FALSE,
            inherit.aes = F,angle=-90) + scale_color_manual(values=colors) +
  scale_shape_manual(values=pop_shapes) + guides(color=guide_legend(title="Species"))

bysex<-ggplot(data=filter(dat,Sex!="ND" & Sex!="J"),aes(Full_ID,SVL,color=Sex,shape=Sex)) + 
  scale_y_continuous(limits=c(10,25))+
  scale_color_manual(values=sex_col) + scale_shape_manual(values=shapes) + 
  geom_point(position=position_dodge(width=0.95),size=3,alpha=0.8) + 
  theme_cowplot() + theme(axis.text.x=element_text(hjust=1,angle = 45,face="italic"),
                          axis.title.x=element_blank()) +
  geom_point(data=filter(dat,Sex=="ND" | Sex=="J"), inherit.aes = F, 
             aes(Full_ID,SVL,shape=Sex,color=Sex),size=2) + guides(color="none",shape="none")+
  geom_boxplot(outlier.shape = NA,position=position_dodge(1),width=0.5,fill=NA) 

top<-plot_grid(bysex,NULL,ncol=2,rel_widths = c(0.7,0.3))
bot<-plot_grid(byele)
plot_grid(bot, top, ncol=1,nrow=2,rel_heights = c(1,1.2))
# the legends were finalized in Adobe Illustrator


#############
## anovas
datA<-subset(dat,Sex=="M" | Sex=="F")
datM<-subset(dat,Sex=="M")
datF<-subset(dat,Sex=="F")
model<-aov(SVL~Sex*Full_ID,data=datA) # interaction term not a significant predictor
model<-aov(SVL~Sex,data=datA) # interaction term not a significant predictor

qqnorm(model$residuals)
qqline(model$residuals)
hist(model$residuals) # left tail; skewed right
shapiro.test(model$residuals) # not normal

# females only
model<-aov(SVL~Full_ID,data=datF)
qqnorm(model$residuals)
qqline(model$residuals) # closer to normal
hist(model$residuals) # left tail; skewed right
shapiro.test(model$residuals) # females normal, males not normal
summary(model)
ttests<-tukey_hsd(model)
subset(ttests,p.adj<0.05)
ttests %>% print(n=Inf)
datA %>% filter(SVL>0) %>%group_by(Full_ID,Sex) %>%summarize(n=length(Sex))

# non-parametric
kruskal.test(SVL~Sex,data=datA) # sig difference
kruskal.test(SVL~Full_ID,data=datM) # sig different
dunn<-dunn_test(datM,SVL~Full_ID)
subset(dunn,p.adj<0.05)
dunn %>% print(n=Inf)

## for boulengeri s.s.
bou<-dat %>% subset(New_ID=="boulengeriSS") %>% na.omit()
model<-t.test(SVL~Locality,data=bou)
summary(model)
model

## linear regression, repeated for each subset of data =='F', =='M', !='J' 
noJ<-subset(dat,Sex=="F") %>% drop_na(Elevation)
model<-lm(noJ$SVL~noJ$Elevation)
qqnorm(model$residuals)
qqline(model$residuals)
shapiro.test(model$residuals)
summary(model)
cor.test(noJ$SVL,noJ$Elevation,method = "spearman",alternative="greater")
plot(noJ$SVL~noJ$Elevation)

# one value per locality
SVLsum<-noJ %>% group_by(Locality) %>% summarize(meanS=mean(SVL),meanE=mean(Elevation))
model<-lm(SVLsum$meanS~SVLsum$meanE)
qqnorm(model$residuals)
qqline(model$residuals)
shapiro.test(model$residuals)
summary(model)
```
